# Identification of microbial markers across populations in early detection of colorectal cancer

**DOI:** 10.1101/2020.08.16.253344

**Authors:** Yuanqi Wu, Na Jiao, Ruixin Zhu, Yida Zhang, Dingfeng Wu, An-Jun Wang, Sa Fang, Liwen Tao, Yichen Li, Sijing Cheng, Xiaosheng He, Ping Lan, Chuan Tian, Ning-Ning Liu, Lixin Zhu

## Abstract

Several studies have investigated the association between microbial and colorectal cancer (CRC). However, the replicable markers for early stage adenoma diagnosis across multiple populations remain elusive. Here, a meta-analysis of six studies, comprising a total of 1057 fecal samples, was performed to identify candidate markers. By adjusting the potential confounders, 11 and 26 markers (*P*<0.05) were identified and separately applied into constructing Random Forest classifier models to discriminate adenoma from control, and adenoma from CRC, achieving robust diagnostic accuracy with AUC = 0.80 and 0.89, respectively. Moreover, these markers demonstrated high diagnostic accuracy in independent validation cohorts. Pooled functional analysis and targeted qRT-PCR based genetic profiles reveal that the altered microbiome triggers different pathways of ADP-heptose and menaquinone biosynthesis (*P*<0.05) in adenoma vs. control and adenoma vs. CRC sequences respectively. The combined analysis of heterogeneous studies confirm adenoma-specific but universal markers across multi-populations, which improves early diagnosis and prompt treatment of CRC.

## Introduction

Colorectal cancer (CRC) is one of the most common cancer with an overall high mortality rate. According to the report of the International Agency for Research on Cancer (IARC), there were over 1,800,000 new CRC cases and over 860,000 deaths in 2018(1). And CRC accounted for approximately 10% of all new cancer cases globally(2). It is estimated that the national expenditures in the United States on cancer care, specifically colorectal cancer, were about 16.63 billion dollars in 2018(3), and the CRC burden is continuously growing over years. Colorectal adenomas are recognized as precursors for the majority of CRC(2). The early detection of CRC at precancerous-stage adenoma has increased the 5-year relative survival rate to about 90%, significantly facilitating early decision making, alleviating the incidence of CRC and reducing economic burden(2, 4).

Gut microbiome is a novel stool-based non-invasive biomarker for metabolic diseases and cancers(5, 6). Many studies have reported that the gut microbiome is an important aetiological element in the initiation and progression of CRC(4, 7) and identified some fecal microbial markers of CRC(8-10). However, there is limited knowledge on whether these biomarkers could more precisely detect early-stage of CRC, adenomas. And this cognitive gap needs to be filled with more intellectual efforts. Furthermore, current knowledge of the associations between microbiome and biomarkers for colorectal adenoma early-detection is poor as well. Only a few studies have investigated the microbial alterations in colorectal adenoma(4, 7, 11-13). However, a substantial variation exists among microbial makers in these studies, and its cause could be various biological factors influencing gut microbiome composition and inconsistent processing of microbial sequencing data.

Meta-analysis offers a set of tools that is powerful, informative and unbiased to improve the robustness of microbiome alterations and reduce the noise of biological and technical confounders so that consistent alterations across multiple studies could be identified. Recently, several meta-analysis of multi-studies have identified universal microbial markers across multiple diseases, such as CRC(11, 13-15), obese(16), Inflammatory bowel disease (IBD)(17), via 16S rRNA sequencing or whole metagenome shotgun sequencing (WMS) technique. However, previous researches based on meta-analysis(11, 13) still could not identify universal stool-based microbial markers for colorectal cancer across multiple cohorts (Supplementary Note 1). Additionally, the commonly used non-invasive stool-based screening test, Faecal Immunochemical Test (FIT), has drawbacks such as poor sensitivity to early and advanced adenoma (7.6% and 38%, respectively)(18). Therefore, it is urgent to explore and identify novel stool-based microbial markers that could more precisely and efficiently diagnose colorectal adenoma and its various stages.

Here, we presented a meta-analysis study, aiming to identify a series of markers that enable distinguishing adenoma from healthy control or CRC with high accuracy across multiple cohorts. We included fecal 16S rRNA sequencing studies considering that 16S rRNA gene-based profiles are more closely matching the “real community”(19). We then investigated the potential mechanisms of the disordered microbiome in colorectal adenomas, which may provide biological insights and therapeutical strategies to detect early syndromes and alleviate symptoms of CRC.

## Results

### Characteristics of the datasets in meta-analysis

In this study, we investigated 16S rRNA sequencing data from four studies to measure the gut microbiome changes as CRC progresses (from control to adenoma to cancer) and to identify the biomarkers specific to adenoma. In total, we collected 307 samples from colorectal adenoma patients, 217 from CRC subjects and 252 samples as control. The demographic information was listed in detail in Table 1. All samples were sequenced at sufficient depth, with average counts of 85637 in each sample. Consistent processing was performed for all raw sequencing data on the QIIME2 platform.

**Table 1.**
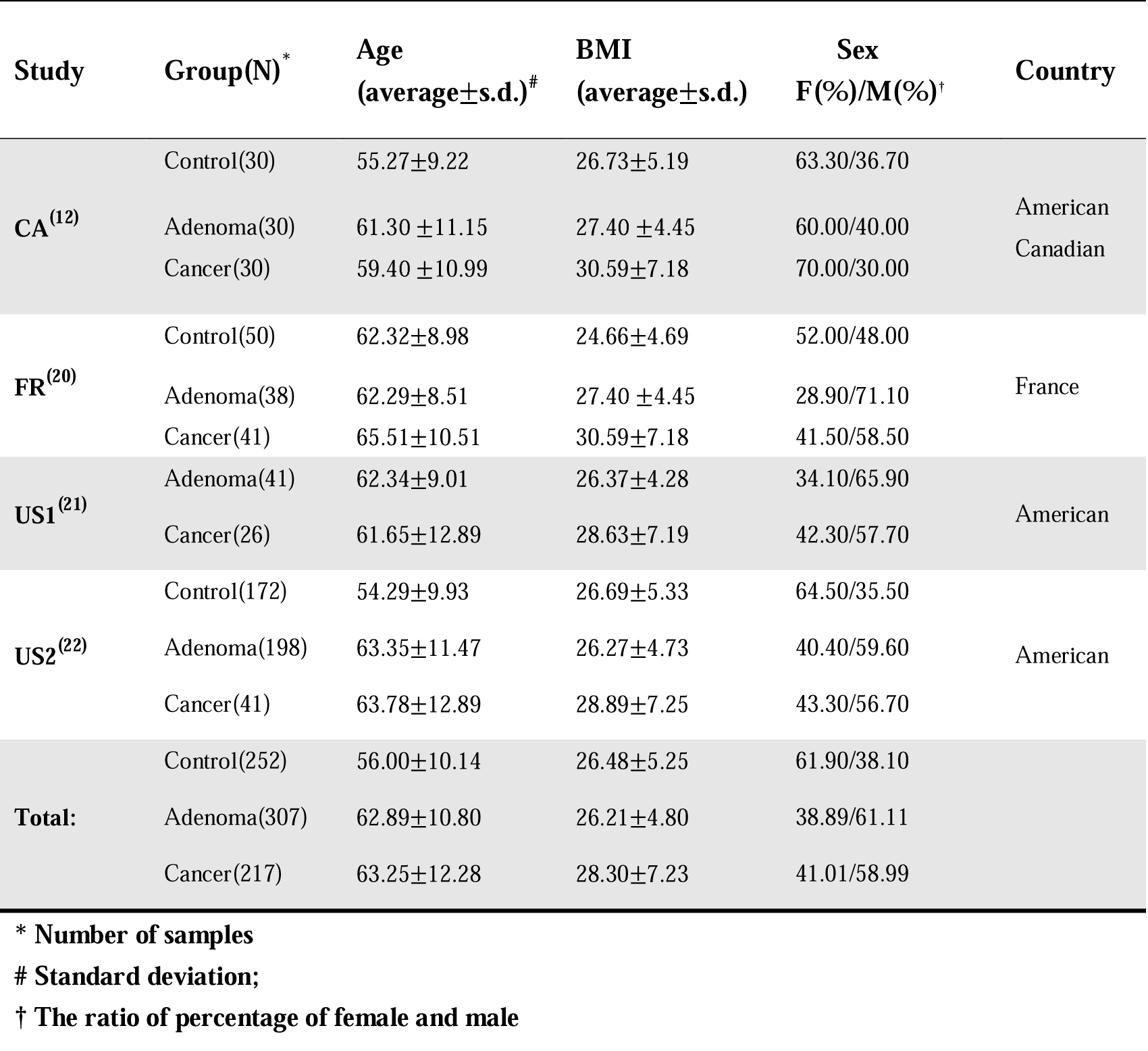
Characteristics of the large-scale adenoma datasets included in this study.

| Study | Group(N)<sup>*</sup> | Age<br>(average±s.d.)<sup>#</sup> | BMI<br>(average±s.d.) | Sex<br>F(%) / M(%)<sup>†</sup> | Country |
| --- | --- | --- | --- | --- | --- |
| CA<sup>(12)</sup> | Control(30) | 55.27±9.22 | 26.73±5.19 | 63.30/36.70 | American<br>Canadian |
|  | Adenoma(30) | 61.30 ± 11.15 | 27.40 ± 4.45 | 60.00/40.00 |  |
|  | Cancer(30) | 59.40 ± 10.99 | 30.59±7.18 | 70.00/30.00 |  |
| FR<sup>(20)</sup> | Control(50) | 62.32±8.98 | 24.66±4.69 | 52.00/48.00 | France |
|  | Adenoma(38) | 62.29±8.51 | 27.40 ± 4.45 | 28.90/71.10 |  |
|  | Cancer(41) | 65.51±10.51 | 30.59±7.18 | 41.50/58.50 |  |
| US1<sup>(21)</sup> | Adenoma(41) | 62.34±9.01 | 26.37±4.28 | 34.10/65.90 | American |
|  | Cancer(26) | 61.65±12.89 | 28.63±7.19 | 42.30/57.70 |  |
| US2<sup>(22)</sup> | Control(172) | 54.29±9.93 | 26.69±5.33 | 64.50/35.50 | American |
|  | Adenoma(198) | 63.35±11.47 | 26.27±4.73 | 40.40/59.60 |  |
|  | Cancer(41) | 63.78±12.89 | 28.89±7.25 | 43.30/56.70 |  |
| Total: | Control(252) | 56.00±10.14 | 26.48±5.25 | 61.90/38.10 |  |
|  | Adenoma(307) | 62.89±10.80 | 26.21±4.80 | 38.89/61.11 |  |
|  | Cancer(217) | 63.25±12.28 | 28.30±7.23 | 41.01/58.99 |  |

\* Number of samples

### Identification of the potential confounder in meta-analysis

Since differences existed among these studies in both technical and biological aspects, we first investigated the potential confounders. The variances explained by disease status for each ASV were calculated to quantify the effects of potential confounders (see method confounder analysis) (Supplementary Fig. 1, 2). This analysis revealed that the factor ‘study’ had a predominant impact on microbial composition (Fig. 1a and Supplementary Fig. 1). Additionally, the microbial alpha and beta diversity also supported that the heterogeneity of studies had a more significant impact on microbial composition than disease status (Fig. 1b and Supplementary Fig. 3). Therefore, we treated ‘study’ as a blocking factor in the subsequent analysis.

**Fig. 1.**
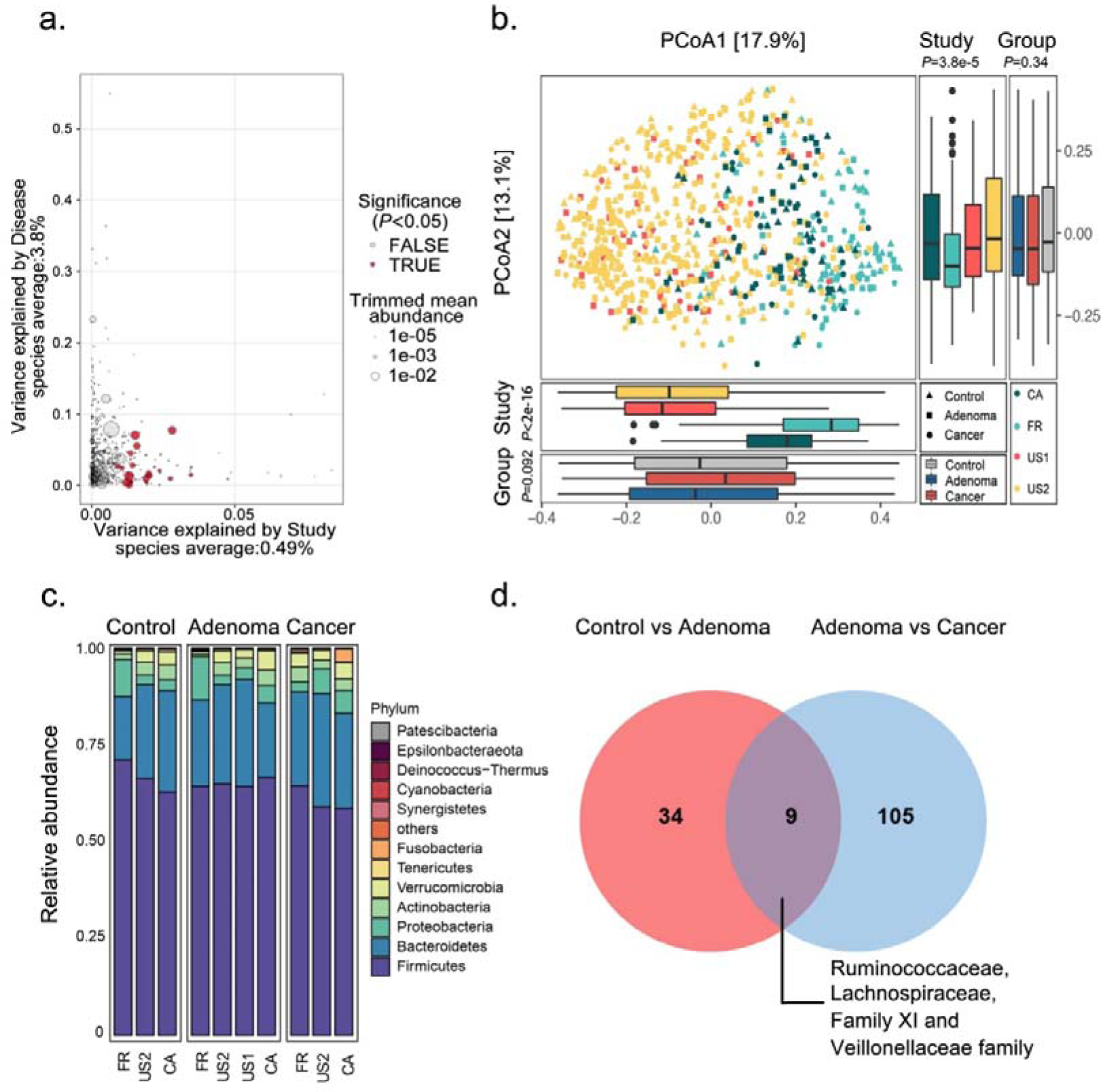
Alterations of gut microbial composition in different disease status accounting for study heterogeneity. a, Variance explained by disease status (adenoma versus cancer) is plotted against variance explained by study effects for individual ASVs. The significantly differential ASVs are colored in red and the dot size is proportional to the abundance of each ASV. b, Principal coordinate analysis of samples from all four studies based on Bray-Curtis distance; the study is color-coded and the group (control, adenoma and cancer) is indicated by different shapes. The upper-right and the bottom-left boxplots illustrate that samples projected onto the first two principal coordinates broken down by study and disease status, respectively. *P* values were calculated with a Kruskal–Wallis test for study and group. All boxplots represent 25th–75th percentile of the distribution; the median is shown in thick line at the middle of the box; the whiskers extend up to values within 1.5 times, and outliers are represented as dots. c, Relative proportions of bacterial phyla in healthy controls, adenomas and CRC across four different studies. d, Venn diagram shows the overlap of differential ASVs between adenomas and healthy controls or CRC.

**Fig. 2.**
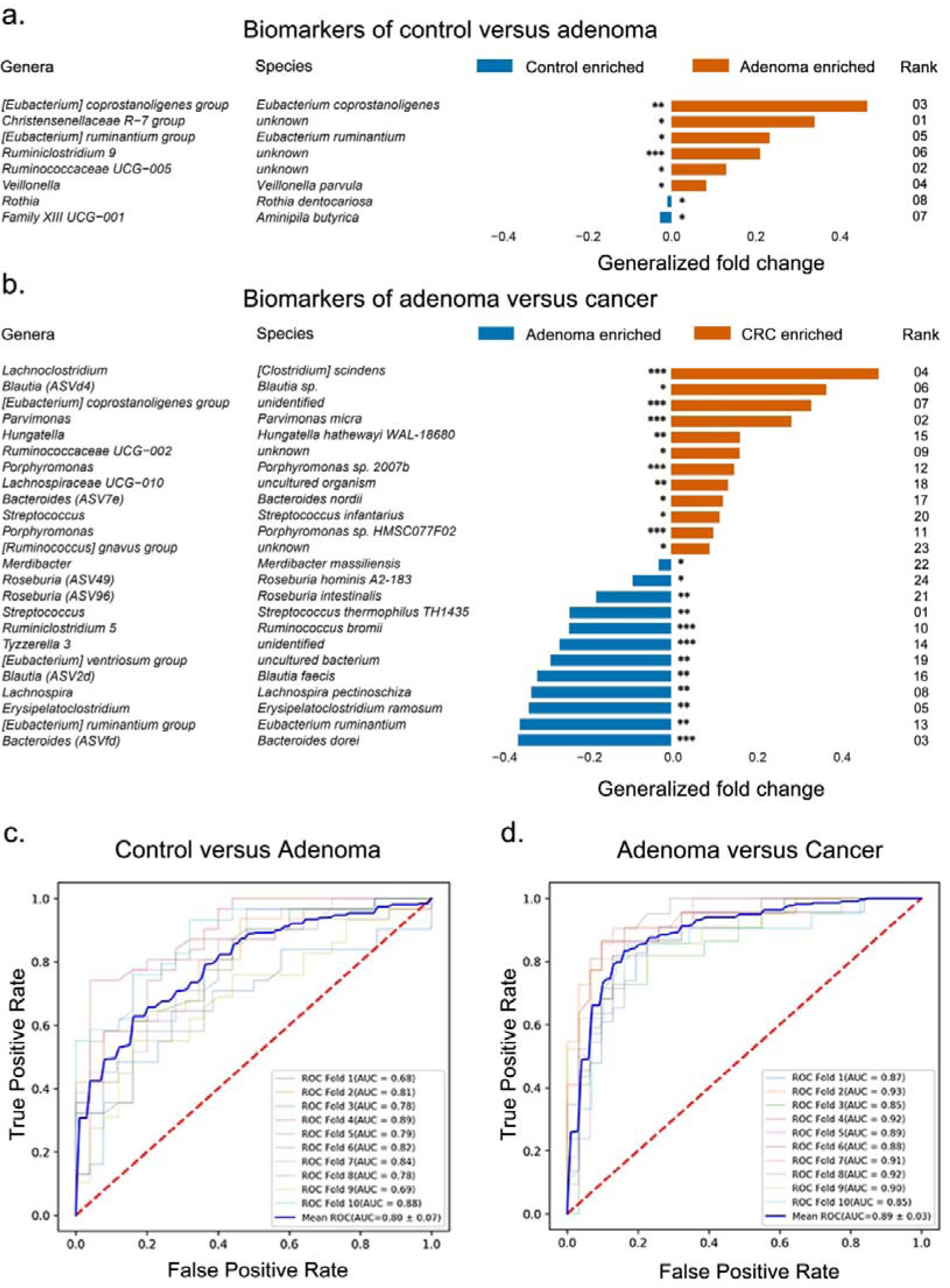
Performance of discriminating adenoma from control or cancer using important features. a, b, The important biomarkers identified to construct RF model for discriminating adenoma from control (a) and CRC (b). The rank in (a) and (b) means the order of feature importance in the RF model; *: *P* < 0.05, **: *P* < 0.01and **: *P* < 0.001. c, d, The AUC of the optimized models constructed with biomarkers and metadata of Control versus Adenoma (c) and Adenoma versus Cancer (d). Mean ROC in (c) and (d): the average AUC from tenfold cross-validation.

**Fig. 3.**
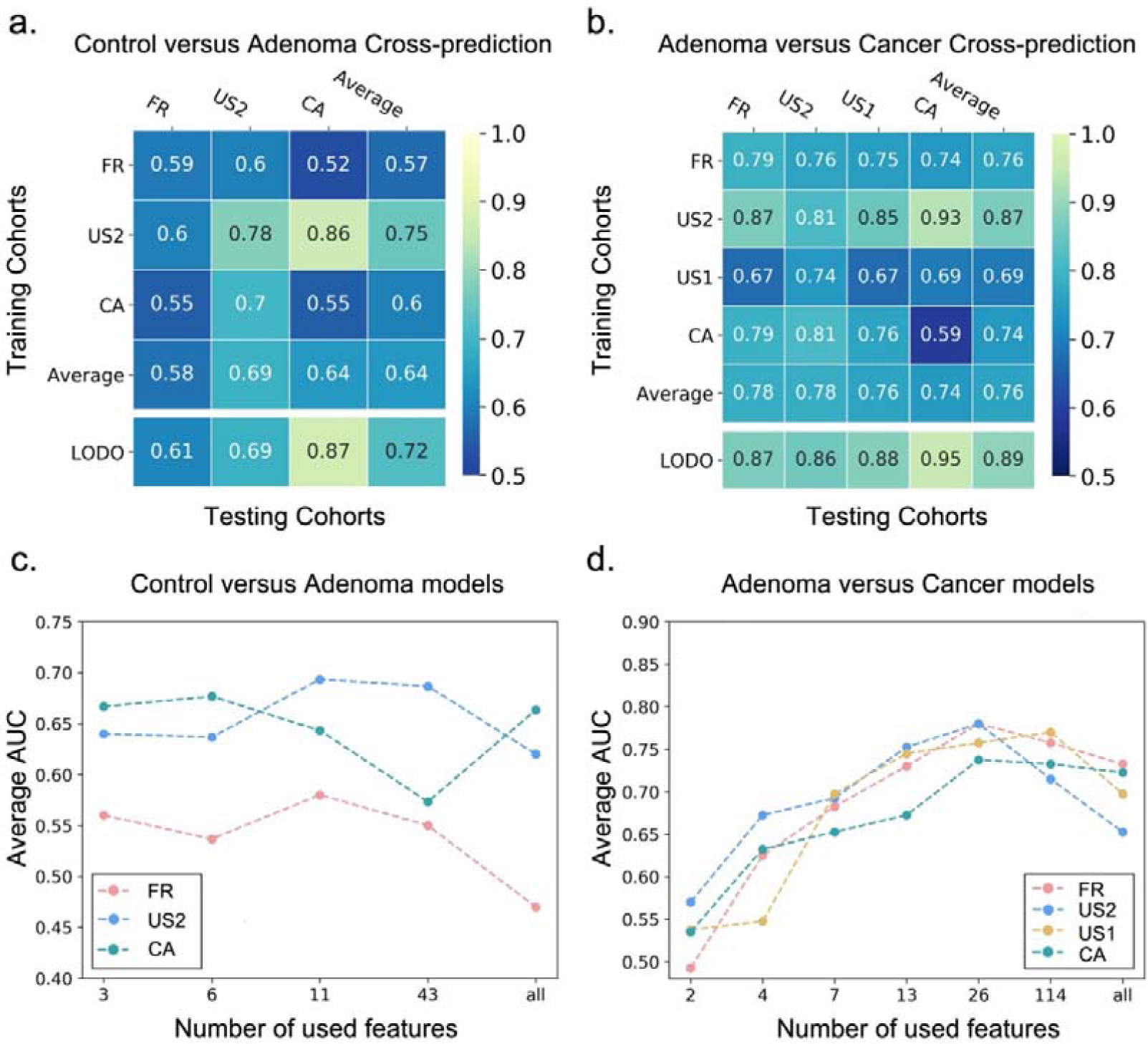
Prediction performance of important features across studies and identification of minimal features for detecting adenoma. a, b, Cross-prediction matrix depicting prediction values for differentiating adenoma from control (a) and CRC (b) as AUC obtained using important features. Values on the diagonal refer to the results of within-cohort validation; Off-diagonal values refer to the AUC values obtained from cross-cohort validation, which training the classifier on the study of the corresponding row and applying it to the study of the corresponding column; The LODO values refer to the performances obtained by training the classifier using all but the study of the corresponding column and applying it to the study of the corresponding column (see methods). c, d, Average AUC of study-to-study transfer validation classifiers for control versus adenoma (c) and adenoma versus cancer (d) at different sets of features. The x-axis in (c) and (d) indicate different sets of features: All (c-d): all ASVs; 43 (c) and 114 (d): differentially abundant ASVs; 11 (c) and 26 (d): all important features and other top-ranking important features. The different studies were indicated in different colors.

### Alterations of gut microbial composition in colorectal adenoma

At the phylum level, the gut microbiota was dominated by members of Firmicutes and Bacteroidetes, followed by Proteobacteria, Actinobacteria, Verrucomicrobia, Tenericutes and Fusobacteria in healthy controls, adenomas and CRC. These dominant phyla were similar to those reported in previous studies on gut microbiota(20). Furthermore, the phylum Fusobacteria, the most CRC-associated bacteria as reported(23), were observed with significantly decreased abundance in adenoma compared to that in cancer, while there was no significant difference between adenoma patients and controls (Fig. 1c).

At the ASV level, significant alterations across studies were observed among different disease status. In the comparison of gut communities between controls and patients with adenoma, 43 ASVs were identified with distinguishable abundances (Supplementary Note 2). Moreover, we also identified 114 differentially abundant ASVs between adenoma and cancer (Supplementary Note 3).

Additionally, pathogenic bacteria with increased abundance were detected in adenoma or cancer compared with control. For instance, *Parvimonas* genus was enriched in adenoma compared with controls while *Fusobacterium, Porphyromonas, Peptostreptococcus, Parvimonas*, and *Escherichia-Shigella* genus were enriched in cancer compared with adenoma. Particularly, *Fusobacterium, Porphyromonas, Parvimonas* and *Peptostreptococcus* were identified as oral pathogens associated with CRC(17, 24). Notably, there were only 9 common differential ASVs between healthy controls versus adenoma and adenoma versus cancer, which could be further classified into Ruminococcaceae, Lachnospiraceae, Family XI and Veillonellaceae family (Fig. 1d). The two sets of differential ASVs with a Jaccard distance of 0.939 indicate that the microbiota has a remarkable difference between adenoma and control or cancer.

### Microbial classification models for colorectal adenoma

Next, we constructed RF models by pooling all samples to select features capable of distinguishing adenoma from control and cancer. Besides using differential ASVs as key metrics, alpha diversity indices including Shannon Index, Simpson Index and Observed ASVs, and three patient metadata, age, gender and BMI were also included in model building. To obtain the best performing models and most important features, an IEF step was further applied.

A robust RF model was constructed with a core set of important features, including 8 differential ASVs (as biomarkers) together with age, gender and BMI, which were proved to have the best capability to distinguish control subjects from patients with adenoma (AUC = 0.80) (Fig. 2a, c and Supplementary Table 3). Among these, the ASV assigned as *Christensenellaceae R-7 group sp*. was the highest-ranking biomarker (Fig. 2a). The biomarkers also included the increased abundance ASVs of *[Eubacterium] coprostanoligenes group, Ruminiclostridium 9 sp*., *Christensenellaceae R-7 group sp*., *Ruminococcaceae UCG-005 sp*. and *Veillonella parvula* as well as the decreased abundance of *Rothia dentocariosa* and *Aminipila butyrica* in adenoma (Supplementary Fig. 4).

**Fig. 4.**
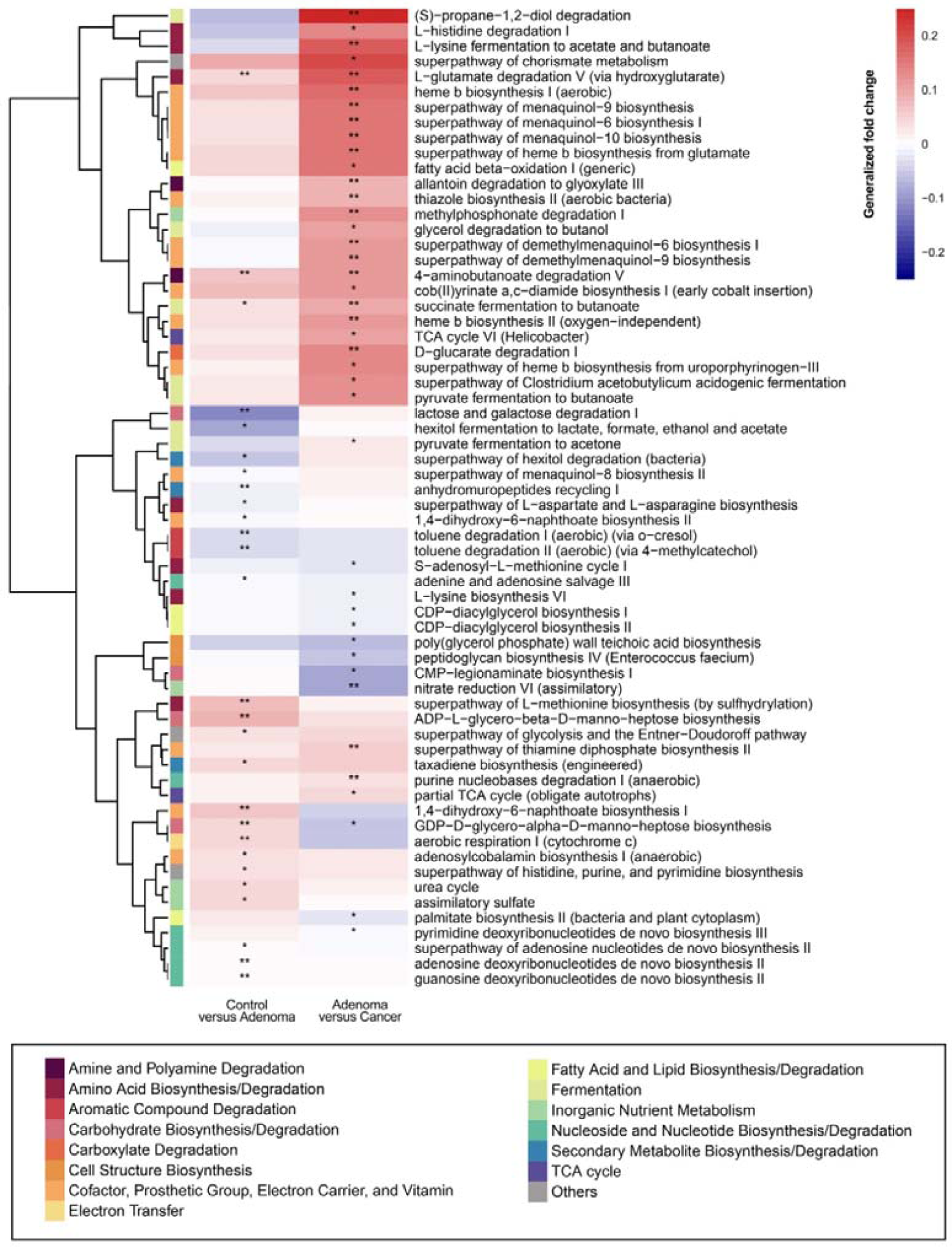
Functional alterations in control, adenoma and cancer. The relative abundances of functional pathways were compared between adenoma and control or cancer. These pathways were differentially abundant ones, with *: *P* < 0.05, **: *P* < 0.01. Generalized fold change (see method) was indicated by a gradient color. The generalized fold change >0: enriched in later; generalized fold change <0: enriched in former.

Similarly, the best performance of the RF model for distinguishing adenoma from cancer is 0.89 (AUC). The RF model was built with 24 ASVs together with age and BMI (Fig. 2b, d and Supplementary Table 4). Among these biomarkers, the ASV belonging to *Streptococcus thermophilus TH1435* was the top-ranking biomarker(Fig. 2b). The following ASVs were assigned as *Parvimonas micra, Bacteroides dorei, [Clostridium] scindens, Erysipelatoclostridium ramosum, Blautia sp*., *[Eubacterium] coprostanoligenes group sp*. and *Lachnospira pectinoschiza* (Fig. 2b). The *[Clostridium] scindens* was significantly (*P* < 0.001) enriched in cancer compared with adenoma with a generalized fold change of 0.49. Additionally, the abundance of *Blautia sp*., *Hungatella hathewayi WAL-18680* and *Eubacterium ruminantium* were gradually increased while *Streptococcus thermophilus TH1435, Erysipelatoclostridium ramosum, [Eubacterium] ventriosum group sp*. and *Roseburia intestinalis* were gradually decreased during CRC carcinogenesis (Supplementary Fig. 5). In the two models, age was ranked as the top and third predictor in the testing phase, respectively. In the two sets of biomarkers, there was only one common ASV classified as *Eubacterium ruminantium*.

**Fig. 5.**
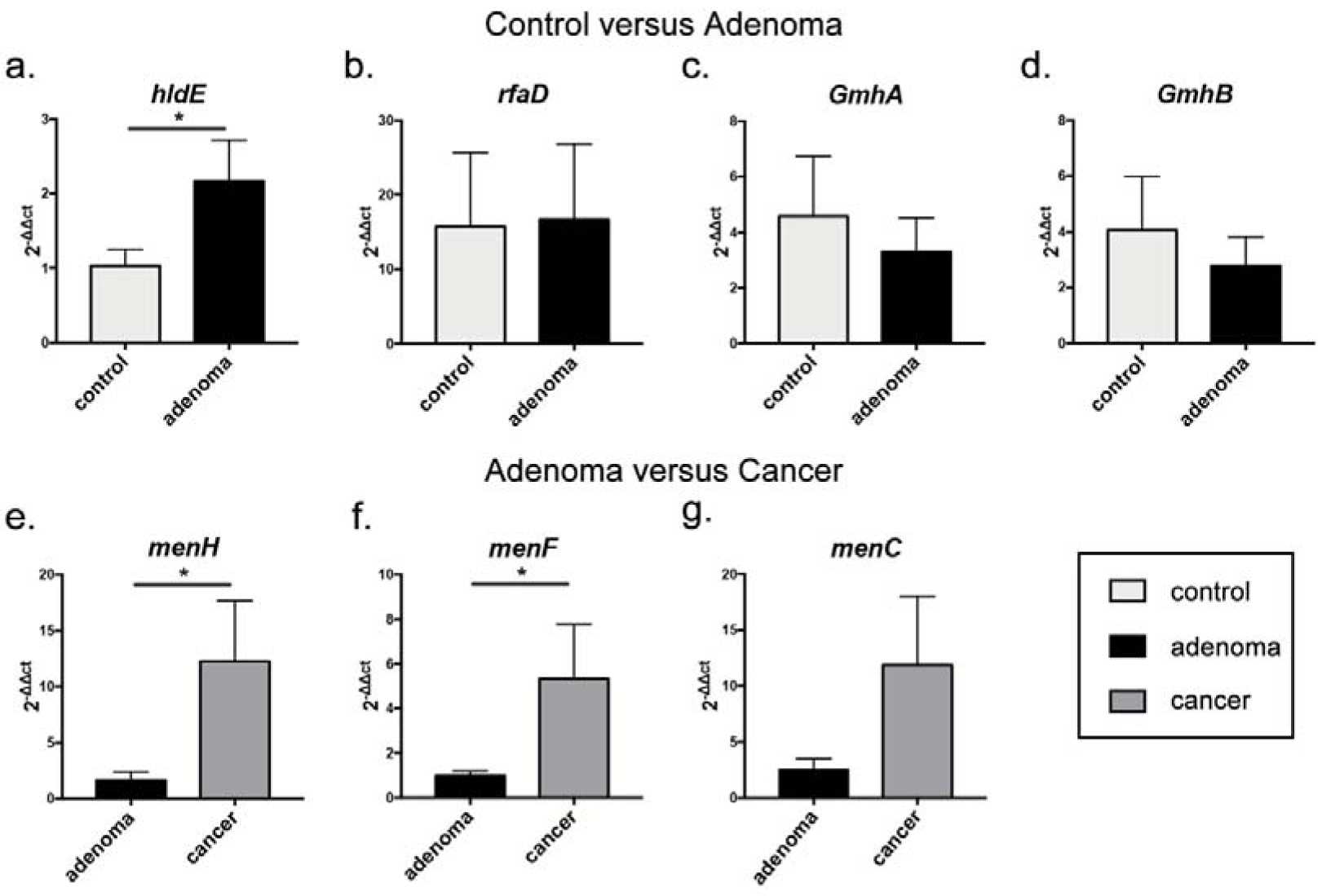
Relative abundance of candidate genes is plotted against qRT-PCR quantification in gDNA extracted from stool samples of healthy controls, adenomas and CRC. The expression of (a) *hldE*; (b) *rfaD*; (c) *GmhA*; (d) *GmhB* were compared between control and adenoma groups, while the expression of (e) *menH*; (f) *menF*; (g) *menC* were compared between between adenoma and cancer groups. All results are presented as mean ± standard error. *: *P* < 0.05.

Moreover, we also identified that a core set of 34 ASVs, together with age, gender and BMI, collectively had the highest capability to distinguish control from cancer (AUC=0.93) (Supplementary Fig. 6, Supplementary Note 4). It is worth noting that there was no common ASV in the two sets of biomarkers between healthy controls and adenomas or CRC(Supplementary Fig. 7). Thus these results highlighted that microbial markers aimed to detect CRC are specific and exclusive, and would not be used as optimal diagnosis of adenoma.

### Co-occurrence and clustering analysis of microbiota in different states

Through the co-occurrence network of differential ASVs, our results suggested that most of the identified biomarkers have functional importance in the network (Supplementary Note 5). To gain further insight, we analyzed metagenomes of patients in adenoma and control. Co-occurrences analysis demonstrated four clusters of biomarkers with distinct taxonomic composition (Supplementary Fig. 9a). These clusters are not tightly associated with patient characteristics such as Age, Sex and BMI (Supplementary Fig. 10a), revealing that the adenoma-associated microbiota closely resembles that of the healthy control. These results further proved the high detection accuracy (AUC of 0.8) and overall success of merely using 11 important features to distinguish control from adenoma.

Moreover, we also explored the CRC patient metagenomes for co-occurrences among a panel of 24 biomarkers and yielded three clusters (Supplementary Fig. 9b). Cluster 2 demonstrated strong taxonomic consistency, which was primarily comprised of members following Clostridiales order. In contrast, the other two clusters exhibited heterogeneous taxonomy, with cluster 1 containing high-ranking biomarkers and cluster 3 assorting together the species that highly prevailed in CRC individuals. We then investigated the association between these three clusters and various tumor characteristics. Clostridiales cluster 2 is significantly enriched in male CRC patients. Besides, both cluster 1 and cluster 3 show a slight tendency toward late-stage CRC (containing stages 3 and 4 according to the American Joint Committee on Cancer), and this tendency is significant for cluster 3. Associations with patient age and BMI are weaker and not significant (Supplementary Fig. 10b). Based on these results, it can be deduced that the adenoma-associated microbiota differs from that of CRC. To consider the impact of using different studies, all of these tests were adjusted by blocking for “study” (see method co-occurrence and clustering analysis).

### Validation of the colorectal adenoma classifiers

To test whether the selected important features are universal and robust across multiple studies, we performed study-to-study transfer validation and LODO validation on the entire samples. In study-to-study transfer validation, the average AUC for healthy controls versus adenoma model was 0.64 and the AUCs of all datasets ranged from 0.52 and 0.86, which maintained the diagnostic accuracy of within-study (Fig. 3a). Notably, the US2 study serves as a better training set than other studies through exhibiting relatively higher testing AUCs. The reason could be that the US2 study has a larger data set that is beneficial to develop accurate classifiers. Moreover, we also compare the diagnostic performance of selected important features with FIT, which illustrates improved adenoma diagnostic ability by combining with non-invasive clinical screening tests (Supplementary Note 6). Additionally, in the LODO analysis, the AUC values of control versus adenoma models range from 0.61 to 0.87 with an average 0.72, which is superior to study-to-study transfer validation owing to using large datasets (Fig. 3a). This reveals that more training samples would in principle improve the robustness of classifiers.

Similar results were observed in the adenoma versus cancer model (Fig. 3b). The average AUC of study-to-study transfer validation is 0.76 and the AUCs of all datasets range from 0.59 to 0.93. Besides, the classification accuracy of the CA within-study is relatively low. The possible reason is that the CA study was small-sized and subjects came from different countries, which indicates that classifiers may be geographic region-specific when dataset is limited. This result reinforced Zhou’s research that region variation limits the usefulness of disease modelling(25). Moreover, the AUC values are also elevated in the LODO analysis, ranging from 0.86 to 0.95 with an average 0.89 (Fig. 3b). We notice that the classifiers performed better in adenoma versus cancer than that in control versus adenoma, which reinforced previous findings that the adenoma-associated stool microbiome closely resembled that of the health status(7, 11, 20).

To determine the maximum subset of important features required to provide comparable accuracy on validation studies and methods, we analyzed sets of features including all ASVs (all), differentially abundant ASVs (control versus adenoma:43, adenoma versus cancer:114), all important features (control versus adenoma:11, adenoma versus cancer:26) and reduced important features according to the feature ranks of the RF classifiers. In both study-to-study transfer validation (Fig. 3c, d) and LODO validation (Supplementary Fig. 12a, b), as the number of important features increases, the average AUC increases and reaches maximum when all the important features are included for all studies except the CA study. This may also be owing to the characteristics of small-sized and geographic heterogeneity in the CA study. As we continued to add more ASVs, especially the ones not part of important features among disease status, the average AUC of cross-validation decreases. Therefore, this result further confirmed that the sets of selected important features contributed to the accuracy of classifiers.

### Validation of colorectal adenoma markers in independent cohorts

To further validate our meta-analysis results, two additional independent cohorts from America (Validation Cohort1) and China (Validation Cohort2) were incorporated into this study. The validation cohort1 is comprised of 70 controls and 102 adenoma patients, while there were 57 adenoma patients and 52 CRC patients in the validation cohort2(Supplementary Table 8). The independent predictive RF model was confirmed to be relatively accurate on the two new cohorts, with an AUC of 0.73 and 0.83 for distinguishing adenoma from controls or cancer, respectively (Supplementary Fig. 13a, b). Although the validation cohort2 was lack of patient metadata, it obtained a relatively high AUC, indicating that the gut microbial biomarkers could distinguish adenomas from CRC precisely. Additionally, *Ruminococcaceae UCG-005 sp*. and *Christensenellaceae R-7 group sp*. were confirmed as the top-ranking biomarkers between controls and adenoma patients in validation cohort1. Furthermore, *Parvimonas micra, Streptococcus thermophilus TH1435* and *Bacteroides dorei* were confirmed as the three top-ranking biomarkers for distinguishing between adenoma and CRC patients in validation cohort2.

### The specificity of colorectal adenoma predictive models

After evaluating the accuracy of the above colorectal adenoma predictive models on different cohorts, we further validated the specificity of colorectal adenoma related important features in other potentially microbiome-linked diseases. Five microbiome-linked diseases including NAFLD, T2D, CD, UC and IBS were considered in this analysis(Supplementary Table 8). We randomly drew samples from each disease and the control of these non-CRC studies and added them to the control class of the validation cohort1. By comparing AUC scores between adding non-CRC cases and adding the corresponded external controls, we found a small decrease (ranging from 1% to 4%) in prediction accuracy for the non-CRC group(Supplementary Fig. 14). When adding both control and case samples of the IBD study, the AUCs decreased more than other studies, which may be caused by lack of several key features including BMI and two biomarkers. Taken together, these results indicated that the colorectal adenoma-specific model might not be necessarily applicable to other microbiome-associated diseases.

### Microbial functional changes in colorectal adenoma

We investigated the microbial-based functional alterations for multiple different disease status. There are 27 differential pathways between control and adenoma (Supplementary Table 9) and 41 differential pathways between adenoma and cancer (Supplementary Table 10) consistently detected across studies. A total of 64 differential pathways (4 pathways were overlapped) were clustered based on their generalized fold change scores. (Fig. 4, Supplementary Note 7).

Notably, the biosynthesis of ADP-heptose, a key metabolic intermediate in the biosynthesis of lipopolysaccharide (LPS) was significantly enriched in adenoma compared with control. It was associated with the activation of Nuclear factor-κB (NF-κB) and a strong pro-inflammatory response(26) which led to colorectal adenoma. The ASV was assigned as *Veillonella*, one of the biomarkers differentiating healthy controls from adenoma samples (Fig. 2a). It was highly ranked among all ASVs in the average contribution of the ADP-heptose. (Supplementary Table 11). There are five limiting steps catalyzed by genes of *hldE, rfaD, gmhA* and *gmhB* in the biosynthesis of ADP-heptose. These four genes were consistently enriched in adenoma compared with control (Supplementary Table 12). To further validate the results and explore its possibility of application in diagnosis, we analyzed the expression patterns of these genes based on qRT-PCR. As shown in Fig. 5a-d, the expressions of the *hldE* and *rfaD* gene were enriched in adenoma compared with control, in consistent with the picrust2 results, especially that the *hldE* gene was statistically significant.

Moreover, it is worth noting that menaquinone (Vitamin K2) biosynthesis was significantly enriched in cancer compared with adenoma. Especially, the MK-10 (one type of Vitamin K2) was mainly produced by *Bacteroides*, one of the biomarkers between adenoma and cancer (Fig. 2b), which reinforced the previous study that *Bacteroides* had high-level production of MK-10(27). The ASV assigned as *Bacteroides* ranked the 3^rd^ and 4^th^ in contribution to MK-10 biosynthesis in adenoma and cancer among all ASVs (Supplementary Table 13). Collectively, the production of Vitamin K2 by microbiota may serve as a response to compensate for induction of feedback inhibition in colorectal cancer cells(28). We found a significantly increased abundance of *menH, menF* and *menC* in CRC samples compared with that of control in pooled datasets by blocked Wilcoxon test (Supplementary Table 12). These results were further confirmed in adenoma and CRC by qRT-PCR on several patient samples (Fig. 5e-g), especially the *menH* and *menF* genes with statistical significances.

## Discussion

This study comprehensively assessed the alterations of CRC-associated gut microbiome and the ability of microbial markers for the early detection of CRC. Thus, we first constructed machine learning classifiers. The best performing model achieves a high accuracy (AUC=0.80) with 11 important features to distinguish colorectal adenoma from non-tumor controls (Fig. 2c). Similarly, the AUC of the best model for detecting colorectal adenoma from CRC with 26 important features is 0.89 (Fig. 2d). Through study-to-study transfer validation and LODO validation across multiple datasets, the selected microbial markers could overcome technical and geographical discrepancies with the average AUC of 0.72 in the adenoma-control model (Fig. 3a) and 0.89 in the adenoma-cancer model (Fig. 3b), while previous researches revealed that the majority of differential microbial taxa differed in given case-control studies(17). Furthermore, the two additional independent cohorts strengthened and validated the extensibility of these makers (Supplementary Fig. 13a, b). The accuracy of classifiers with adenoma-specific markers is higher than that in previous WMS based studies(11, 14), probably due to more complete taxonomic profilings represented by ASVs. WMS data is well-recognized to possess the advantage of species- and even strain-level resolution. However, the current strategies for characterizing microbial community compositions with WMS are “closed annotation” that strongly rely on the known reference genome database(29-31), which is likely missing some species without known genomes or maker genes. It will thus result in biases in relative abundance estimation. Consistently, we built the cancer-control model with a panel of 34 important ASVs and achieved performance AUC of 0.93, whose accuracy is significantly higher than meta-analysis based WMS analysis(AUC=0.84)(11, 14). Importantly, the co-occurrence network analysis confirmed several biomarkers that are crucial in subnetworks, for example, *Ruminococcaceae UCG-005 sp*., *[Clostridium] scindens, Blautia sp*., etc. Furthermore, the performance of these features in diagnosing colorectal adenoma worsened when randomly adding samples from other microbiome-associated diseases, such as IBD and NAFLD(Supplementary Fig. 14), indicating that the panel of markers was adenoma-specific. Overall, all these validations point to the robustness of the classifiers and provide evidence that microbial classifier could serve as an effective non-invasive clinical indicator for colorectal adenoma.

Several other studies have reported that some fecal bacteria could serve as biomarkers for non-invasive diagnosis of colorectal cancer, such as *Fusobacterium nucleatum, Escherichia coli, Bacteroides fragilis*(8, 32-34). Unlike these existing studies, we aim to identify microbial-derived markers that could effectively diagnose adenoma (early stage of colorectal cancer), which represents a primary target for CRC screening at the early stage, as majority of CRC begins with the malignant transformation of benign polyps, the colorectal adenoma(2). Microbial communities altered in both colorectal adenoma and cancer during the progression of CRC, and the difference of microbiome alteration remains unclear. Notably, we found markers for distinguishing adenoma and cancers from healthy controls are not always the same (Supplementary Note 8). What’s more, the combination of the important adenoma-specific features and FIT improved the classifier’s accuracy (AUC=0.81) compared to microbial makers (AUC=0.78) or FIT (AUC=0.60) alone (Supplementary Fig. 11), indicating the non-invasive clinical screening tests could be used as complementary characteristics of gut microbiota for early screening of adenoma. Recently, a 16s rRNA analysis showed that microbiome dysbiosis in adjacent tissues could discriminate colorectal adenomas from healthy controls effectively(13), providing a new insight for following research of adenoma biomarkers.

The functional analysis sheds light on the convoluted underlying mechanisms and would greatly enhance our understanding and interpretation of CRC development (Supplementary Fig. 15). Among the differential pathways, we found the biosynthesis of ADP-heptose is significantly enriched in adenoma compared with control. ADP-heptose is a key metabolic intermediate in the biosynthesis of LPS, which is associated with the activation of NF-κB and then induces a strong pro-inflammatory response(35) in the initiation and progression of colorectal cancer, especially in adenoma(36). More importantly, the contribution decomposition analysis indicated that the adenoma-specific marker Veillonella parvula was highly ranked in the average contribution of the ADP-heptose among all ASVs (Supplementary Table 11). This suggests that the microbial markers contributed to the activation of pro-inflammatory pathways that ultimately led to the progression of colorectal adenoma. Notably, *hldE* was an important bifunctional protein involved in the biosynthesis of ADP-heptose, which catalyzes the nucleotide-activated heptose precursors used in the biosynthesis of LPS and in post-translational protein glycosylation(37). *HldE* was significantly enriched in adenoma compared with control according to computational finding and was further validated by qRT-PCR validation analysis(Fig. 5a). Since the *hldE* was reported to play an important role in bacterial virulence(37), it is promising to utilize it as an attractive target for therapeutic treatment of colorectal adenoma. Moreover, a series of Vitamin K2 biosynthesis is significantly different between adenoma and cancer. Especially, the MK-10 pathway increased in cancer compared with adenoma and the biomarker Bacteroides dorei was ranked as the third and fourth contributor to MK-10 biosynthesis in adenoma and cancer among all ASVs (Supplementary Table 13). Computational finding and qRT-PCR results demonstrated that the *menH* and *menF* gene were significantly increased in CRC compared with adenoma (Fig. 5e, f). Specifically, *menH* catalyzes the first step and plays distinct roles in the biosynthesis of MK-10 biosynthesis(38). The increased abundance of *menH* in CRC samples activated the synthesis of this pathway. Previous studies indicated that Vitamin K2 played a key role in antitumor effect via cell-cycle arrest, cell differentiation and cell apoptosis(28). Therefore, the increased production of Vitamin K2 may be a compensatory effect of the dysregulated microbiota to survive the tumor microenvironment, which also shows a potential novel CRC intervention strategy targeting Vitamin K2 biosynthesis bacteria. Though the main pathways differ between the control-adenoma and the adenoma-CRC stage, all these important pathways triggered by altered microbiome could offer promising perspectives and evidence for intervention and treatment in the CRC carcinogenesis (Supplementary Note 9).

## Methods

### Data collection

We collected data from studies in PubMed.gov that published 16S rRNA sequencing data on patients with CRC, adenomas and healthy controls. Only four studies with accessible metadata of every sample were included in this work. Raw sequencing data of these studies were downloaded from Sequence Read Archive (SRA) and European Nucleotide Archive (ENA) using identifiers: PRJNA389927 for Zeckular et al(12), PRJEB6070 for Zeller et al(20), PRJNA290926 for Baxter et al(22) and PRJNA362366 for Sze et al(21). Besides, two additional cohorts (Supplementary Table 8) were used as independent cohorts with accession numbers PRJNA534511(39) and PRJNA280026(40).

The collection of human data for real-time quantitative PCR (qRT-PCR) analysis was approved by the Review Board of School of Public Health, Shanghai Jiao Tong University School of Medicine. Patients were recruited for initial diagnosis and had never received any treatment before fecal sample collection. Patients with hereditary CRC syndromes, with a previous history of CRC were excluded from the study. Based on pathological section and colonoscopy results, recruited subjects were classified into three groups: (1) healthy subjects, namely controls: individuals with colonoscopy negative for tumor, adenoma or other diseases; (2) patients with adenoma: individuals with colorectal adenoma(s); and (3) patients with CRC: individuals with newly diagnosed CRC. A total of 94□subjects were initially recruited based on criteria sex, age, BMI and other confounding factors. Finally, 43 were remained: 30□patients with CRC, 6□adenomas and 7□controls. Stool was collected in fecal collection tubes and was stored at –80□°C. DNA was extracted from fecal samples using Stool Genomic DNA Kit (CW20925, CWBIO, China) following the manufacturer’s instructions. Relative gene expression by qRT-PCR and the patient characteristics for qRT-PCR were summarized in Supplementary Table 14.

### Data Preprocessing

The 16S rRNA sequencing data were analyzed using Quantitative Insights Into Microbial Ecology (QIIME2 V.2018.11), a plugin-based platform for microbiome analysis(41). DADA2 software, wrapped in QIIME2, was used to filter out sequencing reads with quality score Q > 25 and denoise reads into amplicon sequence variants (ASVs) (i.e. 100% exact sequence match), resulting with feature tables and representative sequences. Taxonomy classification was assigned based on the naϊve Bayes classifier using the classify-sklearn package(42) against the Silva-132-99 reference sequences. ASVs that couldn’t be precisely annotated to species were reassigned to ones having the most similar sequences in the same genus (or family) using NCBI Blast. Subsequently, representative sequences were aligned using Fast Fourier Transform (MAFFT) in Multiple Alignment and a phylogenetic tree was generated with the Fast-Tree plugin. Then, the feature tables were converted to relative abundance tables. A set of ASVs that were confidently detectable in at least three studies and were present in at least 80% of samples were selected for further analysis.

### Confounder analysis

We used ANOVA-like analysis(14) to quantify the effect of potential confounding factors and disease status. The total variance of a given ASV was compared to the variance explained by disease status (control, adenoma and cancer) and the variance by confounding factors (age, BMI, diabetes, Nonsteroidal anti-inflammatory drug (NSAID), platform, race, gender and study) akin to a linear model. Variance calculations were performed on ranks to account for non-Gaussian distribution of microbiome abundance data(14). Potential confounding factors with continuous values were transformed into discrete variables either as quartiles or in the case of BMI as groups of lean(>25), overweight (25-30), and obese(>30) based on conventional cutoffs.

### Meta-analysis of differentially abundant ASVs

The significance of differential abundance was tested on a per ASV using the blocked Wilcoxon test implemented in the R (V.3.5.2) ‘coin’ package (*P* values < 0.05 were deemed as significant in all differential analysis). Confounder with high variance explanation was defined as a block to adjust the differential analysis. Significance was tested against a conditional null distribution derived from permutations of the observed data. Permutations were performed within ‘study’ to control variations in block size and composition(14). For further analysis, we evaluated a generalization of the (logarithmic) fold change for each ASV. This quantity is widely applied to genomic sequencing data such as RNA-seq and GRO-seq and further improved for better resolution of sparse microbiome profiles(43). The generalized fold change was calculated as the mean difference between predefined quantiles (ranging from 0.1 to 0.9 in increments of 0.1 in this study) of the logarithmic control and adenoma, and between adenoma and cancer distributions.

### Model construction and features extraction

Following the differentially abundant ASVs analysis, we built Random Forest (RF) classifier models with stratified 10-fold cross-validation to distinguish adenoma from cancer or control. The features used for model building consisted of patient metadata as well as differential ASVs and alpha diversity indices. The alpha diversity indices consisted of Shannon Index, Simpson Index and Observed ASVs, while the patient metadata features consisted of age, gender and BMI. The RF classifier models were built with 501 estimator trees and each tree had 10% of the total features. Then an iterative feature elimination (IFE) step was used to optimize the performance of subsequent RF models. The top features from the top-performing model were selected as “important features” and the top microbial features as “biomarkers”. Finally, the AUC was used to evaluate the performance of the optimized models.

### Co-occurrence and clustering analysis

To further analyze the co-occurrence of biomarkers, the relative abundances of biomarkers were discretized into binary values ‘positive’ or ‘negative. For each biomarker, the 90th percentile in control or adenoma was used as the threshold. A sample was labeled ‘positive’ when the relative abundance of ASV was above the defined threshold(14). Based on the binarized markers-by-sample matrix, biomarkers were then clustered using the Jaccard index. Associations between clusters and metadata were calculated by a Cochran–Mantel–Haenszel test with ‘study’ as blocking factors.

### Model evaluation

To assess the generalizability of microbial-based adenoma classifiers across geographic and technical differences of metagenomic data generation and processing in multiple patient populations, both study-to-study transfer validation and leave-one-dataset-out (LODO) validation were performed. In study-to-study transfer validation, RF classifiers were trained in one single study and externally assessed on all other studies (off-diagonal cells in Fig. 3a-b). Meanwhile, we applied a nested cross-validation procedure on the training study to calculate within-study accuracy (diagonal cells in Fig. 3a, b). In LODO validation, data from one study was set as the testing set, while data from the remaining three studies were pooled as the training set. The input features of the validation classifiers were the important features identified from the IFE analysis.

To evaluate whether the selected important features would achieve the best performances in study-to-study transfer validation and LODO validation, we constructed RF models with 3 different sets of input features, including (1) all ASVs, (2) differential ASVs and (3) important features. Then we sought to identify if there was a minimal set of important features that could achieve higher accuracy. A few of the top-ranking important features were always included in the minimal set in prior. We used the same methods as the study-to-study transfer validation and LODO validation and then calculated the average AUC of each testing study as each point in Fig. 3c, d. Finally, we compared the predictive values in the testing set across RF models with different sets of input features.

### Additional validation of independent studies and other diseases

As an external test, we used additional independent data to validate the performance of the selected important features to differentiate adenoma from cancer or control. The input features of RF models were the ASVs with the same taxonomy assignments as the selected important features as well as patient metadata (validation cohort2 without the patient metadata only used ASVs as input features).

To assess the specificity of the selected important features for colorectal adenoma, we investigated five non-CRC diseases. For each disease, we randomly drew 30 samples from the control group (excluding NAFLD and IBD diseases of which 15 samples were selected) as well as 30 samples from the cases, and added them to the validation cohort1 dataset in turns, labeled as controls. The random selection process was repeated ten times, and the validation AUC was computed accordingly.

### Functional profile analysis

The functions of gut microbiome were inferred from 16S rRNA sequences with Phylogenetic Investigation of Communities by Reconstruction of Unobserved States (PICRUSt2) as previously published(44). Functional metagenome profiles that have more than 20% samples with relative abundance < 1×10^−5^ and show up in less than three of the studies were removed. The differential analysis and generalized fold change calculations were performed on pathway profiles in the same way as on ASVs profiles (see data preprocessing method). Then, we evaluated the contribution of each ASV to overall differential pathways. The contribution was defined as the ratio of one ASV functional abundance to the total functional abundance of all ASVs in a given pathway.

### Quantitative PCR validation

To quantify the abundance and expression of genes from two selected biosynthesis, the qRT-PCR analysis was performed on 14 healthy controls, 12 adenoma and 30 CRC samples. For these samples, the gDNA was extracted with the FecalGen DNA Kit (Cat# e9604) according to the manufacturer’s instructions. We used the primes in the Supplementary Table 15 for candidate genes; standard primers F515 and R806 for 16S. To perform the qRT-PCR reaction, the final primer concentration was diluted to 0.5 μM including 5 ng of gDNA in a 20 μl final reaction volume with the SYBR Green qPCR Mix (Thermo Fisher Scientific). The used qRT-PCR program was as follows: pre-denaturation at 95 °C for 10 min; denaturation at 95 °C for 15 s for 40 cycles; annealing at 60°C for 60 s followed by melt curve analysis(14). The qRT-PCR analysis was to calculate 2^-^ΔΔCt values between candidate genes and 16S Ct values. The significance of the comparison between adenoma and control or CRC samples was tested by Wilcoxon test (*P*<0.05).

## List of abbreviations

ADP-heptose: ADP-L-glycero-beta-D-manno-heptose biosynthesis
ASVs: amplicon sequence variants
AUC: the area under the ROC curve
BMI: body mass index
CD: Crohn’s disease
CRC: Colorectal Cancer
ENA: European Nucleotide Archive
FIT: Faecal Immunochemical Test
FOBT: fecal occult blood test
IARC: International Agency for Research on Cancer
IBD: Inflammatory bowel disease
IBS: Irritable bowel syndrome
IFE: Iterative feature elimination
LODO: leave-one-dataset-out
LPS: lipopolysaccharide
MAFFT: Multiple Alignment using Fast Fourier Transform
MK-10: menaquinol-10 biosynthesis
NAFLD: Non-alcoholic fatty liver disease
NF-κB: Nuclear factor-κB
NSAID: Nonsteroidal anti-inflammatory drug
OTUs: operational taxonomic units
PICRUSt2: Phylogenetic Investigation of Communities by Reconstruction of Unobserved States 2
QIIME2: Quantitative Insights Into Microbial Ecology 2;
qRT-PCR: real-time quantitative PCR;
RF: Random Forest
ROC: Receiver operating characteristic
SRA: Sequence Read Archive
T2D: type 2 diabetes
UC: ulcerative colitis
WMS: whole metagenome shotgun sequencing

## Declarations

### Ethics approval and consent to participate

Not applicable.

### Consent for publication

Not applicable.

### Availability of data and materials

Raw fastq files are available through the Sequence Read Archive(SRA) and European Nucleotide Archive (ENA), with project ID: PRJNA389927, PRJEB6070, PRJNA290926, PRJNA362366, PRJNA534511, PRJNA280026, PRJEB28350, PRJNA544721, PRJNA541332 and PRJNA82111.

## Author’s contributions

LZ, NL, CT and RZ conceived and designed the project. Each author has contributed significantly to the submitted work. YW and NJ drafted the manuscript. RZ, YZ, DW, AW, SF, WG, YL, SC, XH, PL, CT, NL and LZ revised the manuscript. All authors read and approved the final manuscript.

