## Supplementary for "Identification of microbial markers across populations in early detection of colorectal cancer"

**Supplementary Information**

This file contains Supplementary Notes 1–9, Supplementary Methods, Supplementary Figures 1–15, Supplementary Tables 1-15, and Supplementary References.

Table of Contents

Supplementary Notes…………02

Supplementary Methods….......05

Supplementary Figures……….06

Supplementary Tables.………..21

Supplementary References…..109

**Supplementary Notes**

**Supplementary Note 1**

**Previous researches based on meta-analysis for identifying colorectal adenomas**

But so far, only Thomas et al(1) identified a few microbial markers of colorectal adenomas from WMS-based meta-analysis, and associate gut microbiome with CRC in principle(1). However, their classifier showed low accuracy in distinguishing adenomas from healthy controls (AUC=0.54) or from CRC (AUC=0.69)(1), probably due to the limited coverage of taxonomy and the high dependency on reference genomes in WGS taxonomic profiling(2). A recent meta-analysis study based on 16S rRNA, however, mainly investigated colonic cancerous tissues and identified some tissue-based microbial markers for colorectal adenoma(3). However, stool-based microbial markers of multiple studies still couldn’t properly distinguish adenoma samples from control.

**Supplementary Note 2**

**Differentially abundant** **ASVs** **between control and adenoma**

Specifically, there were six ASVs depleted in adenoma, including *Bifidobacterium longum*, *Lactococcus sp.*, *Aminipila butyrica*, etc. Besides, the abundances of 37 ASVs increased in adenoma compared with controls, which were assigned as *Eubacterium coprostanoligenes*, *Methanobrevibacter millerae*, *Christensenellaceae R-7 group sp.*, *Ruminiclostridium 5 sp.*, etc (Supplementary Table 1). The abundance change of adenoma-associated *Methanobrevibacter* was consistent with previous studies(4).

**Supplementary Note 3**

**Differentially abundant ASVs between adenoma and cancer**

Among these, 56 ASVs were in lower abundance in adenoma compared with cancer, associated with *Lachnoclostridium sp.*, *[Ruminococcus] gnavus group sp.*, *[Clostridium] scindens*, *Escherichia-Shigella sp.*, etc. The ASVs in higher abundance in adenoma than cancer were *Blautia obeum*, *Butyricicoccus faecihominis*, *Erysipelotrichaceae UCG-003 sp.*, *Dorea longicatena*, etc (Supplementary Table 2).

**Supplementary Note 4**

**High ranking biomarkers between healthy controls and CRC**

These ASVs were assigned as *Porphyromonas asaccharolytica*, *Fusobacterium* *nucleatum, Peptostreptococcus stomatis*, *Bacteroides dorei*,etc, which were also reported as CRC biomarkers in previous researches(5). Notably, the ASVs ranked in the top 10 important markers were assigned as *Fusobacterium* *nucleatum* and *Porphyromonas asaccharolytica*, which were also ranked as top markers in two recent meta-analysis of CRC based on WMS data. Moreover, the *Ruminococcus, Bacteroides, Hungatella and Peptostreptococcus* were also highlighted as biomarkers in at least one of the two studies (Supplementary Table 5). Then, we compared the markers across various CRC stages. There were six common biomarkers shared between patients with carcinomas and adenomas (Supplementary Fig. 7).

**Supplementary Note 5**

**Co-occurrence network analysis**

In the co-occurrence network of differential ASVs between adenoma and control, we found a large number of significantly positive correlations while only a few significantly negative correlations (Supplementary Fig. 8a and Supplementary Table 6). Notably, most of the negative correlations associated with the ASV were assigned as *Anaerostipes hadrus* (the 2nd ASV), belonging to *Anaerostipes* genus, which may protect against colon cancer in humans by producing butyric acid(6). It was enriched in control, and in adenoma but negatively correlated with multiple ASVs. The first and second ranking biomarkers between adenoma and control, *Christensenellaceae R-7 group sp.* and *Ruminococcaceae UCG-005 sp.,* were both related to several ASVs in this network, which amplified the importance of these two biomarkers. Clustering coefficient analysis via MCODE(7) from Cytoscape identified a module with the highest score containing 12 nodes and 33 interactions (Supplementary Fig. 8b). In this module, *Ruminococcaceae UCG-005 sp.* was associated with a wide range of nodes, including *Ruminococcaceae UCG-002 sp.* (the 15th ASV), *Methanobrevibacter millerae sp.* (the 35th ASV), *Ruminococcaceae UCG-014* (the 31st ASV), *Eubacterium ruminantium* and so on, which further indicated that this microbiota was a pivotal biomarker between adenoma and control.

Additonally, we investigated the co-occurrence network of differential ASVs between adenoma and CRC (Supplementary Fig. 8c and Supplementary Table 7). This network showed a strong negative association between adenoma-enriched and CRC-enriched ASVs. In contrast, positive correlations among the adenoma-enriched ASVs or CRC-enriched ASVs are observed in general. Our network analysis identified key interactions among several prominent taxonomic members. For example, *[Clostridium] scindens* and *Blautia sp.*, which were the top-ranking members of biomarkers for distinguishing adenoma from CRC, maintained strong negative relationships with several adenoma-enriched ASVs. Two modules were identified by the MCODE from this network (Supplementary Fig. 8d, e). One module comprised 19 nodes and 63 edges obtains a network score of 7.08. In this module, the biomarkers of *[Eubacterium] ventriosum group sp.*, *Lachnospira pectinoschiza*, *Eubacterium ruminantium*, *Blautia faecis*, *[Clostridium] scindens* and *Blautia sp.* were correlated with multiple nodes. Remarkably, all these biomarkers which belong to *Lachnospiraceae* family showed that the *Lachnospiraceae* family played an important role in this sub-network. Moreover, the other small module contained 4 nodes and 4 edges. Two nodes were biomarkers assigned as *Porphyromonas* genus which was a known pathogenic genus in CRC(8). In summary, our results suggested that most of the identified biomarkers have functional importance in the network.

**Supplementary Note 6**

**Improved adenoma diagnostic ability by combining with non-invasive clinical screening tests**

To compare the diagnostic performance of selected important features with FIT, the most widely used non-invasive stool test, we collected the publicly available FIT samples (including 172 control individuals and 198 adenoma patients) from the US2 study(9). The performance of the RF model constructed with FIT being the only one feature for distinguishing adenoma from control is 0.60 (AUC). The model constructed with important features tested on this study was proved to be superior to that of the FIT, with the AUC of 0.78. Moreover, the combination of FIT with the important features elevated the diagnostic accuracy for adenoma (about 3%) and reached the best performance of 0.81 (AUC) (Supplementary Fig. 11). Altogether, our results demonstrate that the microbial-derived biomarker panel is superior to FIT for detecting colorectal adenoma and their combination can somehow improve the quality of non-invasive diagnosis of adenoma.

**Supplementary Note 7**

**Differential pathways in adenoma vs. control and adenoma vs. CRC sequences**

In detail, a series of pathways involved in Carbohydrate Biosynthesis (e.g. ADP-L-glycero-beta-D-manno-heptose (ADP-heptose) biosynthesis), Inorganic Nutrient Metabolism, Nucleoside and Nucleotide Biosynthesis were enriched in adenoma compared with control, whereas, pathways of Aromatic Compound Degradation, Secondary Metabolite Biosynthesis were decreased in adenoma samples compared with healthy controls. Meanwhile, compared to adenoma, pathways mainly involved in Cofactor, Prosthetic Group, Electron Carrier, and Vitamin Biosynthesis (e.g. menaquinol-10 biosynthesis (MK-10)), Amino Acid Degradation and Fermentation increased in cancer. Moreover, Cell Structure Biosynthesis and Fatty Acid and Lipid Biosynthesis/Degradation pathways decreased in adenoma compared with cancer.

**Supplementary Note 8**

**CRC-associated markers limit themselves to detect colorectal adenoma**

The ASV classified as *Eubacterium ruminantium* is the only one common adenoma-associated marker while *Porphyromonas sp. HMSC077F02*, *Lachnospira pectinoschiza*, *Hungatella hathewayi* and etc are common cancer-associated markers (Supplementary Fig. 7). Fusobacterium, one of the universal biomarkers between control samples and patients with CRC in our cancer-control model and the two recent CRC meta-analysis(1, 10), is neither a significantly different bacteria nor a biomarker between controls and adenomas. It had also been reported that the diagnostic capability of *Fusobacterium sp.* for colorectal adenoma was not as good as stains ‘*m3*’ belonging to the *Lachnoclostridium sp.*(11). These results suggested that the CRC-associated markers limit themselves to detect colorectal adenoma and highlighted the importance of identification on adenoma-specific signatures. In addition, the adenoma-specific markers may contribute to the early screening and significantly reduce the risk of subsequent CRC.

**Supplementary Note 9**

**Insight of microbial-derived markers for clinical diagnosis of colorectal adenoma**

Taken together, through extensive and statistically rigorous validation, we identified microbial-derived markers for distinguishing adenoma from healthy control and CRC across multiple studies. Independent validation confirmed that the microbial-derived markers exhibted high accuracy and specificity on detecting adenoma. These microbial-derived markers may contribute to the non-invasive clinical diagnosis of colorectal adenoma and could be targeted to suppress CRC. Furthermore, different functions of microbiome are discovered in the development of CRC carcinogenesis. The alteration of microbiome mediated the activation of inflammation in adenoma while the disordered microbiome played a compensatory effect via elevated Vitamin K2 production in CRC carcinogenesis.

**Supplementary Mothods**

**Data collection for non-CRC studies**

Sequencing data of four non-CRC studies were also utilized to evaluate the specificity of adenoma features. These datasets are mapped to five diseases (Non-alcoholic fatty liver disease (NAFLD)(12), Type 2 diabetes (T2D)(13), Crohn’s disease(CD) (14), Ulcerative colitis (UC) (14) and Irritable bowel syndrome (IBS)) (Supplementary Table 8).

**Co-occurrence networks**

To examine co-occurrence networks in bacterial communities, network analysis was constructed on the relative abundance of differential ASVs. Spearman’s correlation coefficients with P values < 0.01 (defined as significant) and with a magnitude of 0.15 (control versus adenoma), 0.2 (adenoma versus cancer) or above were selected for further visualization in Cytoscape (version 3.8.0). Modular structure and groups of highly interconnected nodes were analyzed using the MCODE application with standard parameters(7).

**The diagnostic ability of FIT for colorectal adenoma**

To compare the diagnostic ability of traditional non-invasive tests, FIT, and the identified important features for colorectal adenoma, we constructed the RF models using important features, FIT and their combination for differentiating adenoma from control. The parameters of the RF models were kept the same as before (see method multivariable statistical modeling).

**Supplementary Figures**

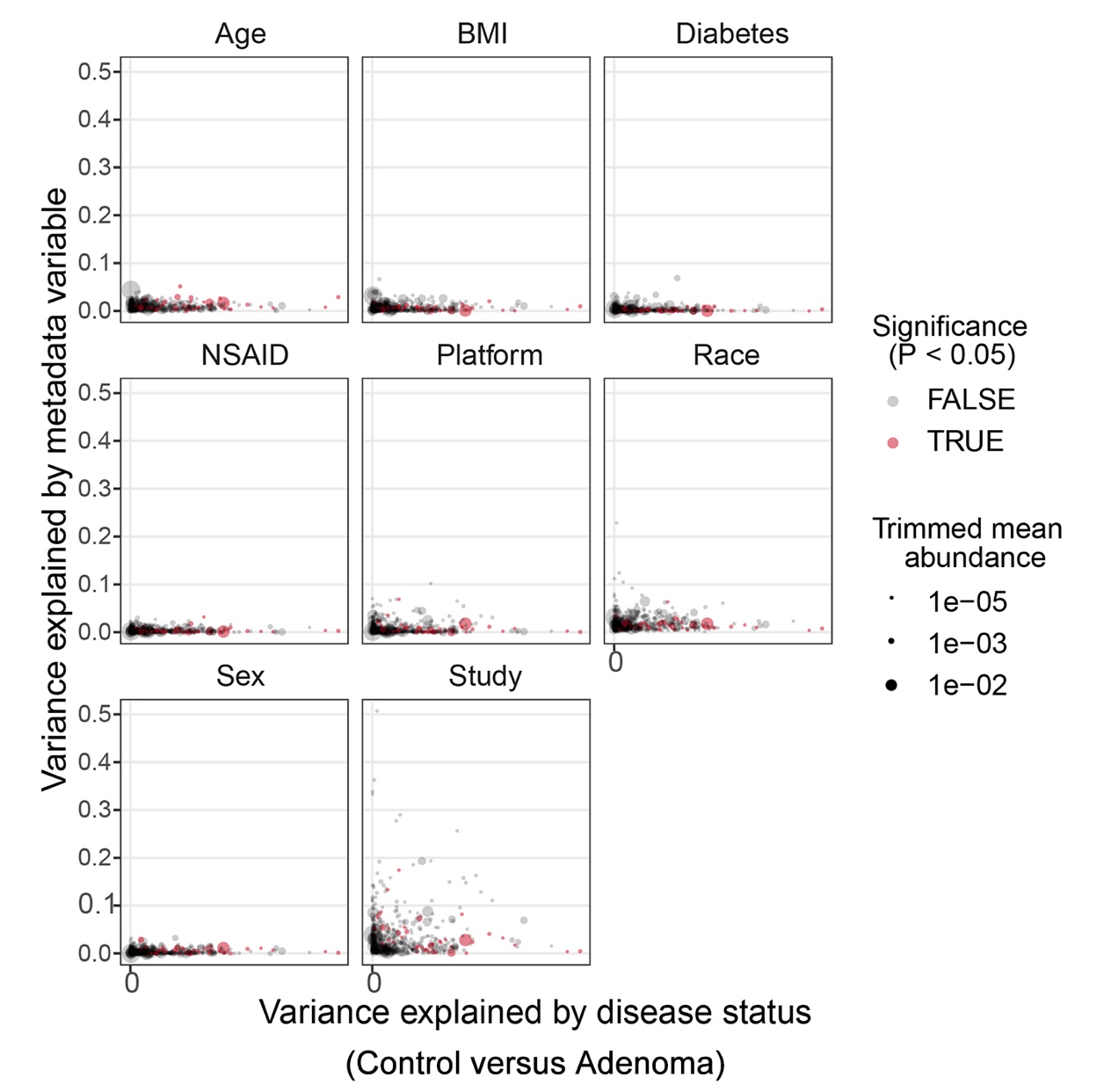

**Supplementary Fig. 1 |** **Variance explained by putative confounding factors and disease status (****Control versus Adenoma)**

Variance explained by disease status (Control versus Adenoma) is plotted against variance explained by different potential confounders (Age, BMI, Diabetes, NSAID, Platform, Race, Gender and Study) for individual ASVs. The abundance of each ASV is represented by different size of dot; the differentially abundant ASVs identified in the meta-analysis are highlighted in red; For the confounder analysis, the BMI was split into lean/overweight/obese according to conventional cutoffs; The variance explained by disease status was computed for all data *P* values between control and adenoma were calculated by blocked Wilcoxon tests (see methods).

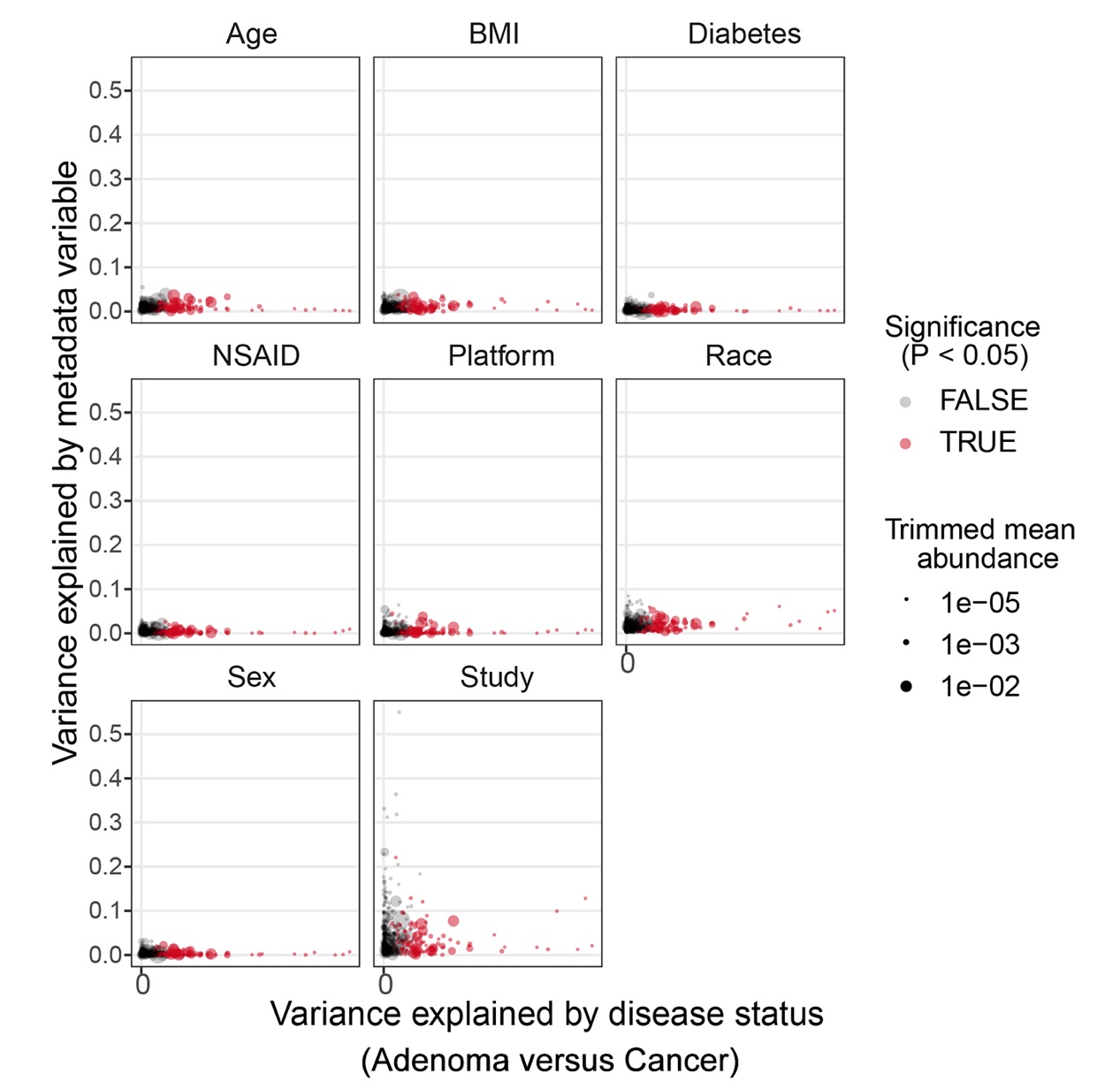

**Supplementary Fig. 2 | Variance explained by putative confounding factors and disease status (Adenoma versus Cancer)**

Variance explained by disease status (Adenoma versus Cancer) is plotted against variance explained by different potential confounders (Age, BMI, Diabetes, NSAID, Platform, Race, Gender and Study) for individual ASVs. The abundance of each ASV is represented by different size of dot; the differentially abundant ASVs identified in the meta-analysis are highlighted in red; For the confounder analysis, the BMI was split into lean/overweight/obese according to conventional cutoffs; The variance explained by disease status was computed for all data *P* values between control and adenoma were calculated by blocked Wilcoxon tests (see methods).

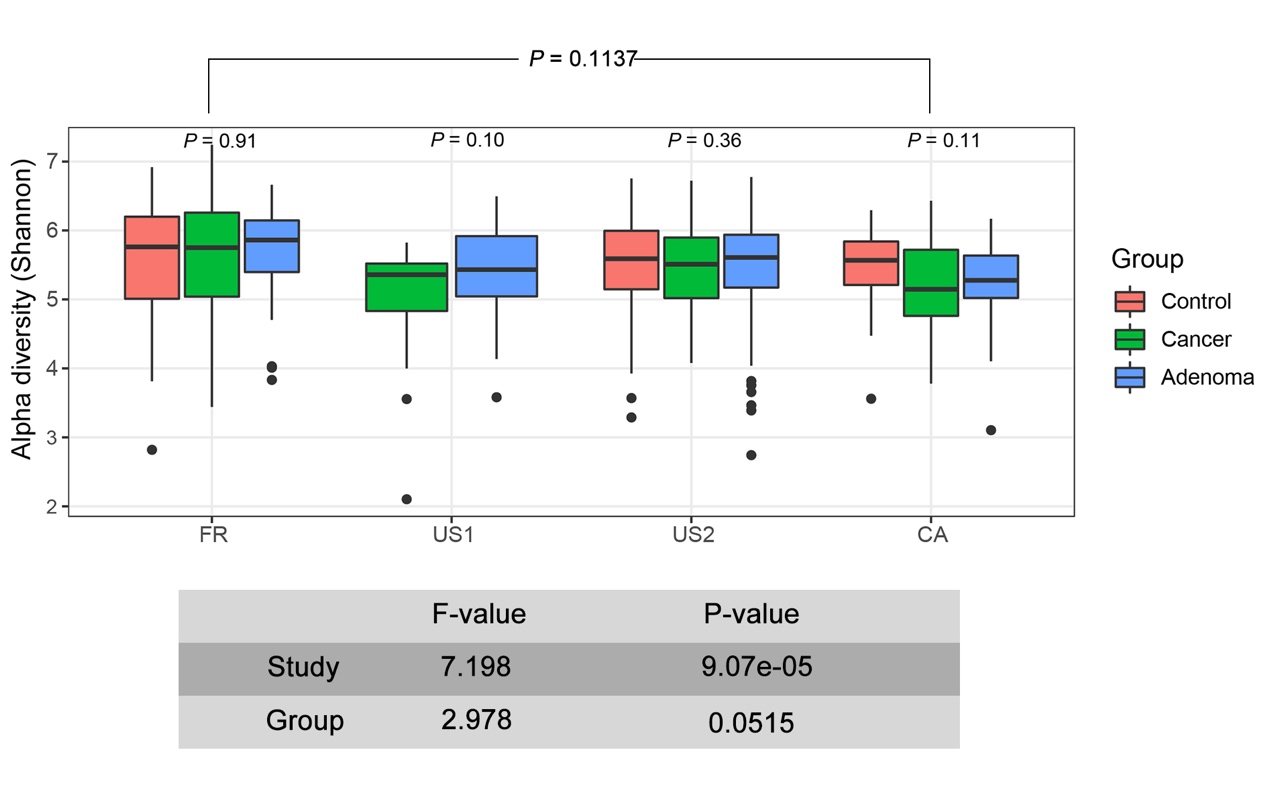

**Supplementary Fig. 3 | Study heterogeneity shows a significant influence on alpha diversity**

Alpha diversity as measured with the Shannon index was computed for all ASVs. *P* values were computed using a two-sided Wilcoxon test, while the overall *P* value (on top) was calculated using a two-sided blocked Wilcoxon test*.* The ANOVA F-values below the panel were calculated using the R function ‘aov’.

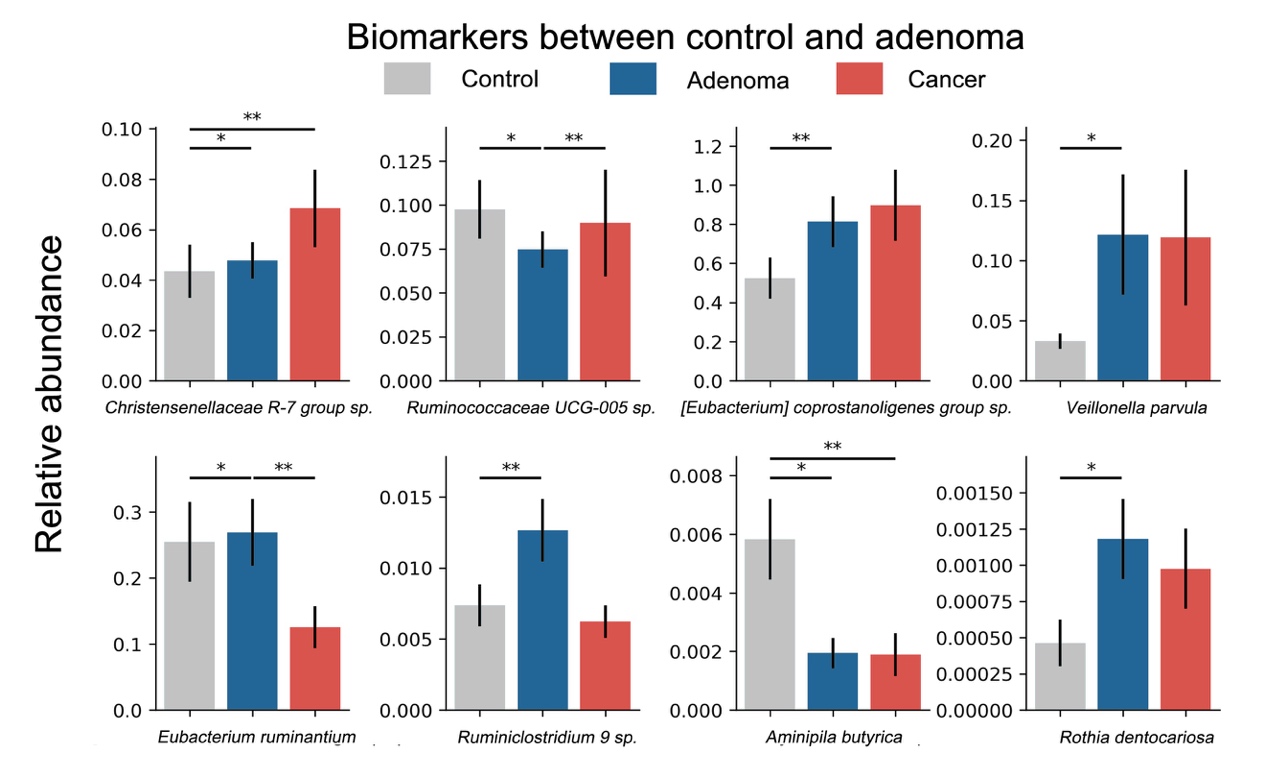

**Supplementary Fig. 4 | Relative Abundances of Biomarkers between control and adenoma**

Bar plots depicting the relative abundance of 8 biomarkers used to distinguish samples between healthy controls and adenomas. *P* values were calculated with a blocked Wilcoxon test, *: *P*<0.05 and **: *P*<0.01.

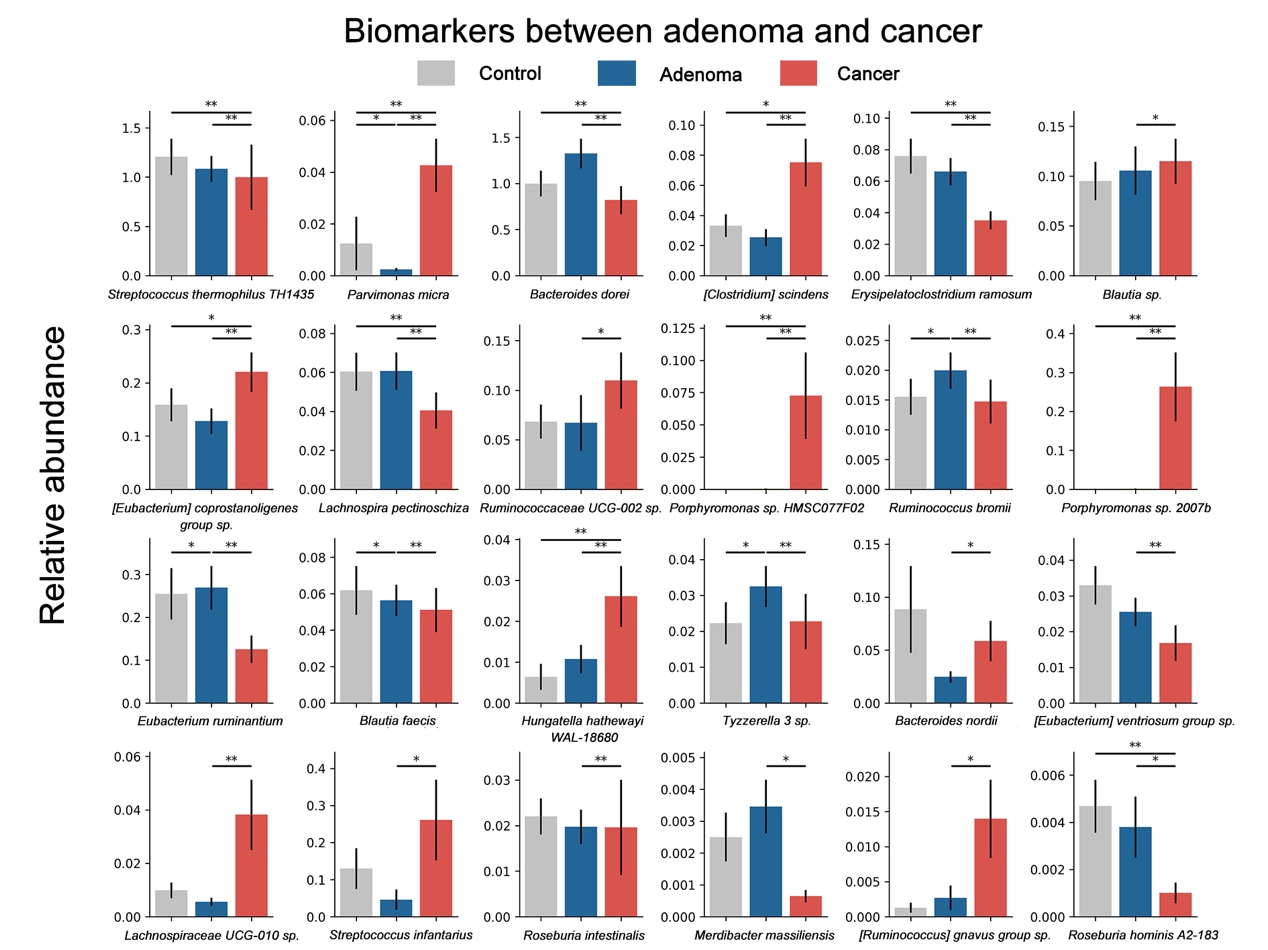

**Supplementary Fig. 5 | Relative Abundances of Biomarkers between adenoma and cancer**

Bar plots depicting the relative abundances of 24 biomarkers used to distinguish samples between adenomas and cancers. *P* values were calculated with blocked Wilcoxon test, *: *P*<0.05 and **: *P*<0.01.

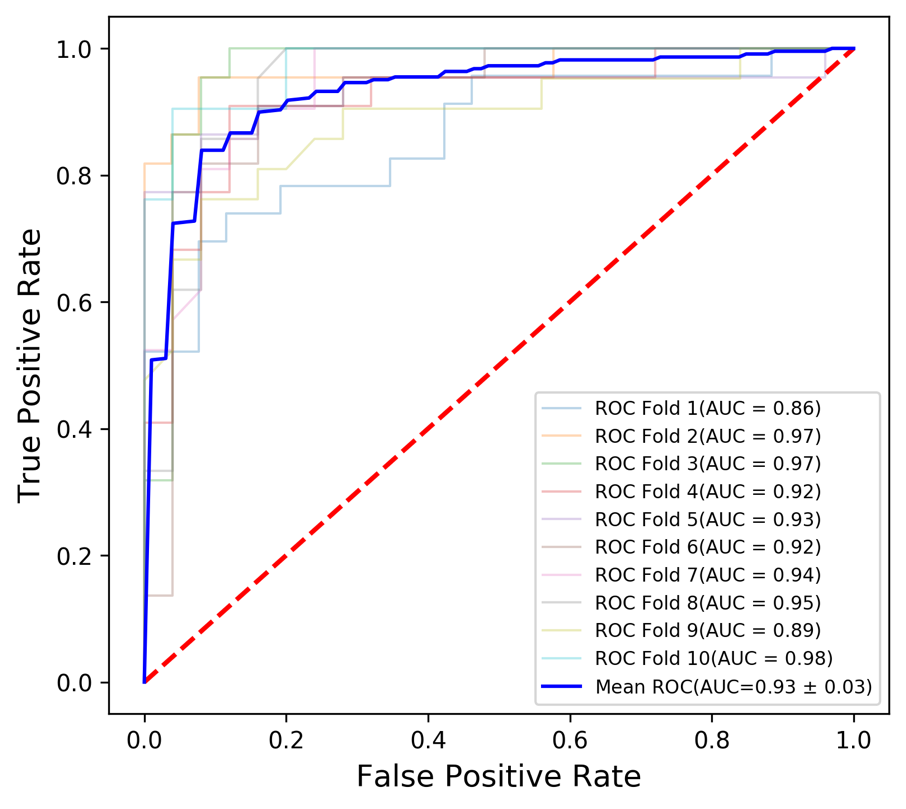

**Supplementary Fig. 6 | Performance of the RF Models for CRC detection**

Receiver operating characteristic (ROC) curve of the RF model (Control versus Cancer) constructed using the relative abundances of the 35 ASVs together with age and BMI in meta-analysis.

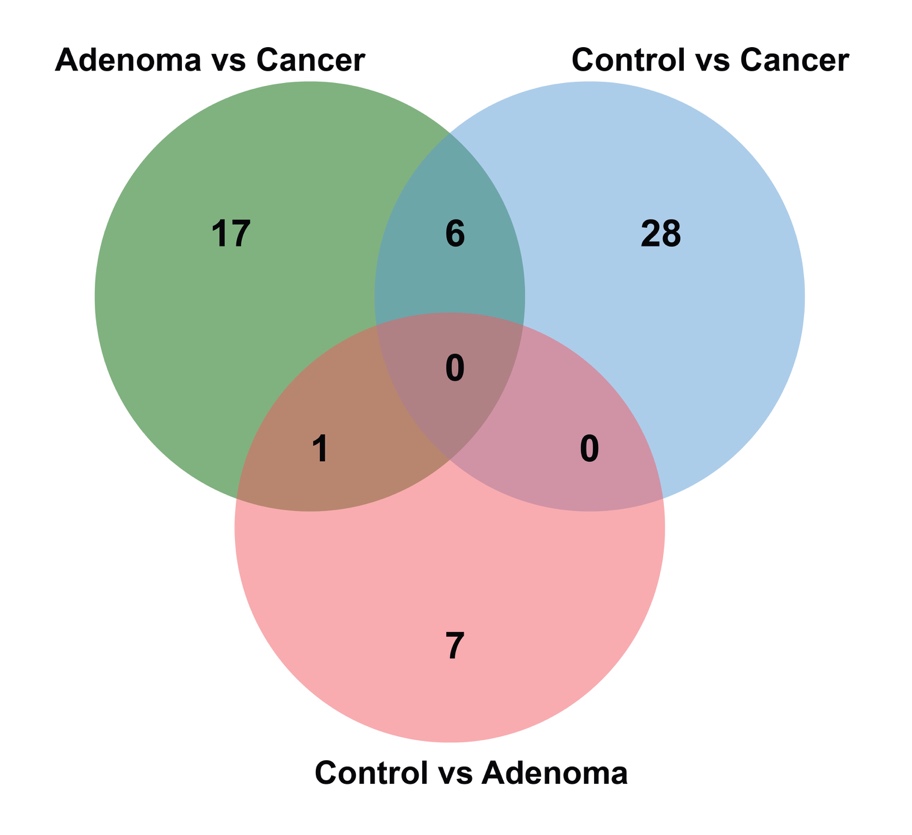

**Supplementary Fig. 7 | Overlap of three sets of biomarkers in Venn diagram**

Venn diagram shows the overlap of three sets of biomarkers for each two status.

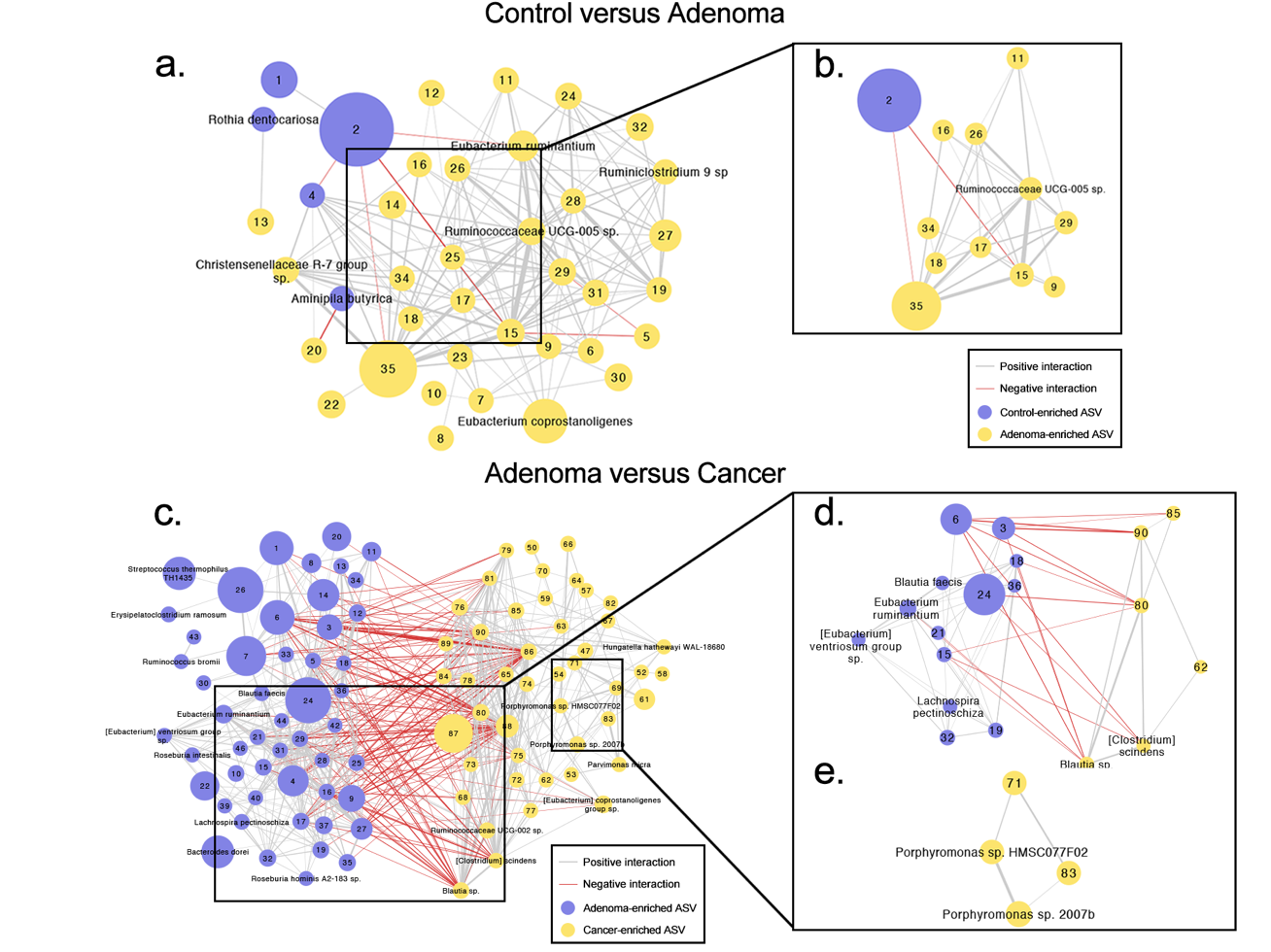

**Supplementary Fig. 8 | Microbial correlation networks for differential ASVs**

Correlation network of differential ASVs between adenoma and (a) control (n = 43 differential ASVs) or (c) CRC (n = 117 differential ASVs). Correlation coefficients were calculated by the Spearman algorithm. Modules (b) or (d-e) were analyzed using the MCODE application from (a) or (c). Node size represents mean ASV abundance; biomarkers are denoted by ASVs annotated to species; other differential ASVs are denoted by node numbers accordingly; Edges indicate correlations: the edge thickness represents the magnitude and the color represents the sign (gray are positive correlations, red are negative correlations).

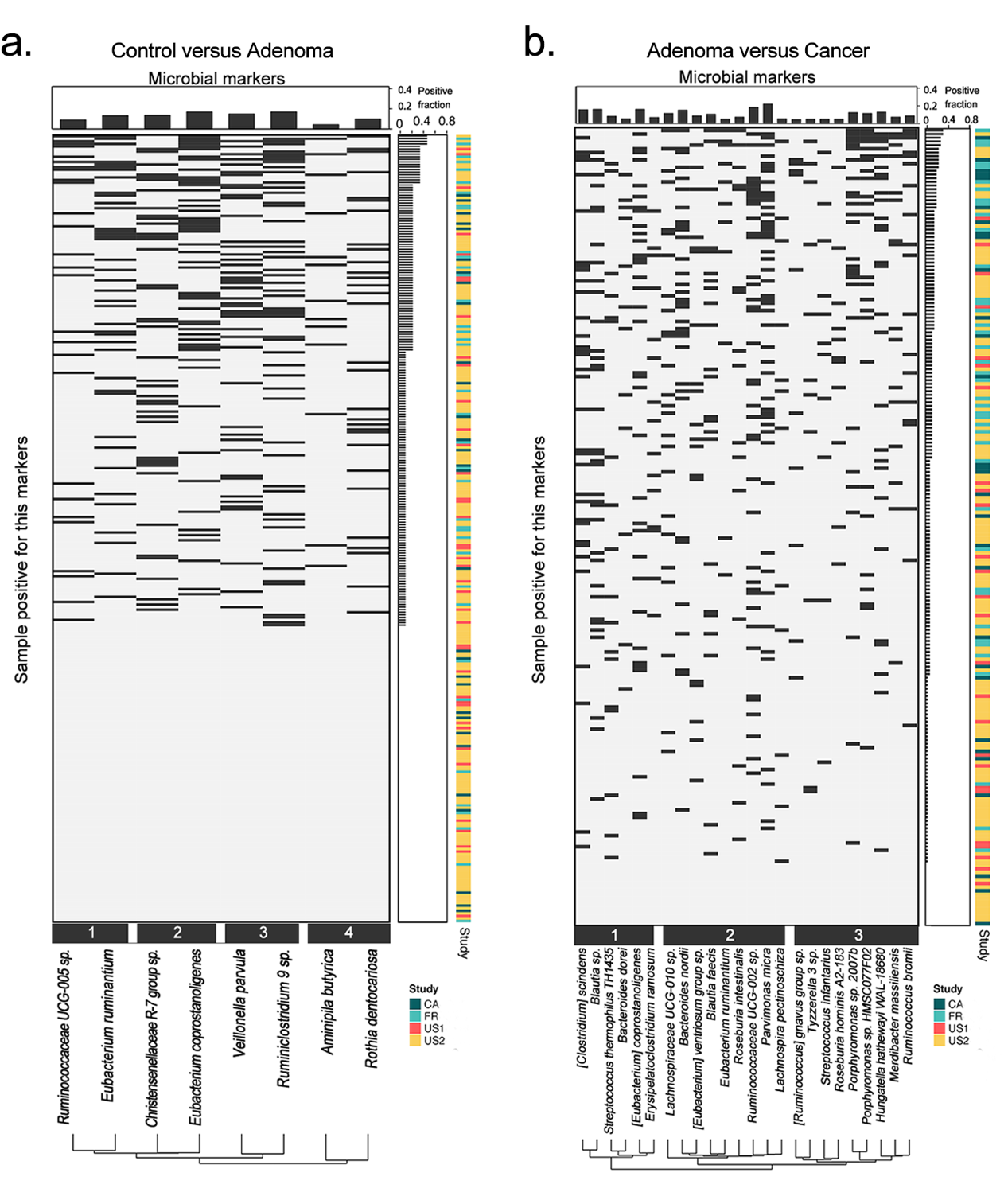

**Supplementary Fig. 9 | Co-occurrence analysis of biomarkers for distinguishing adenoma from control or adenoma**

a, b, For all patients with (a) adenoma (n = 307 samples) or (b) CRC (n = 217), the heatmap demonstrates whether the respective sample is positive for each of the biomarkers. Samples are ordered by the sum of positive biomarkers, and the biomarkers are clustered for four clusters (a) or three clusters (b) based on the Jaccard index of positive samples.

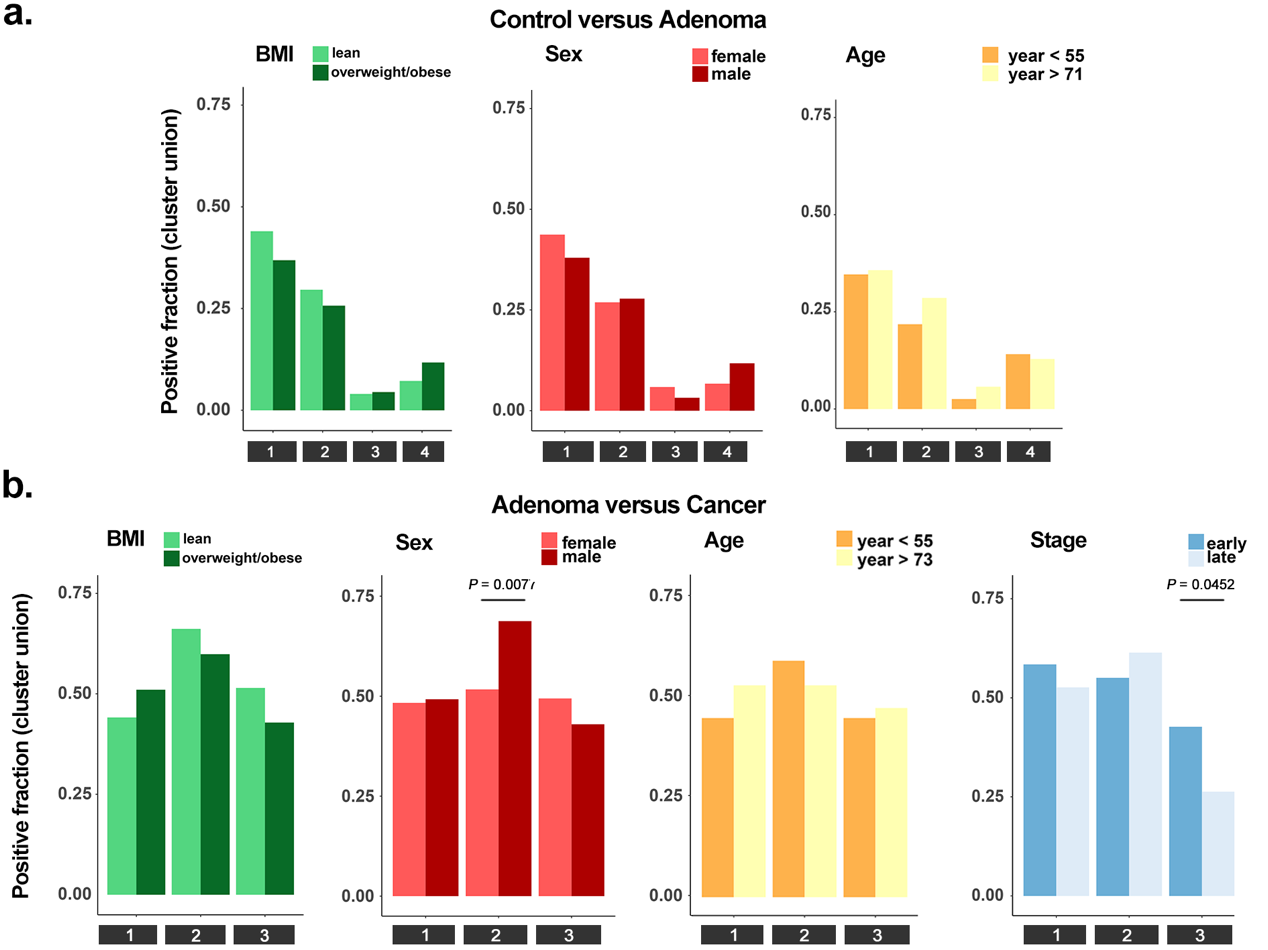

**Supplementary Fig. 10 | Co-occurrence of biomarkers identified clusters linked to different patient characteristics**

a, b, The barplots manifested the positive fraction for clusters of biomarkers between adenoma and (a) control (n = 8 biomarkers) or (b) CRC (n = 24 biomarkers) broken down by patient subgroups based on sex (a-b), age (a-b), BMI (a-b) and Stage (b), respectively. The significant associations between adenoma subgroups (a) or CRC subgroups (b) and biomarker clusters were identified by the Cochran–Mantel–Haenszel test blocked for “study”.

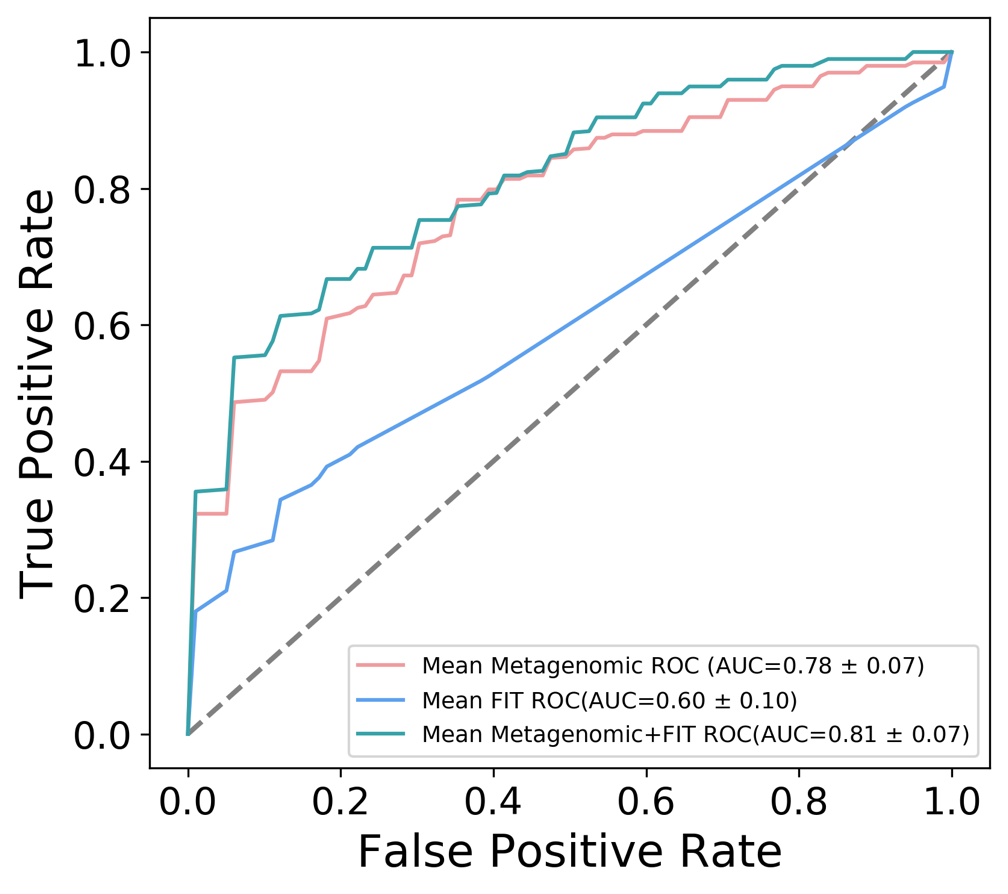

**Supplementary Fig. 11 | Improved adenoma diagnostic ability by combining with FIT tests**

AUC values for the prediction of colorectal adenoma using selected important features, FIT or a combination of both, AUC value is highest for the combination test.

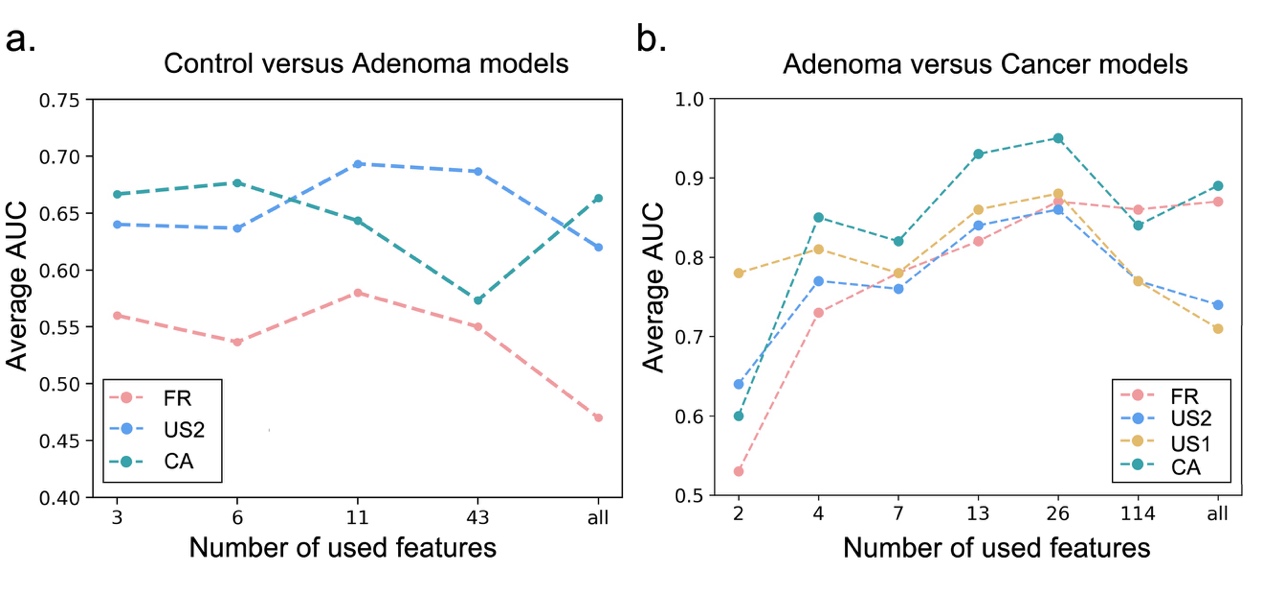

**Supplementary Fig. 12 | Prediction performances of** **LODO validation classifiers different sets of features**

a, b, Average AUC of LODO validation classifiers for control versus adenoma (a) and adenoma versus cancer (b) at different sets of features. The x-axis in (a) and (b) indicate different sets of features: All (a-b): all ASVs; 43 (a) and 114 (b): differentially abundant ASVs; 11 (a) and 26 (b): all important features; other top-ranking important features. The different studies were indicated in different colors.

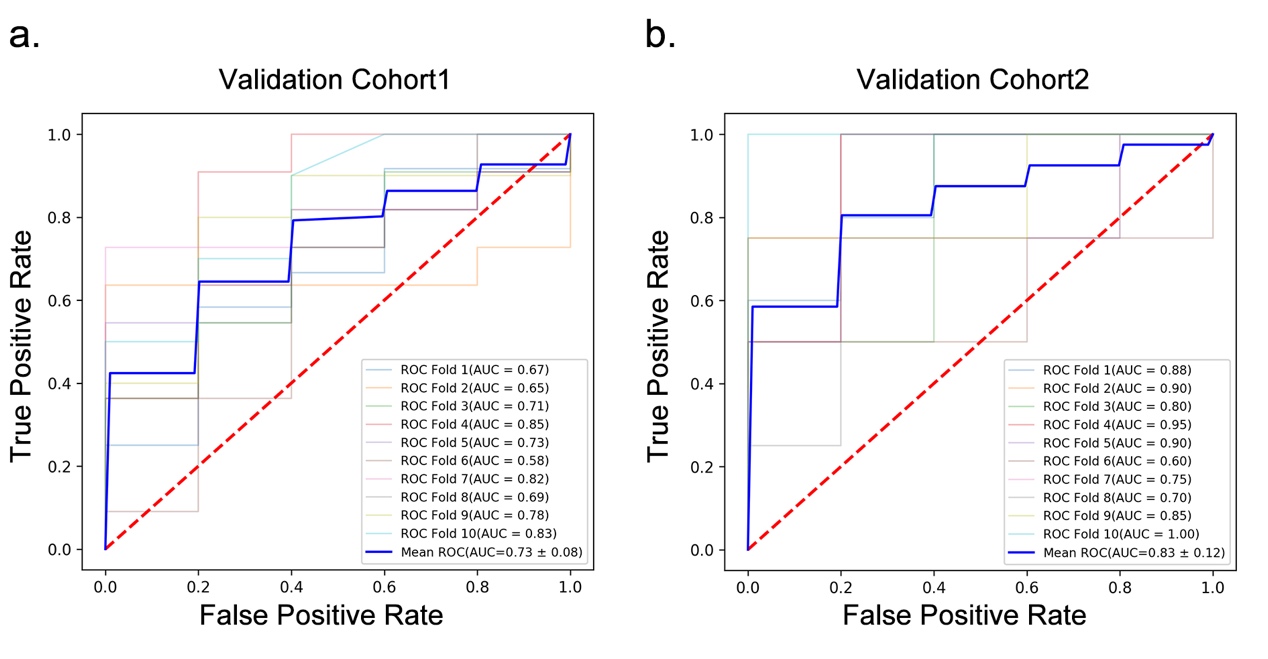

**Supplementary Fig. 13 | Validation performance of two independent cohorts**

a, b, Validation performance of two independent cohorts for discriminating adenoma from control (a) and CRC (b).

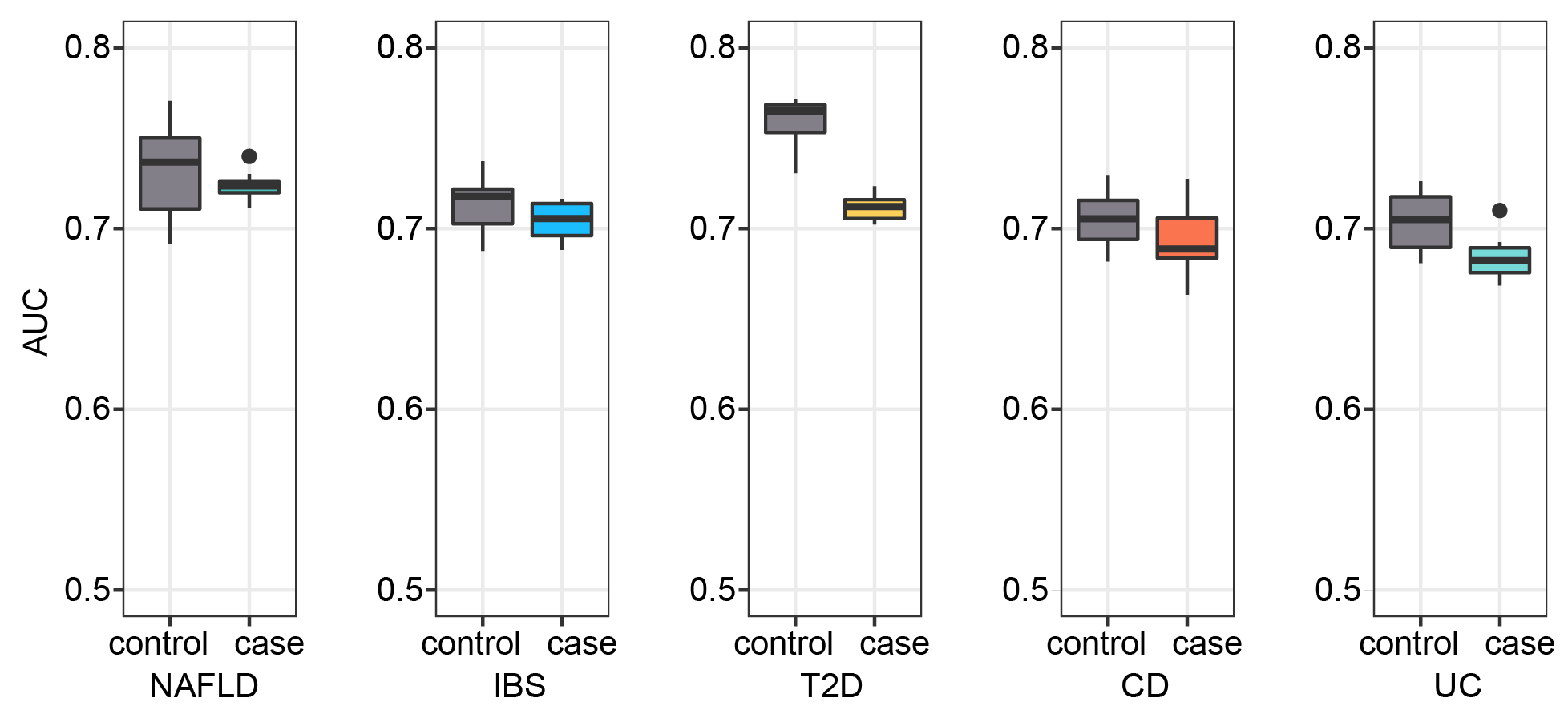

**Supplementary Fig. 14 | Colorectal adenoma specificity of predictive models**

Prediction abilities as AUC values on the validation cohorts when adding an external set of case and control samples from studies of non-CRC diseases (NAFLD, T2D, CD, UC and IBS).

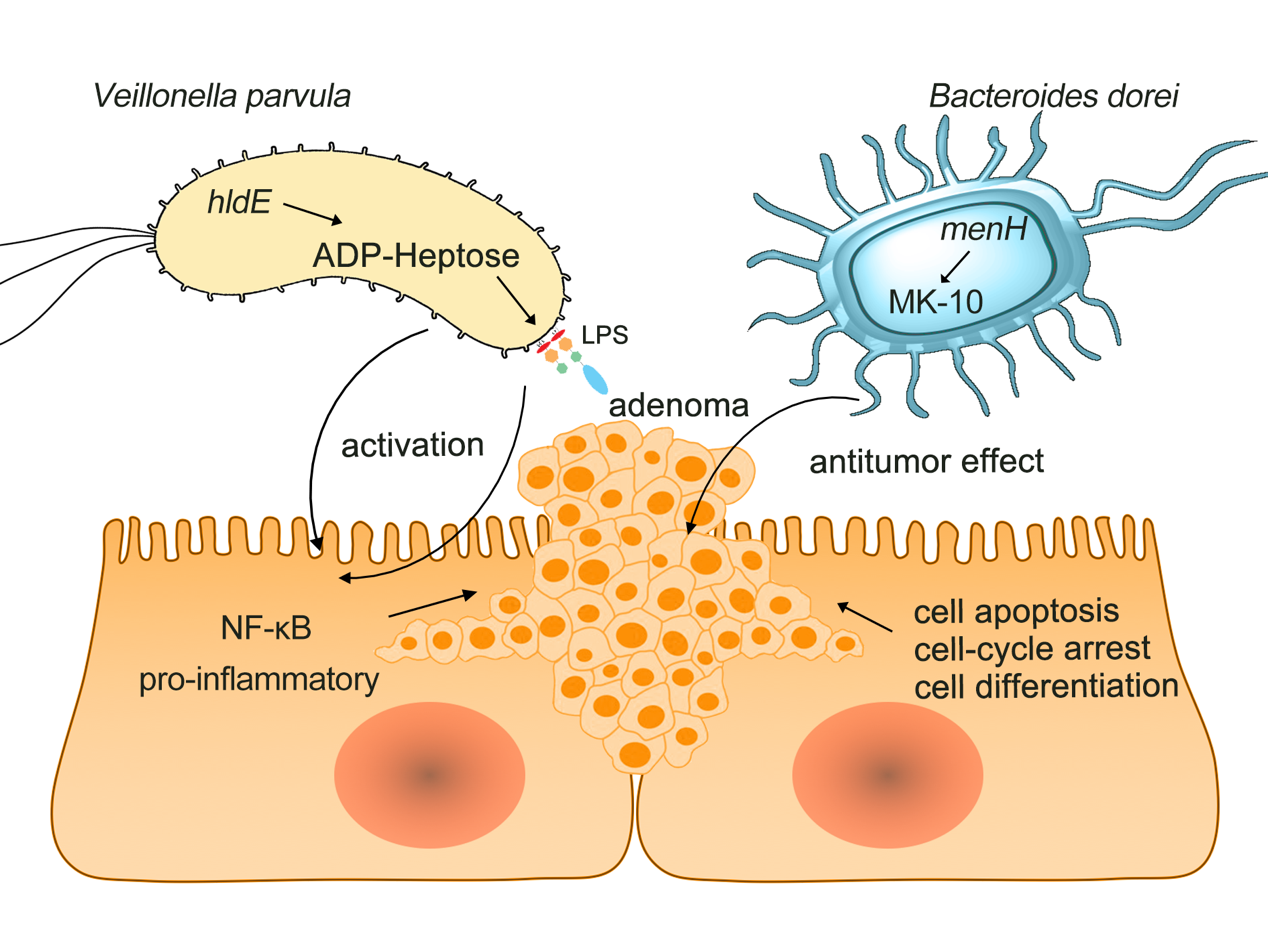

**Supplementary Fig. 15 | Potential microbial driver during adenoma-control and adenoma-CRC sequences**

The biosynthesis of ADP-heptose coded by *hldE* and etc genes is associated with the activation of NF-κB and inducing a strong pro-inflammatory response. While the MK-10 pathway coded by *menH* and etc genes played a key role in antitumor effect via cell-cycle arrest, cell differentiation and cell apoptosis.

**Supplementary Table 1 | Differentially abundant ASVs between control and adenoma (P < 0.05)**

| **ASV ID*** | **Taxon^†^** | **pvalue-FR^‡^** | **pvalue-US2^‡^** | **pvalue-CA^‡^** | **pvalue-meta^#^** | **GFOLD-meta^$^** |
| --- | --- | --- | --- | --- | --- | --- |
| 35ffcc3b809d667286737d79670b8de5 | *D_0__Bacteria;D_1__Actinobacteria;D_2__Actinobacteria;D_3__Bifidobacteriales;D_4__Bifidobacteriaceae;D_5__Bifidobacterium;D_6__Bifidobacterium longum* | 0.0247 | 0.1457 | 0.6272 | 0.0152 | -0.3658 |
| 33518e48b174ac428cb7be1a216c6c6e | *D_0__Bacteria;D_1__Firmicutes;D_2__Clostridia;D_3__Clostridiales;D_4__Lachnospiraceae;D_5__Anaerostipes;D_6__Anaerostipes hadrus* | 0.0050 | 0.0562 | 0.9470 | 0.0049 | -0.1781 |
| d114fb4c335125128be28401522dd41a | *D_0__Bacteria;D_1__Firmicutes;D_2__Bacilli;D_3__Lactobacillales;D_4__Streptococcaceae;D_5__Lactococcus;D_6__Lactococcus taiwanensis* | 0.3094 | 0.0318 | 0.8798 | 0.0329 | -0.1326 |
| 6e9ac49ab03e8cf905432ad62df55c30 | *D_0__Bacteria;D_1__Firmicutes;D_2__Clostridia;D_3__Clostridiales;D_4__Family XIII;D_5__Family XIII UCG-001;D_6__Aminipila butyrica* | 0.8882 | 0.0268 | 0.5440 | 0.0487 | -0.0251 |
| d83f60183d81253a505beaeef3cd168f | *D_0__Bacteria;D_1__Actinobacteria;D_2__Actinobacteria;D_3__Micrococcales;D_4__Micrococcaceae;D_5__Rothia;D_6__Rothia dentocariosa* | 0.3590 | 0.0004 | 0.9041 | 0.0310 | -0.0090 |
| a7e28b3f4879359a929a0da8c8557757 | *D_0__Bacteria;D_1__Firmicutes;D_2__Clostridia;D_3__Clostridiales;D_4__Clostridiaceae;D_5__Clostridiaceae;D_6__Clostridiaceae* | 0.3833 | 0.0523 | 0.0351 | 0.0233 | -0.0001 |
| 868be84b9daccd25a648f872c8d5e9ab | *D_0__Bacteria;D_1__Firmicutes;D_2__Negativicutes;D_3__Selenomonadales;D_4__Veillonellaceae;D_5__uncultured;D_6__uncultured organism* | NA | 0.0866 | 0.1608 | 0.0289 | 0.0000 |
| 1eaa3cb7baaac3220b0606e3621afd16 | *D_0__Bacteria;D_1__Bacteroidetes;D_2__Bacteroidia;D_3__Bacteroidales;D_4__Rikenellaceae;D_5__Alistipes;D_6__Faecalibacterium prausnitzii* | 0.5071 | 0.0129 | 0.9523 | 0.0432 | 0.0160 |
| da1d26d90a7443d34675778207d2c227 | *D_0__Bacteria;D_1__Firmicutes;D_2__Clostridia;D_3__Clostridiales;D_4__Ruminococcaceae;D_5__Ruminiclostridium 9* | 0.3748 | 0.0203 | 0.1121 | 0.0293 | 0.0198 |
| e93b313352019b8113e7fc66bf757917 | *D_0__Bacteria;D_1__Firmicutes;D_2__Clostridia;D_3__Clostridiales;D_4__Family XI;D_5__Parvimonas;D_6__Parvimonas micra* | 0.1969 | 0.0648 | 1.0000 | 0.0373 | 0.0384 |
| e4f195476bffa394e3048063f99360ea | *D_0__Bacteria;D_1__Bacteroidetes;D_2__Bacteroidia;D_3__Bacteroidales;D_4__Barnesiellaceae;D_5__Coprobacter;D_6__Coprobacter fastidiosus NSB1* | 0.3784 | 0.0301 | 0.6326 | 0.0387 | 0.0462 |
| b204a17b1a7f29074d81303d5ca68464 | *D_0__Bacteria;D_1__Firmicutes;D_2__Clostridia;D_3__Clostridiales;D_4__Ruminococcaceae;D_5__Phocea;D_6__Phocea massiliensis* | 0.0374 | 0.0890 | 0.1899 | 0.0240 | 0.0486 |
| 8ca0b9917d926bb6241ce6e23673fce4 | *D_0__Bacteria;D_1__Firmicutes;D_2__Clostridia;D_3__Clostridiales;D_4__Lachnospiraceae;D_5__Blautia;D_6__Blautia schinkii* | 0.5494 | 0.0949 | 0.7083 | 0.0276 | 0.0509 |
| 74792adac452eb3544cdc17064d6bbee | *D_0__Bacteria;D_1__Proteobacteria;D_2__Deltaproteobacteria;D_3__Desulfovibrionales;D_4__Desulfovibrionaceae;D_5__Desulfovibrio;D_6__Desulfovibrio piger* | 0.0999 | 0.2476 | NA | 0.0371 | 0.0613 |
| 2ef1e51ab1cf99a3c6417b05a060830e | *D_0__Bacteria;D_1__Firmicutes;D_2__Bacilli;D_3__Lactobacillales;D_4__Lactobacillaceae;D_5__Lactobacillus;D_6__Lactobacillus caviae* | 0.0410 | 0.0100 | 0.3545 | 0.0029 | 0.0643 |
| ad03570a6623c3fe543fb52f514b234f | *D_0__Bacteria;D_1__Firmicutes;D_2__Clostridia;D_3__Clostridiales;D_4__Ruminococcaceae;D_5__Ruminococcus 1* | 0.8245 | 0.0292 | 0.9701 | 0.0377 | 0.0653 |
| 3219af75ee8056ed06ba81e873b0d0cb | *D_0__Bacteria;D_1__Firmicutes;D_2__Clostridia;D_3__Clostridiales;D_4__Ruminococcaceae;D_5__Ruminococcaceae UCG-002;D_6__uncultured rumen bacterium* | 0.6035 | 0.0940 | 0.4936 | 0.0423 | 0.0706 |
| 2740cf2417c92847cc298cbd71dd1fcd | *D_0__Bacteria;D_1__Firmicutes;D_2__Negativicutes;D_3__Selenomonadales;D_4__Veillonellaceae;D_5__Veillonella;D_6__Veillonella parvula* | 0.1778 | 0.0336 | 0.4085 | 0.0120 | 0.0766 |
| 7427cc5ba277f690df113d56e5c6b538 | *D_0__Bacteria;D_1__Actinobacteria;D_2__Coriobacteriia;D_3__Coriobacteriales;D_4__Coriobacteriaceae;D_5__Olegusella;D_6__Olegusella massiliensis* | 0.4758 | 0.0049 | 0.4188 | 0.0268 | 0.0772 |
| 1c719e69d668107c05bdba29829217c5 | *D_0__Bacteria;D_1__Proteobacteria;D_2__Deltaproteobacteria;D_3__Desulfovibrionales;D_4__Desulfovibrionaceae;D_5__Desulfovibrio;D_6__Desulfovibrio longreachensis* | 0.1690 | 0.0095 | 0.8217 | 0.0024 | 0.0785 |
| cd604e575361dd5195144bf8e5007cb7 | *D_0__Bacteria;D_1__Firmicutes;D_2__Clostridia;D_3__Clostridiales;D_4__Ruminococcaceae;D_5__Fournierella;D_6__Fournierella massiliensis* | 0.0455 | 0.4275 | 0.6547 | 0.0198 | 0.0834 |
| bdb6ddef008a6d31c019642fb28d7c1d | *D_0__Bacteria;D_1__Firmicutes;D_2__Clostridia;D_3__Clostridiales;D_4__Ruminococcaceae;D_5__Ruminiclostridium 9;D_6__Anaerostipes hadrus* | 0.2369 | 0.0100 | 0.9637 | 0.0081 | 0.0898 |
| 74d519816d4cfb9cf4ec9605f76301d6 | *D_0__Bacteria;D_1__Firmicutes;D_2__Clostridia;D_3__Clostridiales;D_4__Family XIII;D_5__Family XIII UCG-001;D_6__uncultured bacterium* | 0.0937 | 0.0704 | 0.5689 | 0.0191 | 0.1035 |
| a333ee090a52b582e1a0e2dd776eb937 | *D_0__Bacteria;D_1__Firmicutes;D_2__Erysipelotrichia;D_3__Erysipelotrichales;D_4__Erysipelotrichaceae;D_5__Coprobacillus;D_6__Coprobacillus cateniformis JCM 10604* | 0.0883 | 0.0163 | 0.8905 | 0.0010 | 0.1064 |
| 9ea77e95fbfd1a99c9c28db3b03a01a8 | *D_0__Bacteria;D_1__Bacteroidetes;D_2__Bacteroidia;D_3__Bacteroidales;D_4__Barnesiellaceae;D_5__Barnesiella;D_6__Barnesiella intestinihominis* | 0.9681 | 0.0578 | 0.5968 | 0.0354 | 0.1141 |
| aae4c4ed528ae16ec417df816699cfdb | *D_0__Bacteria;D_1__Firmicutes;D_2__Clostridia;D_3__Clostridiales;D_4__Ruminococcaceae;D_5__Ruminococcaceae UCG-005* | 0.5740 | 0.1345 | 0.9709 | 0.0459 | 0.1203 |
| b0e56c25ca193e27096e8fb4eb560eee | *D_0__Bacteria;D_1__Firmicutes;D_2__Clostridia;D_3__Clostridiales;D_4__Lachnospiraceae;D_5__Eisenbergiella;D_6__Eisenbergiella tayi* | 0.3407 | 0.1454 | 0.6110 | 0.0267 | 0.1272 |
| 2de0f958e30d26f04cba8d2a920bbe46 | *D_0__Bacteria;D_1__Firmicutes;D_2__Clostridia;D_3__Clostridiales;D_4__Lachnospiraceae;D_5__Blautia;D_6__Blautia faecis* | 0.0108 | 0.4965 | 0.2207 | 0.0227 | 0.1420 |
| fec2da4ccc40b9ad85a020553063366b | *D_0__Bacteria;D_1__Proteobacteria;D_2__Gammaproteobacteria;D_3__Betaproteobacteriales;D_4__Burkholderiaceae;D_5__Oxalobacteraceae;D_6__Noviherbaspirillum agri* | 0.0117 | 0.0475 | 0.2380 | 0.0019 | 0.1426 |
| 54ec3041ed02d098b3c5e4395466c441 | *D_0__Bacteria;D_1__Firmicutes;D_2__Clostridia;D_3__Clostridiales;D_4__Ruminococcaceae;D_5__Ruminococcaceae UCG-002;D_6__uncultured organism* | 0.0740 | 0.2084 | 0.9591 | 0.0204 | 0.1473 |
| ec6732c2e0d4cf64b3d0350e7fe3defb | *D_0__Bacteria;D_1__Firmicutes;D_2__Clostridia;D_3__Clostridiales;D_4__Lachnospiraceae;D_5__Roseburia;D_6__Roseburia inulinivorans DSM 16841* | 0.0231 | 0.2017 | 0.7596 | 0.0042 | 0.1570 |
| c6b90711837508687841b1d5cbff2f65 | *D_0__Bacteria;D_1__Firmicutes;D_2__Clostridia;D_3__Clostridiales;D_4__Lachnospiraceae;D_5__uncultured;D_6__uncultured bacterium adhufec382* | 0.0492 | 0.2892 | 0.6326 | 0.0292 | 0.1677 |
| 51d4f17b8dda9f7d972bf3dd0ecc4e59 | *D_0__Bacteria;D_1__Bacteroidetes;D_2__Bacteroidia;D_3__Bacteroidales;D_4__Rikenellaceae;D_5__Alistipes;D_6__Alistipes obesi* | 0.8970 | 0.0341 | 0.3923 | 0.0225 | 0.1795 |
| eebc27bcf8bcb13a692001530892f5d5 | *D_0__Bacteria;D_1__Firmicutes;D_2__Negativicutes;D_3__Selenomonadales;D_4__Acidaminococcaceae;D_5__Phascolarctobacterium;D_6__uncultured Firmicutes bacterium* | 0.0029 | 0.3372 | 0.1699 | 0.0039 | 0.1821 |
| c5ec23b5e73a33133aa0fcf9acd58650 | *D_0__Bacteria;D_1__Firmicutes;D_2__Clostridia;D_3__Clostridiales;D_4__Ruminococcaceae;D_5__Ruminiclostridium 9* | 0.1209 | 0.0079 | 0.2711 | 0.0006 | 0.1951 |
| f50508546ae13143015f8c4cb976d0e4 | *D_0__Bacteria;D_1__Firmicutes;D_2__Clostridia;D_3__Clostridiales;D_4__Lachnospiraceae;D_5__[Eubacterium] ruminantium group;D_6__Eubacterium ruminantium* | 0.4122 | 0.0655 | 0.5024 | 0.0269 | 0.2161 |
| c97ea25ba069877eac769cb05968724c | *D_0__Bacteria;D_1__Firmicutes;D_2__Clostridia;D_3__Clostridiales;D_4__Ruminococcaceae;D_5__Ruminococcaceae UCG-014* | 0.2546 | 0.0544 | 0.5588 | 0.0120 | 0.2267 |
| fcffedae608fe4871be42936a1cb523a | *D_0__Bacteria;D_1__Firmicutes;D_2__Clostridia;D_3__Clostridiales;D_4__Lachnospiraceae;D_5__[Ruminococcus] torques group* | 0.0066 | 0.3108 | 0.3966 | 0.0201 | 0.2275 |
| 91587a85e342f8dba27f54e15ac0ea77 | *D_0__Bacteria;D_1__Firmicutes;D_2__Clostridia;D_3__Clostridiales;D_4__Lachnospiraceae;D_5__Tyzzerella 3;D_6__unidentified* | 0.6985 | 0.0166 | 0.0595 | 0.0306 | 0.2367 |
| aff3962bbd7607a2971161d9840ca409 | *D_0__Bacteria;D_1__Firmicutes;D_2__Clostridia;D_3__Clostridiales;D_4__Ruminococcaceae;D_5__Ruminiclostridium 5* | 0.0476 | 0.1037 | 0.0752 | 0.0126 | 0.3053 |
| 94928715987dcb9638dd5c0e8f9b20f4 | *D_0__Bacteria;D_1__Firmicutes;D_2__Clostridia;D_3__Clostridiales;D_4__Christensenellaceae;D_5__Christensenellaceae R-7 group* | 0.0465 | 0.0212 | 0.1612 | 0.0143 | 0.3150 |
| 44396f15b10f6577d61a11c6047c939f | *D_0__Archaea;D_1__Euryarchaeota;D_2__Methanobacteria;D_3__Methanobacteriales;D_4__Methanobacteriaceae;D_5__Methanobrevibacter;D_6__Methanobrevibacter millerae* | 0.3123 | 0.0332 | 0.4276 | 0.0216 | 0.3634 |
| 410e1eaa1468a3d40595898f671ba0b3 | *D_0__Bacteria;D_1__Firmicutes;D_2__Clostridia;D_3__Clostridiales;D_4__Ruminococcaceae;D_5__[Eubacterium] coprostanoligenes group;D_6__Eubacterium coprostanoligenes* | 0.0622 | 0.0451 | 0.4973 | 0.0018 | 0.4308 |

*** The ID of each amplicon sequence variant (ASV)**

**† Taxonomic information of each ASV**

**‡ Single-study P-value calculated by a two-sided Wilcoxon test**

**# Meta-analysis P-value calculated by a two-sided blocked Wilcoxon test (n=559 independent observations)**

**$ Mean of generalized fold changes across studies, GFOLD-meta >0: ASVs enriched in adenoma compared with control; <0: ASVs enriched in control compared with adenoma**

**Supplementary Table 2 | Differentially abundant ASVs between adenoma and cancer (P < 0.05)**

| **ASV ID*** | **Taxon^†^** | **pvalue-FR^‡^** | **pvalue-US1^‡^** | **pvalue-US2^‡^** | **pvalue-CA^‡^** | **pvalue-meta^#^** | **GFOLD-meta^$^** |
| --- | --- | --- | --- | --- | --- | --- | --- |
| 00a96fbd0ac34bfd245f9c24f8737f7d | *D_0__Bacteria;D_1__Firmicutes;D_2__Clostridia;D_3__Clostridiales;D_4__Lachnospiraceae;D_5__Blautia;D_6__Blautia obeum* | 0.3487 | 0.0069 | 0.6086 | 0.0009 | 0.0029 | -0.6457 |
| a18c0c17fce3d52dc60506e8730bc996 | *D_0__Bacteria;D_1__Firmicutes;D_2__Clostridia;D_3__Clostridiales;D_4__Ruminococcaceae;D_5__Butyricicoccus;D_6__Butyricicoccus faecihominis* | 0.0632 | 0.0080 | 0.1446 | 0.1016 | 0.0029 | -0.5078 |
| e865f29a716e8d51f83931befa951240 | *D_0__Bacteria;D_1__Firmicutes;D_2__Erysipelotrichia;D_3__Erysipelotrichales;D_4__Erysipelotrichaceae;D_5__Erysipelotrichaceae UCG-003* | 0.2860 | 0.0698 | 0.0087 | 0.1403 | 0.0003 | -0.5020 |
| 2e4f2b53b856c4def6d021d01f5abb70 | *D_0__Bacteria;D_1__Firmicutes;D_2__Clostridia;D_3__Clostridiales;D_4__Lachnospiraceae;D_5__Dorea;D_6__Dorea longicatena* | 0.0479 | 0.2522 | 0.1871 | 0.4089 | 0.0088 | -0.4835 |
| 69b90707f6f3da7941026fbe6a1597ce | *D_0__Bacteria;D_1__Firmicutes;D_2__Clostridia;D_3__Clostridiales;D_4__Lachnospiraceae;D_5__Lachnospiraceae FCS020 group;D_6__uncultured organism* | 0.0121 | 0.0320 | 0.2015 | 0.1010 | 0.0011 | -0.4742 |
| c48070f3061b086b60ff32f77e0002fa | *D_0__Bacteria;D_1__Firmicutes;D_2__Clostridia;D_3__Clostridiales;D_4__Ruminococcaceae;D_5__Subdoligranulum;D_6__Subdoligranulum variabile* | 0.1678 | 0.2192 | 0.1207 | 0.3304 | 0.0314 | -0.4570 |
| d76d59ec71de0e3b22da0c9cd564d41a | *D_0__Bacteria;D_1__Firmicutes;D_2__Clostridia;D_3__Clostridiales;D_4__Lachnospiraceae;D_5__Fusicatenibacter;D_6__Fusicatenibacter saccharivorans* | 0.0396 | 0.0098 | 0.0677 | 0.8883 | 0.0015 | -0.4405 |
| 1a9df3fbbc7fc8cecec4239cf801d2fc | *D_0__Bacteria;D_1__Firmicutes;D_2__Clostridia;D_3__Clostridiales;D_4__Ruminococcaceae;D_5__Ruminococcaceae UCG-013* | 0.0387 | 0.0203 | 0.0050 | 0.3181 | 0.0000 | -0.4289 |
| 9639a3291729a3758207b47715d9205f | *D_0__Bacteria;D_1__Firmicutes;D_2__Clostridia;D_3__Clostridiales;D_4__Ruminococcaceae;D_5__SubdoligranulumD_6__Subdoligranulum variabile* | 0.0201 | 0.0371 | 0.3962 | 0.8350 | 0.0183 | -0.4097 |
| caea322863a57bb2b14c7317fa120dc4 | *D_0__Bacteria;D_1__Firmicutes;D_2__Clostridia;D_3__Clostridiales;D_4__Lachnospiraceae;D_5__Lachnospiraceae NK4A136 group* | 0.1914 | 0.4164 | 0.0700 | 0.0224 | 0.0043 | -0.3935 |
| fd44d4cb468fd7dc9b3227867714ed87 | *D_0__Bacteria;D_1__Bacteroidetes;D_2__Bacteroidia;D_3__Bacteroidales;D_4__Bacteroidaceae;D_5__Bacteroides;D_6__Bacteroides dorei* | 0.4595 | 0.7252 | 0.0002 | 0.6272 | 0.0008 | -0.3615 |
| 394eda29c886632f514dd94b58381186 | *D_0__Bacteria;D_1__Proteobacteria;D_2__Gammaproteobacteria;D_3__Pasteurellales;D_4__Pasteurellaceae;D_5__Haemophilus;D_6__Haemophilus parainfluenzae ATCC 33392* | 0.2733 | 0.5881 | 0.0000 | 0.2166 | 0.0000 | -0.3596 |
| f50508546ae13143015f8c4cb976d0e4 | *D_0__Bacteria;D_1__Firmicutes;D_2__Clostridia;D_3__Clostridiales;D_4__Lachnospiraceae;D_5__[Eubacterium] ruminantium group;D_6__Eubacterium ruminantium* | 0.1323 | 0.2497 | 0.0619 | 0.1166 | 0.0061 | -0.3572 |
| 2c0c62ea09b2efe01bacdcbcf558d2b8 | *D_0__Bacteria;D_1__Firmicutes;D_2__Clostridia;D_3__Clostridiales;D_4__Lachnospiraceae;D_5__Lachnoclostridium;D_6__uncultured Firmicutes bacterium* | 0.2154 | 0.0261 | 0.0354 | 0.6019 | 0.0015 | -0.3400 |
| e9f900358bb9297e10b9fbb1321b1e6e | *D_0__Bacteria;D_1__Firmicutes;D_2__Erysipelotrichia;D_3__Erysipelotrichales;D_4__Erysipelotrichaceae;D_5__Erysipelatoclostridium;D_6__Erysipelatoclostridium ramosum* | 0.0784 | 0.0992 | 0.0673 | 0.1847 | 0.0019 | -0.3358 |
| 6ec1023583647a401269990c4e8ad716 | *D_0__Bacteria;D_1__Firmicutes;D_2__Clostridia;D_3__Clostridiales;D_4__Lachnospiraceae;D_5__Lachnospira;D_6__Lachnospira pectinoschiza* | 0.0474 | 0.0499 | 0.0377 | 0.9418 | 0.0045 | -0.3301 |
| 0a035a7ec1f4a194dd7fbda3c5660db0 | *D_0__Bacteria;D_1__Firmicutes;D_2__Clostridia;D_3__Clostridiales;D_4__Lachnospiraceae;D_5__Lachnoclostridium;D_6__human gut metagenome* | 0.1971 | 0.0069 | 0.0238 | 0.7240 | 0.0012 | -0.3240 |
| 6851a4ee264b56be2fff4686ce269907 | *D_0__Bacteria;D_1__Firmicutes;D_2__Clostridia;D_3__Clostridiales;D_4__Lachnospiraceae;D_5__[Eubacterium] hallii group* | 0.2122 | 0.6622 | 0.1649 | 0.0203 | 0.0324 | -0.3229 |
| 2de0f958e30d26f04cba8d2a920bbe46 | *D_0__Bacteria;D_1__Firmicutes;D_2__Clostridia;D_3__Clostridiales;D_4__Lachnospiraceae;D_5__Blautia;D_6__Blautia faecis* | 0.0146 | 0.1683 | 0.3829 | 0.4540 | 0.0062 | -0.3161 |
| 421cbd1d71d34704c6d53377261e213c | *D_0__Bacteria;D_1__Firmicutes;D_2__Clostridia;D_3__Clostridiales;D_4__Lachnospiraceae;D_5__[Eubacterium] ventriosum group* | 0.0094 | 0.0844 | 0.6135 | 0.8009 | 0.0476 | -0.2986 |
| 55beed6d1a19ce34788b66488ffe3917 | *D_0__Bacteria;D_1__Firmicutes;D_2__Clostridia;D_3__Clostridiales;D_4__Ruminococcaceae;D_5__Acetanaerobacterium;D_6__Acetanaerobacterium elongatum* | 0.0872 | 0.5220 | 0.0276 | 0.1647 | 0.0031 | -0.2936 |
| 4e2e6735331ad01a30c94d2f4c285dc2 | *D_0__Bacteria;D_1__Firmicutes;D_2__Clostridia;D_3__Clostridiales;D_4__Lachnospiraceae;D_5__CAG-56;D_6__uncultured bacterium* | 0.2009 | 0.9831 | 0.0013 | 0.6617 | 0.0011 | -0.2901 |
| a55a010c9525ce2943a553dca1421b1c | *D_0__Bacteria;D_1__Firmicutes;D_2__Clostridia;D_3__Clostridiales;D_4__Lachnospiraceae;D_5__[Eubacterium] ventriosum group;D_6__uncultured bacterium* | 0.0131 | 0.1730 | 0.0888 | 0.2454 | 0.0013 | -0.2851 |
| a891fb44241e433fa8acf251ef4d328f | *D_0__Bacteria;D_1__Firmicutes;D_2__Clostridia;D_3__Clostridiales;D_4__Lachnospiraceae;D_5__Lacrimispora;D_6__Lacrimispora amygdalina* | 0.0338 | 0.0373 | 0.0517 | 0.5168 | 0.0031 | -0.2757 |
| 208c0c43c7c5d4a6bb549e0c8365cd21 | *D_0__Bacteria;D_1__Firmicutes;D_2__Clostridia;D_3__Clostridiales;D_4__Lachnospiraceae;D_5__[Eubacterium] eligens group* | 0.0928 | 0.0668 | 0.2743 | 0.4513 | 0.0371 | -0.2638 |
| 91587a85e342f8dba27f54e15ac0ea77 | *D_0__Bacteria;D_1__Firmicutes;D_2__Clostridia;D_3__Clostridiales;D_4__Lachnospiraceae;D_5__Tyzzerella 3;D_6__unidentified* | 0.0817 | 0.3200 | 0.0044 | 0.0199 | 0.0003 | -0.2637 |
| e335f74033bc634af43ee6baa84fa247 | *D_0__Bacteria;D_1__Firmicutes;D_2__Clostridia;D_3__Clostridiales;D_4__Peptostreptococcaceae;D_5__Romboutsia;D_6__Romboutsia timonensis* | 0.5126 | 0.1468 | 0.0692 | 0.7169 | 0.0109 | -0.2632 |
| c97ea25ba069877eac769cb05968724c | *D_0__Bacteria;D_1__Firmicutes;D_2__Clostridia;D_3__Clostridiales;D_4__Ruminococcaceae;D_5__Ruminococcaceae UCG-014* | 0.0679 | 0.5257 | 0.0086 | 0.0585 | 0.0002 | -0.2593 |
| ee72ee97db064e5d745356f479b9e576 | *D_0__Bacteria;D_1__Firmicutes;D_2__Clostridia;D_3__Clostridiales;D_4__Ruminococcaceae;D_5__Ruminococcus 2* | 0.2759 | 0.2497 | 0.0054 | 0.2918 | 0.0141 | -0.2508 |
| df16a09f3e448ddd3b1ec54066d3081d | *D_0__Bacteria;D_1__Firmicutes;D_2__Clostridia;D_3__Clostridiales;D_4__Ruminococcaceae;D_5__Butyricicoccus;D_6__Butyricicoccus faecihominis* | 0.0633 | 0.2876 | 0.0184 | 0.0362 | 0.0101 | -0.2426 |
| aff3962bbd7607a2971161d9840ca409 | *D_0__Bacteria;D_1__Firmicutes;D_2__Clostridia;D_3__Clostridiales;D_4__Ruminococcaceae;D_5__Ruminiclostridium 5;D_6__Ruminococcus bromii* | 0.1263 | 0.5902 | 0.0065 | 0.3108 | 0.0005 | -0.2412 |
| fd496fd32dc8c08ade2e8b6c9d8ee13d | *D_0__Bacteria;D_1__Firmicutes;D_2__Bacilli;D_3__Lactobacillales;D_4__Streptococcaceae;D_5__Streptococcus;D_6__Streptococcus thermophilus TH1435* | 0.0183 | 0.3617 | 0.0059 | 0.2519 | 0.0025 | -0.2402 |
| f5f5e0da89730462abaf6301a9557193 | *D_0__Bacteria;D_1__Firmicutes;D_2__Clostridia;D_3__Clostridiales;D_4__Ruminococcaceae;D_5__Faecalibacterium;D_6__Faecalibacterium prausnitzii A2-165* | 0.1397 | 0.0835 | 0.0474 | 0.8187 | 0.0052 | -0.2400 |
| 86ff0efd7fd88b82809500c90c4a7832 | *D_0__Bacteria;D_1__Firmicutes;D_2__Clostridia;D_3__Clostridiales;D_4__Ruminococcaceae;D_5__Faecalibacterium;D_6__Faecalibacterium prausnitzii* | 0.1487 | 0.1308 | 0.0726 | 0.4572 | 0.0232 | -0.2302 |
| 33518e48b174ac428cb7be1a216c6c6e | *D_0__Bacteria;D_1__Firmicutes;D_2__Clostridia;D_3__Clostridiales;D_4__Lachnospiraceae;D_5__Anaerostipes;D_6__Anaerostipes hadrus* | 0.2826 | 0.1361 | 0.1012 | 0.7562 | 0.0372 | -0.2186 |
| c66ec67719e54d2cb0afc3154b3c23af | *D_0__Bacteria;D_1__Firmicutes;D_2__Clostridia;D_3__Clostridiales;D_4__Christensenellaceae;D_5__Christensenellaceae R-7 group* | 0.1579 | 0.4934 | 0.0069 | 0.8687 | 0.0097 | -0.2092 |
| eb51a6482b09f0c5962e362d2d23abb9 | *D_0__Bacteria;D_1__Firmicutes;D_2__Clostridia;D_3__Clostridiales;D_4__Ruminococcaceae;D_5__Ruminococcaceae UCG-014* | 0.2129 | 0.0263 | 0.0295 | 0.1865 | 0.0024 | -0.2042 |
| aae4c4ed528ae16ec417df816699cfdb | *D_0__Bacteria;D_1__Firmicutes;D_2__Clostridia;D_3__Clostridiales;D_4__Ruminococcaceae;D_5__Ruminococcaceae UCG-005* | 0.6500 | 0.7833 | 0.0073 | 0.3822 | 0.0085 | -0.1996 |
| 15b5966398a85e5c75501ad243691dfc | *D_0__Bacteria;D_1__Firmicutes;D_2__Clostridia;D_3__Clostridiales;D_4__Lachnospiraceae;D_5__Lachnoclostridium* | 0.1048 | 0.5475 | 0.1229 | 0.0999 | 0.0173 | -0.1866 |
| 96040114b5274bdb987f28c63b6b6b87 | *D_0__Bacteria;D_1__Firmicutes;D_2__Clostridia;D_3__Clostridiales;D_4__Lachnospiraceae;D_5_Roseburia;D_6__Roseburia intestinalis* | 0.0447 | NA | 0.0610 | 0.2276 | 0.0024 | -0.1772 |
| 54a2bfb0a96adc52c4ccca239ad01466 | *D_0__Bacteria;D_1__Firmicutes;D_2__Clostridia;D_3__Clostridiales;D_4__Ruminococcaceae;D_5__Ruminiclostridium 6;D_6__Ruminiclostridium hungatei* | 0.0950 | 0.8319 | 0.0848 | 0.5961 | 0.0350 | -0.1591 |
| 6c324279c1fc2965e594486ac26b0548 | *D_0__Bacteria;D_1__Firmicutes;D_2__Clostridia;D_3__Clostridiales;D_4__Lachnospiraceae;D_5__Coprococcus 2* | 0.4817 | 0.2162 | 0.0140 | 0.9547 | 0.0092 | -0.1551 |
| 43202fe954709a694e31c3a1be7e889b | *D_0__Bacteria;D_1__Firmicutes;D_2__Clostridia;D_3__Clostridiales;D_4__Ruminococcaceae;D_5__Ruminococcaceae UCG-014* | 0.1633 | 0.2458 | 0.3508 | 0.5316 | 0.0437 | -0.1502 |
| cd9401a6bce4a63af516d06d2a843f9d | *D_0__Bacteria;D_1__Firmicutes;D_2__Negativicutes;D_3__Selenomonadales;D_4__Veillonellaceae;D_5__Veillonella;D_6__Veillonella parvula DSM 2008* | 0.1477 | 0.8390 | 0.0296 | 0.6852 | 0.0394 | -0.1406 |
| fb428d07d03e32be0b85eacbed10df0a | *D_0__Bacteria;D_1__Firmicutes;D_2__Clostridia;D_3__Clostridiales;D_4__Lachnospiraceae;D_5__Coprococcus 2;D_6__Coprococcus eutactus* | 0.4022 | 0.4935 | 0.0045 | 1.0000 | 0.0434 | -0.1191 |
| 369bbf86b13d5924c095aad30bba9af1 | *D_0__Bacteria;D_1__Firmicutes;D_2__Clostridia;D_3__Clostridiales;D_4__Lachnospiraceae;D_5__Blautia;D_6__Blautia schinkii* | 0.0357 | 0.1022 | 0.2247 | 0.5440 | 0.0030 | -0.1155 |
| c0044f57ec04cd70905d690c5b7bd147 | *D_0__Bacteria;D_1__Firmicutes;D_2__Clostridia;D_3__Clostridiales;D_4__Lachnospiraceae;D_5__Coprococcus 2* | 0.6056 | 0.0414 | 0.0344 | 0.6771 | 0.0244 | -0.1089 |
| 4924bfca291797ea1fca3e384f4143f9 | *D_0__Bacteria;D_1__Firmicutes;D_2__Clostridia;D_3__Clostridiales;D_4__Lachnospiraceae;D_6__Roseburia hominis A2-183* | 0.0121 | NA | 0.5961 | 0.1608 | 0.0145 | -0.0909 |
| 366bb3ddd63d135bec172051cf2cfe0a | *D_0__Bacteria;D_1__Actinobacteria;D_2__Actinobacteria;D_3__Actinomycetales;D_4__Actinomycetaceae;D_5__Actinomyces;D_6__Actinomyces graevenitzii F0530* | NA | 0.9172 | 0.1096 | 0.0464 | 0.0393 | -0.0881 |
| 25bf7b31746631bbb8582ebe892fe837 | *D_0__Bacteria;D_1__Firmicutes;D_2__Clostridia;D_3__Clostridiales;D_4__Ruminococcaceae;D_5__Ruminococcaceae UCG-014* | 0.1199 | NA | 0.0155 | 0.3384 | 0.0055 | -0.0760 |
| 0b53415990b1e4435ba981cc617f6be0 | *D_0__Bacteria;D_1__Firmicutes;D_2__Clostridia;D_3__Clostridiales;D_4__Lachnospiraceae;D_5__Lachnospiraceae UCG-003;D_6__uncultured bacterium* | 0.1248 | NA | 0.0690 | 0.1608 | 0.0072 | -0.0690 |
| cd5b7f7211526a4109099a6fd4b01aab | *D_0__Bacteria;D_1__Firmicutes;D_2__Clostridia;D_3__Clostridiales;D_4__Defluviitaleaceae;D_5__Defluviitaleaceae UCG-011;D_6__uncultured bacterium* | 0.1166 | NA | 0.2617 | 0.3337 | 0.0323 | -0.0594 |
| 54886b8e1a829f2bf0f874ea7a002750 | *D_0__Bacteria;D_1__Firmicutes;D_2__Clostridia;D_3__Clostridiales;D_4__Lachnospiraceae;D_5__Acetitomaculum;D_6__Firmicutes bacterium CAG_194_44_15* | 0.7221 | NA | 0.0262 | 0.0815 | 0.0106 | -0.0349 |
| 4e0a8ac6434b87de6ed53d016fd7c87c | *D_0__Bacteria;D_1__Firmicutes;D_2__Clostridia;D_3__Clostridiales;D_4__Family XIII;D_5__Family XIII AD3011 group* | 0.8732 | 0.2261 | 0.0229 | 0.2892 | 0.0123 | -0.0327 |
| ee5b57a629da222ad236b5d4bc329d71 | *D_0__Bacteria;D_1__Firmicutes;D_2__Erysipelotrichia;D_3__Erysipelotrichales;D_4__Erysipelotrichaceae;D_5__Merdibacter;D_6__Merdibacter massiliensis* | 0.7320 | 0.1022 | 0.0732 | 0.0938 | 0.0197 | -0.0292 |
| e35e7d4cf6b6c4837a7bf77ca88c05ca | *D_0__Bacteria;D_1__Firmicutes;D_2__Clostridia;D_3__Clostridiales;D_4__Ruminococcaceae;D_5__Ruminococcaceae UCG-010;D_6__uncultured bacterium* | 0.8734 | NA | 0.0288 | 0.3414 | 0.0255 | -0.0176 |
| 989f2970e8c86c75bf4b205be62fd345 | *D_0__Bacteria;D_1__Firmicutes;D_2__Clostridia;D_3__Clostridiales;D_4__Ruminococcaceae;D_5__Ruminiclostridium 5;D_6__uncultured bacterium* | NA | 0.1022 | 0.1071 | 0.2892 | 0.0227 | -0.0074 |
| 8ee6f70260b29cb40fdff8d4500ae32a | *D_0__Bacteria;D_1__Firmicutes;D_2__Clostridia;D_3__Clostridiales;D_4__Lachnospiraceae;D_5__Roseburia;D_6__Roseburia hominis A2-183* | 0.6258 | NA | 0.0569 | 0.0815 | 0.0335 | -0.0071 |
| 8d5153b41e59c5ffb5a1fc080eb784cf | *D_0__Bacteria;D_1__Firmicutes;D_2__Clostridia;D_3__Clostridiales;D_4__Ruminococcaceae;D_5__Ruminococcaceae UCG-004;D_6__uncultured bacterium* | 0.7784 | 0.2488 | 0.0095 | 0.3384 | 0.0310 | 0.0021 |
| 003e253932cf83d92de80661474011ca | *D_0__Bacteria;D_1__Firmicutes;D_2__Clostridia;D_3__Clostridiales;D_4__Christensenellaceae;D_5__Catabacter;D_6__Christensenella massiliensis* | 0.5345 | NA | 0.0045 | 0.3722 | 0.0392 | 0.0040 |
| 1d49df56b83a383d8b998af9d148f52c | *D_0__Bacteria;D_1__Firmicutes;D_2__Clostridia;D_3__Clostridiales;D_4__Clostridiaceae 1;D_5__Clostridium sensu stricto 1* | 0.7254 | NA | 0.0963 | 0.0938 | 0.0367 | 0.0215 |
| 0c410683e68edf42337559941e5d3018 | *D_0__Bacteria;D_1__Firmicutes;D_2__Clostridia;D_3__Clostridiales;D_4__Ruminococcaceae;D_5__Ruminococcaceae UCG-013* | 0.1239 | NA | 0.0253 | 0.6426 | 0.0121 | 0.0231 |
| b5a69fb7a66fdb2bf437bd286b439706 | *D_0__Bacteria;D_1__Bacteroidetes;D_2__Bacteroidia;D_3__Bacteroidales;D_4__Rikenellaceae;D_5__Alistipes;D_6__Alistipes shahii WAL 8301* | NA | 0.9886 | 0.0131 | 0.4681 | 0.0473 | 0.0274 |
| bc7f6c40d5f8e134763db884ca78cb1d | *D_0__Bacteria;D_1__Firmicutes;D_2__Clostridia;D_3__Clostridiales;D_4__Lachnospiraceae;D_5__Blautia;D_6__Blautia producta ATCC 27340* | 0.1626 | 0.1359 | 0.3323 | NA | 0.0329 | 0.0338 |
| 9ee9d7f1e7a96963f1ee2668f858fced | *D_0__Bacteria;D_1__Firmicutes;D_2__Clostridia;D_3__Clostridiales;D_4__Family XI;D_5__Anaerococcus;D_6__Anaerococcus vaginalis ATCC 51170* | 0.4516 | NA | 0.0073 | 0.6051 | 0.0093 | 0.0353 |
| 868be84b9daccd25a648f872c8d5e9ab | *D_0__Bacteria;D_1__Firmicutes;D_2__Negativicutes;D_3__Selenomonadales;D_4__Veillonellaceae;D_5__uncultured;D_6__uncultured organism* | NA | 0.1671 | 0.1375 | 0.0419 | 0.0116 | 0.0418 |
| 12cc4c8a5d46fec0b6738bfe7c6c8a08 | *D_0__Bacteria;D_1__Bacteroidetes;D_2__Bacteroidia;D_3__Bacteroidales;D_4__Bacteroidaceae;D_5__Bacteroides;D_6__Bacteroides intestinalis* | 0.1022 | NA | 0.0157 | 0.6547 | 0.0082 | 0.0518 |
| 25394a604bbcd078c80186b300dab8ee | *D_0__Bacteria;D_1__Firmicutes;D_2__Erysipelotrichia;D_3__Erysipelotrichales;D_4__Erysipelotrichaceae;D_5__Holdemanella;D_6__Holdemanella biformis* | 0.2613 | 0.3238 | 0.0095 | NA | 0.0070 | 0.0749 |
| 8e2daae0f076acb2d186378cb883210a | *D_0__Bacteria;D_1__Firmicutes;D_2__Clostridia;D_3__Clostridiales;D_4__Ruminococcaceae;D_5__UBA1819* | 0.7556 | NA | 0.0041 | 0.1365 | 0.0055 | 0.0750 |
| 5e581b06024ab980e25ee550e0b97247 | *D_0__Bacteria;D_1__Firmicutes;D_2__Clostridia;D_3__Clostridiales;D_4__Christensenellaceae;D_5__Catabacter;D_6__Catabacter hongkongensis* | 0.1239 | 0.6684 | 0.2129 | 0.0399 | 0.0158 | 0.0762 |
| 6ba98911efd5b1fa1cbba9f869562cce | *D_0__Bacteria;D_1__Synergistetes;D_2__Synergistia;D_3__Synergistales;D_4__Synergistaceae;D_5__Cloacibacillus;D_6__Cloacibacillus evryensis* | NA | 0.3321 | 0.0482 | 0.3351 | 0.0103 | 0.0848 |
| 33ba975cdb88b5073a2355ffdf01e9cb | *D_0__Bacteria;D_1__Actinobacteria;D_2__Actinobacteria;D_3__Bifidobacteriales;D_4__Bifidobacteriaceae;D_5__Bifidobacterium;D_6__Bifidobacterium dentium* | 0.0613 | 0.4219 | 0.0152 | 0.6946 | 0.0139 | 0.0890 |
| d1a75593a480570277f05ec729b865de | *D_0__Bacteria;D_1__Firmicutes;D_2__Clostridia;D_3__Clostridiales;D_4__Lachnospiraceae;D_5__[Ruminococcus] gnavus group* | 0.3459 | 0.1444 | NA | 0.0845 | 0.0130 | 0.0907 |
| 19da890972569b6859c360e9b4c8751e | *D_0__Bacteria;D_1__Bacteroidetes;D_2__Bacteroidia;D_3__Bacteroidales;D_4__Porphyromonadaceae;D_5__Porphyromonas;D_6__Porphyromonas sp. HMSC077F02* | 0.0045 | NA | 0.0000 | 0.0110 | 0.0000 | 0.0995 |
| 08bd43071a06abf56df235fca6655bea | *D_0__Bacteria;D_1__Firmicutes;D_2__Clostridia;D_3__Clostridiales;D_4__Lachnospiraceae;D_5__[Ruminococcus] torques group* | 0.2999 | 0.8424 | 0.0372 | 0.0689 | 0.0076 | 0.1033 |
| 24cccadb274fef1f88113e8f019e2a63 | *D_0__Bacteria;D_1__Firmicutes;D_2__Clostridia;D_3__Clostridiales;D_4__Family XIII;D_5__[Eubacterium] nodatum group* | 1.0000 | 0.0096 | 0.0396 | 0.0815 | 0.0018 | 0.1091 |
| edd418053a744b62aa296a5a85039b3a | *D_0__Bacteria;D_1__Firmicutes;D_2__Clostridia;D_3__Clostridiales;D_4__Ruminococcaceae;D_5__Hydrogenoanaerobacterium;D_6__[Clostridium] methylpentosum* | 0.0806 | NA | 0.0116 | 0.4592 | 0.0016 | 0.1095 |
| 9fec7bdd6bd88e710bd69b15692e54a0 | *D_0__Bacteria;D_1__Firmicutes;D_2__Bacilli;D_3__Lactobacillales;D_4__Streptococcaceae;D_5__Streptococcus;D_6__Streptococcus infantarius* | 0.5769 | 0.0255 | 0.0667 | NA | 0.0155 | 0.1143 |
| 7ee3e4343242684ba3ce459672348ff7 | *D_0__Bacteria;D_1__Bacteroidetes;D_2__Bacteroidia;D_3__Bacteroidales;D_4__Bacteroidaceae;D_5__Bacteroides;D_6__Bacteroides nordii* | 0.0689 | 0.9413 | 0.1276 | 0.9818 | 0.0262 | 0.1225 |
| ef1243f7818d1e35c1e968e8539173cf | *D_0__Bacteria;D_1__Firmicutes;D_2__Clostridia;D_3__Clostridiales;D_4__Ruminococcaceae;D_5__Anaerotruncus;D_6__Anaerotruncus sp. AT3* | 0.1118 | 0.1515 | 0.3082 | 0.0938 | 0.0126 | 0.1295 |
| 11b7d5a0dd0446562be236a4a9144c75 | *D_0__Bacteria;D_1__Firmicutes;D_2__Clostridia;D_3__Clostridiales;D_4__Ruminococcaceae;D_5__Ruminiclostridium 9;D_6__uncultured Flavonifractor sp.* | 0.8353 | 0.9631 | 0.0105 | 0.2027 | 0.0096 | 0.1296 |
| ab149525cad479f6599c7923acc49691 | *D_0__Bacteria;D_1__Firmicutes;D_2__Clostridia;D_3__Clostridiales;D_4__Lachnospiraceae;D_5__Lachnospiraceae UCG-010;D_6__uncultured organism* | 0.0654 | 0.3238 | 0.1073 | 0.2592 | 0.0017 | 0.1346 |
| c532c2c02cdb1cec71b6500c5d6c2edf | *D_0__Bacteria;D_1__Bacteroidetes;D_2__Bacteroidia;D_3__Bacteroidales;D_4__Porphyromonadaceae;D_5__Porphyromonas;D_6__Porphyromonas asaccharolytica* | 0.0153 | NA | 0.0001 | 0.1023 | 0.0000 | 0.1377 |
| d515ae0a0b58aa13869a5966a80c98a9 | *D_0__Bacteria;D_1__Firmicutes;D_2__Clostridia;D_3__Clostridiales;D_4__Lachnospiraceae* | 0.1642 | 0.1024 | 0.1291 | 0.7129 | 0.0172 | 0.1448 |
| 31f567f59096ed2d57feecc6c3621a58 | *D_0__Bacteria;D_1__Bacteroidetes;D_2__Bacteroidia;D_3__Bacteroidales;D_4__Porphyromonadaceae;D_5__Porphyromonas;D_6__Porphyromonas sp. 2007b* | 0.1516 | NA | 0.0000 | 0.0098 | 0.0000 | 0.1489 |
| c84d13308b8608a006759b82fcb5e4ac | *D_0__Bacteria;D_1__Firmicutes;D_2__Clostridia;D_3__Clostridiales;D_4__Lachnospiraceae;D_5__Blautia;D_6__Blautia hydrogenotrophica* | 0.9597 | 0.1686 | 0.0382 | 0.8162 | 0.0454 | 0.1508 |
| 7a6dba47692f821cabeaf266b6b36593 | *D_0__Bacteria;D_1__Firmicutes;D_2__Negativicutes;D_3__Selenomonadales;D_4__Veillonellaceae;D_5__Dialister;D_6__Dialister pneumosintes* | 0.0004 | NA | 0.0000 | 0.0425 | 0.0000 | 0.1558 |
| 4bdbaae586dedec92168f93266de0979 | *D_0__Bacteria;D_1__Firmicutes;D_2__Clostridia;D_3__Clostridiales;D_4__Christensenellaceae;D_5__Christensenellaceae R-7 group* | 0.0134 | NA | 0.2023 | 0.4022 | 0.0116 | 0.1581 |
| e502ebea2a54a6f839d310d6d57e7d47 | *D_0__Bacteria;D_1__Fusobacteria;D_2__Fusobacteriia;D_3__Fusobacteriales;D_4__Fusobacteriaceae;D_5__Fusobacterium;D_6__Fusobacterium nucleatum subsp.animalis* | 0.0024 | 0.6684 | 0.0001 | 0.0221 | 0.0000 | 0.1614 |
| 16530e5b50b780018210dac05a471715 | *D_0__Bacteria;D_1__Firmicutes;D_2__Clostridia;D_3__Clostridiales;D_4__Ruminococcaceae;D_5__Ruminococcaceae UCG-002* | 0.1399 | NA | 0.1153 | 0.6256 | 0.0411 | 0.1618 |
| 8646bffa210e3f061cc57f1b1cf2f751 | *D_0__Bacteria;D_1__Firmicutes;D_2__Clostridia;D_3__Clostridiales;D_4__Lachnospiraceae;D_5__Hungatella;D_6__Hungatella hathewayi WAL-18680* | 0.0719 | 0.2855 | 0.0875 | 0.3233 | 0.0056 | 0.1626 |
| 0df6c802966e8670279671824da4f10a | *D_0__Bacteria;D_1__Firmicutes;D_2__Bacilli;D_3__Lactobacillales;D_4__Lactobacillaceae;D_5__Lactobacillus;D_6__Lactobacillus gasseri ATCC 33323* | 0.9165 | 0.7166 | 0.0347 | 0.0231 | 0.0290 | 0.1706 |
| 6b5c5532185593c41966b2173ea9df1c | *D_0__Bacteria;D_1__Firmicutes;D_2__Clostridia;D_3__Clostridiales;D_4__Lachnospiraceae;D_5__Enterocloster;D_6__Enterocloster lavalensis* | 0.0157 | 0.0882 | 0.7197 | 0.6692 | 0.0091 | 0.1884 |
| 07699fa14cb8970eee5e8dda19dd5dbe | *D_0__Bacteria;D_1__Firmicutes;D_2__Clostridia;D_3__Clostridiales;D_4__Lachnospiraceae;D_5__Eisenbergiella;D_6__Eisenbergiella massiliensis* | 0.2559 | 0.0178 | 0.1631 | 0.2541 | 0.0337 | 0.1924 |
| 05dbbf0301a9625e1f1a41470a92f7fe | *D_0__Bacteria;D_1__Firmicutes;D_2__Clostridia;D_3__Clostridiales;D_4__Lachnospiraceae;D_5__Anaerostipes;D_6__Anaerostipes caccae* | 0.1425 | 0.0782 | 0.5640 | 0.0790 | 0.0203 | 0.2010 |
| f479c23321346918723839e33a3544f4 | *D_0__Bacteria;D_1__Firmicutes;D_2__Clostridia;D_3__Clostridiales;D_4__Ruminococcaceae;D_5__UBA1819* | 0.1407 | 0.3150 | 0.0267 | 0.3320 | 0.0032 | 0.2077 |
| 4912cd4c1f9a773f88a73c258c25c825 | *D_0__Bacteria;D_1__Firmicutes;D_2__Clostridia;D_3__Clostridiales;D_4__Lachnospiraceae;D_5__[Eubacterium] fissicatena group* | 0.1066 | 0.0385 | 0.3971 | 0.1789 | 0.0093 | 0.2200 |
| 58bd181f757283f5a4ceebdf90ff6243 | *D_0__Bacteria;D_1__Firmicutes;D_2__Clostridia;D_3__Clostridiales;D_4__Lachnospiraceae;D_5__Sellimonas;D_6__Sellimonas intestinalis* | 0.1587 | 0.1456 | 0.0809 | 0.7090 | 0.0116 | 0.2340 |
| da8644094f1d480f3b3a6a03a7027ab4 | *D_0__Bacteria;D_1__Firmicutes;D_2__Clostridia;D_3__Clostridiales;D_4__Lachnospiraceae;D_5__Hungatella;D_6__Hungatella effluvii* | 0.0199 | 0.0603 | 0.0199 | 0.7289 | 0.0009 | 0.2441 |
| a248d70e41d75f4773ef7c94e831a21a | *D_0__Bacteria;D_1__Firmicutes;D_2__Clostridia;D_3__Clostridiales;D_4__Lachnospiraceae;D_5__Faecalimonas;D_6__Faecalimonas umbilicata* | NA | 0.0804 | 0.1817 | 0.3722 | 0.0396 | 0.2506 |
| 47f770032218f91b352999ff1f663807 | *D_0__Bacteria;D_1__Firmicutes;D_2__Clostridia;D_3__Clostridiales;D_4__Lachnospiraceae;D_5__Lachnoclostridium;D_6__Lachnoclostridium pacaense* | 0.1776 | 0.0895 | 0.0110 | 0.0191 | 0.0002 | 0.2705 |
| e93b313352019b8113e7fc66bf757917 | *D_0__Bacteria;D_1__Firmicutes;D_2__Clostridia;D_3__Clostridiales;D_4__Family XI;D_5__Parvimonas;;D_6__Parvimonas micra* | 0.0019 | NA | 0.0000 | 0.0200 | 0.0000 | 0.2841 |
| 52573485a4521e007e4c5217fc5d7c68 | *D_0__Bacteria;D_1__Firmicutes;D_2__Clostridia;D_3__Clostridiales;D_4__Lachnospiraceae;D_5__Anaerotignum;D_6__Anaerotignum aminivorans* | 0.5249 | 0.1756 | 0.3043 | 0.0007 | 0.0064 | 0.2846 |
| 1f0532e025d966a685fd0d1f3144bc8a | *D_0__Bacteria;D_1__Firmicutes;D_2__Clostridia;D_3__Clostridiales;D_4__Peptostreptococcaceae;D_5__Peptostreptococcus;D_6__Peptostreptococcus stomatis* | 0.0000 | NA | 0.0000 | 1.0000 | 0.0000 | 0.2851 |
| 5fbc483f79cd97d3a7253931f37c33e8 | *D_0__Bacteria;D_1__Firmicutes;D_2__Clostridia;D_3__Clostridiales;D_4__Ruminococcaceae;D_5__Flavonifractor;D_6__Flavonifractor plautii* | 0.0959 | 0.1875 | 0.0904 | 0.2382 | 0.0089 | 0.3024 |
| 7af6fc12809fdafab918881f836460f8 | *D_0__Bacteria;D_1__Firmicutes;D_2__Clostridia;D_3__Clostridiales;D_4__Lachnospiraceae;D_5__Tyzzerella 4* | 0.4706 | 0.1524 | 0.0062 | 0.0805 | 0.0019 | 0.3191 |
| a663a01aeb4dbb88bbe0a9840e8f64dc | *D_0__Bacteria;D_1__Firmicutes;D_2__Erysipelotrichia;D_3__Erysipelotrichales;D_4__Erysipelotrichaceae;D_5__Erysipelatoclostridium;D_6__Erysipelatoclostridium ramosum* | 0.0965 | 0.0051 | 0.2120 | 0.9691 | 0.0146 | 0.3243 |
| 8538d07b8c1e8676f9b9c8c242aab14a | *D_0__Bacteria;D_1__Firmicutes;D_2__Clostridia;D_3__Clostridiales;D_4__Ruminococcaceae;D_5__[Eubacterium] coprostanoligenes group;D_6__unidentified* | 0.0420 | 0.1065 | 0.0016 | 0.7169 | 0.0002 | 0.3310 |
| d49783e7800974b5f2bcce554e26072e | *D_0__Bacteria;D_1__Firmicutes;D_2__Clostridia;D_3__Clostridiales;D_4__Lachnospiraceae;D_5__Blautia;D_6__Blautia sp.* | 0.9080 | 0.0096 | 0.3312 | 0.2293 | 0.0252 | 0.3666 |
| d46e2205f0c6ecf67b51f83d111c509c | *D_0__Bacteria;D_1__Proteobacteria;D_2__Gammaproteobacteria;D_3__Enterobacteriales;D_4__Enterobacteriaceae;D_5__Escherichia-Shigella* | 0.0187 | 0.0072 | 0.5588 | 0.9705 | 0.0080 | 0.4050 |
| 5149bdbf03de13bb38557d48d5141b87 | *D_0__Bacteria;D_1__Firmicutes;D_2__Clostridia;D_3__Clostridiales;D_4__Lachnospiraceae;D_5__Lachnoclostridium;D_6__[Clostridium] scindens* | 0.5878 | 0.0001 | 0.0147 | 0.1532 | 0.0000 | 0.4902 |
| c24e0e391aa836b5eae25567c7eb89ee | *D_0__Bacteria;D_1__Firmicutes;D_2__Clostridia;D_3__Clostridiales;D_4__Lachnospiraceae;D_5__[Ruminococcus] gnavus group* | 0.5567 | 0.0084 | 0.0109 | 0.1368 | 0.0014 | 0.5355 |
| 67e8859e108281ee7971084bfa759522 | *D_0__Bacteria;D_1__Firmicutes;D_2__Clostridia;D_3__Clostridiales;D_4__Lachnospiraceae;D_5__Lachnoclostridium* | 0.0007 | 0.0311 | 0.0028 | 0.0002 | 0.0000 | 0.5539 |
| c67445ed68d61f49e41b7bdd1de019fb | *D_0__Bacteria;D_1__Firmicutes;D_2__Clostridia;D_3__Clostridiales;D_4__Lachnospiraceae;D_5__Lachnoclostridium* | 0.1276 | 0.0057 | 0.0062 | 0.0002 | 0.0000 | 0.6710 |

*** The ID of each amplicon sequence variant (ASV)**

**† Taxonomic information of each ASV**

**‡ Single-study P-value calculated by a two-sided Wilcoxon test**

**# Meta-analysis P-value calculated by a two-sided blocked Wilcoxon test (n=524 independent observations)**

**$ Mean of generalized fold changes across studies, GFOLD-meta >0: ASVs enriched in cancer compared with adenoma; <0: ASVs enriched in adenoma compared with cancer**

**Supplementary Table 3 | Gut microbiome-based markers discriminated between control and adenoma**

| **Important features*** | **Taxon^†^** | **Feature Ranks^#^** | **Biomarker Ranks^$^** |
| --- | --- | --- | --- |
| Age | */* | 1 | / |
| BMI | */* | 2 | / |
| 94928715987dcb9638dd5c0e8f9b20f4 | *D_0__Bacteria;D_1__Firmicutes;D_2__Clostridia;D_3__Clostridiales;D_4__Christensenellaceae;D_5__Christensenellaceae R-7 group* | 3 | 1 |
| aae4c4ed528ae16ec417df816699cfdb | *D_0__Bacteria;D_1__Firmicutes;D_2__Clostridia;D_3__Clostridiales;D_4__Ruminococcaceae;D_5__Ruminococcaceae UCG-005* | 4 | 2 |
| 410e1eaa1468a3d40595898f671ba0b3 | *D_0__Bacteria;D_1__Firmicutes;D_2__Clostridia;D_3__Clostridiales;D_4__Ruminococcaceae;D_5__[Eubacterium] coprostanoligenes group;D_6__Eubacterium coprostanoligenes* | 5 | 3 |
| 2740cf2417c92847cc298cbd71dd1fcd | *D_0__Bacteria;D_1__Firmicutes;D_2__Negativicutes;D_3__Selenomonadales;D_4__Veillonellaceae;D_5__Veillonella;D_6__Veillonella parvula* | 6 | 4 |
| f50508546ae13143015f8c4cb976d0e4 | *D_0__Bacteria;D_1__Firmicutes;D_2__Clostridia;D_3__Clostridiales;D_4__Lachnospiraceae;D_5__[Eubacterium] ruminantium group;D_6__Eubacterium coprostanoligenes.* | 7 | 5 |
| c5ec23b5e73a33133aa0fcf9acd58650 | *D_0__Bacteria;D_1__Firmicutes;D_2__Clostridia;D_3__Clostridiales;D_4__Ruminococcaceae;D_5__Ruminiclostridium 9* | 8 | 6 |
| Gender | */* | 9 | / |
| 6e9ac49ab03e8cf905432ad62df55c30 | *D_0__Bacteria;D_1__Firmicutes;D_2__Clostridia;D_3__Clostridiales;D_4__Family XIII;D_5__Family XIII UCG-001;D_6__Aminipila butyrica* | 10 | 7 |
| d83f60183d81253a505beaeef3cd168f | *D_0__Bacteria;D_1__Actinobacteria;D_2__Actinobacteria;D_3__Micrococcales;D_4__Micrococcaceae;D_5__Rothia;D_6__Rothia dentocariosa* | 11 | 8 |

*** The selceted features that distinguish control from adenoma**

**†Taxonomic information of each ASV**

**# The ranks of important fatures (8ASVs and 3 patient metadata)**

**$ The ranks of biomarkers (8ASVs)**

**Supplementary Table 4 | Gut microbiome-based markers discriminated between adenoma and cancer**

| **Important features*** | **Taxon^†^** | **Feature Ranks^#^** | **Biomarker Ranks^$^** |
| --- | --- | --- | --- |
| BMI | */* | 1 | / |
| fd496fd32dc8c08ade2e8b6c9d8ee13d | *D_0__Bacteria;D_1__Firmicutes;D_2__Bacilli;D_3__Lactobacillales;D_4__Streptococcaceae;D_5__Streptococcus;D_6__Streptococcus thermophilus TH1435* | 2 | 1 |
| Age | */* | 3 | / |
| e93b313352019b8113e7fc66bf757917 | *D_0__Bacteria;D_1__Firmicutes;D_2__Clostridia;D_3__Clostridiales;D_4__Family XI;D_5__Parvimonas;D_6__Parvimonas micra* | 4 | 2 |
| fd44d4cb468fd7dc9b3227867714ed87 | *D_0__Bacteria;D_1__Bacteroidetes;D_2__Bacteroidia;D_3__Bacteroidales;D_4__Bacteroidaceae;D_5__Bacteroides;D_6__Bacteroides dorei* | 5 | 3 |
| 5149bdbf03de13bb38557d48d5141b87 | *D_0__Bacteria;D_1__Firmicutes;D_2__Clostridia;D_3__Clostridiales;D_4__Lachnospiraceae;D_5__Lachnoclostridium;D_6__[Clostridium] scindens* | 6 | 4 |
| e9f900358bb9297e10b9fbb1321b1e6e | *D_0__Bacteria;D_1__Firmicutes;D_2__Erysipelotrichia;D_3__Erysipelotrichales;D_4__Erysipelotrichaceae;D_5__Erysipelatoclostridium;D_6__Erysipelatoclostridium ramosum* | 7 | 5 |
| d49783e7800974b5f2bcce554e26072e | *D_0__Bacteria;D_1__Firmicutes;D_2__Clostridia;D_3__Clostridiales;D_4__Lachnospiraceae;D_5__Blautia;D_6__Blautia sp.* | 8 | 6 |
| 8538d07b8c1e8676f9b9c8c242aab14a | *D_0__Bacteria;D_1__Firmicutes;D_2__Clostridia;D_3__Clostridiales;D_4__Ruminococcaceae;D_5__[Eubacterium] coprostanoligenes group;D_6__unidentified* | 9 | 7 |
| 6ec1023583647a401269990c4e8ad716 | *D_0__Bacteria;D_1__Firmicutes;D_2__Clostridia;D_3__Clostridiales;D_4__Lachnospiraceae;D_5__Lachnospira;D_6__Lachnospira pectinoschiza* | 10 | 8 |
| 16530e5b50b780018210dac05a471715 | *D_0__Bacteria;D_1__Firmicutes;D_2__Clostridia;D_3__Clostridiales;D_4__Ruminococcaceae;D_5__Ruminococcaceae UCG-002* | 11 | 9 |
| aff3962bbd7607a2971161d9840ca409 | *D_0__Bacteria;D_1__Firmicutes;D_2__Clostridia;D_3__Clostridiales;D_4__Ruminococcaceae;D_5__Ruminiclostridium 5;D_6__Ruminococcus bromii* | 12 | 10 |
| 19da890972569b6859c360e9b4c8751e | *D_0__Bacteria;D_1__Bacteroidetes;D_2__Bacteroidia;D_3__Bacteroidales;D_4__Porphyromonadaceae;D_5__Porphyromonas;D_6__Porphyromonas sp. HMSC077F02* | 13 | 11 |
| 31f567f59096ed2d57feecc6c3621a58 | *D_0__Bacteria;D_1__Bacteroidetes;D_2__Bacteroidia;D_3__Bacteroidales;D_4__Porphyromonadaceae;D_5__Porphyromonas;D_6__Porphyromonas sp. 2007b* | 14 | 12 |
| f50508546ae13143015f8c4cb976d0e4 | *D_0__Bacteria;D_1__Firmicutes;D_2__Clostridia;D_3__Clostridiales;D_4__Lachnospiraceae;D_5__[Eubacterium] ruminantium group;D_6__Eubacterium ruminantium* | 15 | 13 |
| 91587a85e342f8dba27f54e15ac0ea77 | *D_0__Bacteria;D_1__Firmicutes;D_2__Clostridia;D_3__Clostridiales;D_4__Lachnospiraceae;D_5__Tyzzerella 3;D_6__unidentified* | 16 | 14 |
| 8646bffa210e3f061cc57f1b1cf2f751 | *D_0__Bacteria;D_1__Firmicutes;D_2__Clostridia;D_3__Clostridiales;D_4__Lachnospiraceae;D_5__Hungatella;D_6__Hungatella hathewayi WAL-18680* | 17 | 15 |
| 2de0f958e30d26f04cba8d2a920bbe46 | *D_0__Bacteria;D_1__Firmicutes;D_2__Clostridia;D_3__Clostridiales;D_4__Lachnospiraceae;D_5__Blautia;D_6__Blautia faecis* | 18 | 16 |
| 7ee3e4343242684ba3ce459672348ff7 | *D_0__Bacteria;D_1__Bacteroidetes;D_2__Bacteroidia;D_3__Bacteroidales;D_4__Bacteroidaceae;D_5__Bacteroides;D_6__Bacteroides nordii* | 19 | 17 |
| ab149525cad479f6599c7923acc49691 | *D_0__Bacteria;D_1__Firmicutes;D_2__Clostridia;D_3__Clostridiales;D_4__Lachnospiraceae;D_5__Lachnospiraceae UCG-010;D_6__uncultured organism* | 20 | 18 |
| a55a010c9525ce2943a553dca1421b1c | *D_0__Bacteria;D_1__Firmicutes;D_2__Clostridia;D_3__Clostridiales;D_4__Lachnospiraceae;D_5__[Eubacterium] ventriosum group;D_6__uncultured bacterium* | 21 | 19 |
| 9fec7bdd6bd88e710bd69b15692e54a0 | *D_0__Bacteria;D_1__Firmicutes;D_2__Bacilli;D_3__Lactobacillales;D_4__Streptococcaceae;D_5__Streptococcus;D_6__Streptococcus infantarius* | 22 | 20 |
| 96040114b5274bdb987f28c63b6b6b87 | *D_0__Bacteria;D_1__Firmicutes;D_2__Clostridia;D_3__Clostridiales;D_4__Lachnospiraceae;D_5_Roseburia;D_6__Roseburia intestinalis* | 23 | 21 |
| ee5b57a629da222ad236b5d4bc329d71 | *D_0__Bacteria;D_1__Firmicutes;D_2__Erysipelotrichia;D_3__Erysipelotrichales;D_4__Erysipelotrichaceae;D_5__Merdibacter;D_6__Merdibacter massiliensis* | 24 | 22 |
| d1a75593a480570277f05ec729b865de | *D_0__Bacteria;D_1__Firmicutes;D_2__Clostridia;D_3__Clostridiales;D_4__Lachnospiraceae;D_5__[Ruminococcus] gnavus group* | 25 | 23 |
| 4924bfca291797ea1fca3e384f4143f9 | *D_0__Bacteria;D_1__Firmicutes;D_2__Clostridia;D_3__Clostridiales;D_4__Lachnospiraceae;D_5__Roseburia;D_6__Roseburia hominis A2-183* | 26 | 24 |

*** The selceted features that distinguish adenoma from cancer**

**† Taxonomic information of each ASV**

**# The ranks of important fatures (24ASVs and 2 patient metadata)**

**$ The ranks of biomarkers (24ASVs)**

**Supplementary Table 5 | Biomarkers of two meta-analysis and this study for distinguishing control and CRC.**

| **This study** | **Thomas^#^** | **Zeller*** |
| --- | --- | --- |
| *D_0__Bacteria;D_1__Proteobacteria;D_2__Gammaproteobacteria;D_3__Betaproteobacteriales;D_4__Burkholderiaceae;D_5__Oxalobacteraceae;D_6__Oxalobacter* | *Fusobacterium nucleatum* | *Fusobacterium nucleatum s. animalis* |
| *D_0__Bacteria;D_1__Bacteroidetes;D_2__Bacteroidia;D_3__Bacteroidales;D_4__Bacteroidaceae;D_5__Bacteroides;D_6__Bacteroides dorei* | *Parvimonas micra* | *Parvimonas micra* |
| *D_0__Bacteria;D_1__Fusobacteria;D_2__Fusobacteriia;D_3__Fusobacteriales;D_4__Fusobacteriaceae;D_5__Fusobacterium;D_6__Fusobacterium nucleatum subsp. Animalis* | *Gemella morbillorum* | *Gemella morbillorum* |
| *D_0__Bacteria;D_1__Firmicutes;D_2__Clostridia;D_3__Clostridiales;D_4__Clostridiaceae 1;D_5__Clostridium sensu stricto 1;D_6__unknown Clostridiales* | *Peptostreptococcus stomatis* | *Peptostreptococcus stomatis* |
| *D_0__Bacteria;D_1__Firmicutes;D_2__Clostridia;D_3__Clostridiales;D_4__Ruminococcaceae;D_5__Ruminococcus 1;D_6__Ruminococcus bicirculans* | *Parvimonas spp.* | *unknown Dialister* |
| *D_0__Bacteria;D_1__Firmicutes;D_2__Clostridia;D_3__Clostridiales;D_4__Ruminococcaceae;D_5__Ruminiclostridium 5;D_6__[Clostridium] leptum* | *Solobacterium moorei* | *unknown Porphyromonas* |
| *D_0__Bacteria;D_1__Bacteroidetes;D_2__Bacteroidia;D_3__Bacteroidales;D_4__Porphyromonadaceae;D_5__Porphyromonas;D_6__Porphyromonas asaccharolytica* | *Clostridium symbiosum* | *Solobacterium moorei* |
| *D_0__Bacteria;D_1__Firmicutes;D_2__Clostridia;D_3__Clostridiales;D_4__Christensenellaceae;D_5__Christensenellaceae R-7 group;D_6__uncultured prokaryote* | *Porphyromonas asaccharolytica* | *Clostridium symbiosum* |
| *D_0__Bacteria;D_1__Firmicutes;D_2__Clostridia;D_3__Clostridiales;D_4__Lachnospiraceae;D_5__Roseburia;D_6__Roseburia hominis A2-183* | *Bacteroides fragilis* | *Porphyromonas uenonis* |
| *D_0__Bacteria;D_1__Firmicutes;D_2__Clostridia;D_3__Clostridiales;D_4__Ruminococcaceae;D_5__Subdoligranulum* | *Porphyromonas somerae* | *unknown Clostridiales* |
| *D_0__Bacteria;D_1__Firmicutes;D_2__Clostridia;D_3__Clostridiales;D_4__Ruminococcaceae;D_5__Faecalibacterium；D_6__Faecalibacterium sp. Marseille-P9312* | *Anaerococcus vaginalis* | *Hungatella hathewayi* |
| *D_0__Bacteria;D_1__Firmicutes;D_2__Clostridia;D_3__Clostridiales;D_4__Lachnospiraceae;D_5__GCA-900066575* | *Porphyromonas uenonis* | *Prevotella intermedia* |
| *D_0__Bacteria;D_1__Firmicutes;D_2__Clostridia;D_3__Clostridiales;D_4__Lachnospiraceae;D_5__Hungatella;D_6__Hungatella hathewayi WAL-18680* | *Anaerococcus obesiensis* | *Porphyromonas somerae* |
| *D_0__Bacteria;D_1__Firmicutes;D_2__Clostridia;D_3__Clostridiales;D_4__Ruminococcaceae;D_5__[Eubacterium] coprostanoligenes group;D_6__unidentified* | *Prevotella intermedia* | *Porphyromonas asaccharolytica* |
| *D_0__Bacteria;D_1__Firmicutes;D_2__Clostridia;D_3__Clostridiales;D_4__Christensenellaceae;D_5__Christensenellaceae R-7 group* | *Peptostreptococcus anaerobius* | *Fusobacterium nucleatum s. nucleatum* |
| *D_0__Bacteria;D_1__Firmicutes;D_2__Clostridia;D_3__Clostridiales;D_4__Lachnospiraceae;D_5__Tyzzerella 4;D_6__Tyzzerella nexilis* | *Bifidobacterium catenulatum* | *Parvimonas sp.* |
| *D_0__Bacteria;D_1__Firmicutes;D_2__Negativicutes;D_3__Selenomonadales;D_4__Veillonellaceae;D_5__Dialister;D_6__Dialister pneumosintes* | *Gordonibacter pamelaeae* | *Prevotella nigrescens* |
| *D_0__Bacteria;D_1__Firmicutes;D_2__Clostridia;D_3__Clostridiales;D_4__Lachnospiraceae;D_5__Blautia;D_6__Blautia sp.* | *Streptococcus constellatus* | *unknown Porphyromonas* |
| *D_0__Bacteria;D_1__Firmicutes;D_2__Clostridia;D_3__Clostridiales;D_4__Lachnospiraceae;D_5__Lachnospira;D_6__Lachnospira pectinoschiza* | *Methanobrevibacter smithii* | *Ruminococcus torques* |
| *D_0__Bacteria;D_1__Firmicutes;D_2__Clostridia;D_3__Clostridiales;D_4__Ruminococcaceae;D_5__Faecalibacterium;D_6__Faecalibacterium prausnitzii* | *Roseburia intestinalis* | *Fusobacterium nucleatum s. vincentii* |
| *D_0__Bacteria;D_1__Firmicutes;D_2__Clostridia;D_3__Clostridiales;D_4__Christensenellaceae;D_5__Christensenellaceae R-7 group* | *Granulicatella adiacens* | *Fusobacterium sp. oral taxon 370* |
| *D_0__Bacteria;D_1__Firmicutes;D_2__Clostridia;D_3__Clostridiales;D_4__Lachnospiraceae* | *Peptostreptococcus spp.* | *unknown Peptostreptococcaceae* |
| *D_0__Bacteria;D_1__Firmicutes;D_2__Clostridia;D_3__Clostridiales;D_4__Lachnospiraceae;D_5__Lachnoclostridium;D_6__Enterocloster aldensis* | *Ruminococcus torques* | *Anaerococcus obesiensis/vaginalis* |
| *D_0__Bacteria;D_1__Bacteroidetes;D_2__Bacteroidia;D_3__Bacteroidales;D_4__Prevotellaceae;D_5__Prevotella 7;D_6__Massiliprevotella massiliensis* | *Faecalibacterium prausnitzii* | *unknown Anaerotruncus* |
| *D_0__Bacteria;D_1__Proteobacteria;D_2__Gammaproteobacteria;D_3__Pasteurellales;D_4__Pasteurellaceae;D_5__Haemophilus;D_6__Haemophilus parainfluenzae* | *Eikenella corrodens* | *Porphyromonas uenonis* |
| *D_0__Bacteria;D_1__Firmicutes;D_2__Clostridia;D_3__Clostridiales;D_4__Peptococcaceae;D_5__uncultured* | *Methanobrevibacter spp.* | *unknown Clostridiales* |
| *D_0__Bacteria;D_1__Firmicutes;D_2__Clostridia;D_3__Clostridiales;D_4__Lachnospiraceae;D_5__Lachnospira;D_6__uncultured Lachnospira sp.* |  | *unknown Porphyromonas* |
| *D_0__Bacteria;D_1__Firmicutes;D_2__Clostridia;D_3__Clostridiales;D_4__Lachnospiraceae* |  | *Clostridium boltae/clostridioforme* |
| *D_0__Bacteria;D_1__Firmicutes;D_2__Clostridia;D_3__Clostridiales;D_4__Ruminococcaceae;D_5__Ruminococcaceae UCG-014* |  | *Subdoligranulum sp. 4_3_54A2FAA* |
| *D_0__Bacteria;D_1__Firmicutes;D_2__Clostridia;D_3__Clostridiales;D_4__Peptostreptococcaceae;D_5__Peptostreptococcus;D_6__Peptostreptococcus stomatis* |  |  |
| *D_0__Bacteria;D_1__Bacteroidetes;D_2__Bacteroidia;D_3__Bacteroidales;D_4__Bacteroidaceae;D_5__Bacteroides;D_6__Bacteroides dorei* |  |  |
| *D_0__Bacteria;D_1__Bacteroidetes;D_2__Bacteroidia;D_3__Bacteroidales;D_4__Porphyromonadaceae;D_5__Porphyromonas;D_6__Porphyromonas sp. HMSC077F02* |  |  |
| *D_0__Bacteria;D_1__Bacteroidetes;D_2__Bacteroidia;D_3__Bacteroidales;D_4__Bacteroidaceae;D_5__Bacteroides;D_6__Bacteroides sp. HGA0138* |  |  |
| *D_0__Bacteria;D_1__Firmicutes;D_2__Clostridia;D_3__Clostridiales;D_4__Ruminococcaceae;D_5__Ruminococcaceae UCG-013* |  |  |

**# Thomas AM, et al. (2019) Metagenomic analysis of colorectal cancer datasets identifies cross-cohort microbial diagnostic signatures and a link with choline degradation. Nat Med 25(4):667-678.**

*** Wirbel J, et al. (2019) Meta-analysis of fecal metagenomes reveals global microbial signatures that are specific for colorectal cancer. Nat Med 25(4):679-689.**

**Supplementary Table 6 | Node numbers of differential ASVs between control and adenoma**

| **ASV ID*** | **Taxon^†^** | **node^$^** |
| --- | --- | --- |
| 6e9ac49ab03e8cf905432ad62df55c30 | *D_0__Bacteria;D_1__Firmicutes;D_2__Clostridia;D_3__Clostridiales;D_4__Family XIII;D_5__Family XIII UCG-001;D_6__Aminipila butyrica* | *Aminipila butyrica* |
| 94928715987dcb9638dd5c0e8f9b20f4 | *D_0__Bacteria;D_1__Firmicutes;D_2__Clostridia;D_3__Clostridiales;D_4__Christensenellaceae;D_5__Christensenellaceae R-7 group* | *Christensenellaceae R-7 group sp.* |
| 410e1eaa1468a3d40595898f671ba0b3 | *D_0__Bacteria;D_1__Firmicutes;D_2__Clostridia;D_3__Clostridiales;D_4__Ruminococcaceae;D_5__[Eubacterium] coprostanoligenes group;D_6__Eubacterium coprostanoligenes* | *Eubacterium coprostanoligenes* |
| f50508546ae13143015f8c4cb976d0e4 | *D_0__Bacteria;D_1__Firmicutes;D_2__Clostridia;D_3__Clostridiales;D_4__Lachnospiraceae;D_5__[Eubacterium] ruminantium group;D_6__Eubacterium ruminantium* | *Eubacterium ruminantium* |
| d83f60183d81253a505beaeef3cd168f | *D_0__Bacteria;D_1__Actinobacteria;D_2__Actinobacteria;D_3__Micrococcales;D_4__Micrococcaceae;D_5__Rothia;D_6__Rothia dentocariosa* | *Rothia dentocariosa* |
| c5ec23b5e73a33133aa0fcf9acd58650 | *D_0__Bacteria;D_1__Firmicutes;D_2__Clostridia;D_3__Clostridiales;D_4__Ruminococcaceae;D_5__Ruminiclostridium 9* | *Ruminiclostridium 9 sp.* |
| aae4c4ed528ae16ec417df816699cfdb | *D_0__Bacteria;D_1__Firmicutes;D_2__Clostridia;D_3__Clostridiales;D_4__Ruminococcaceae;D_5__Ruminococcaceae UCG-005* | *Ruminococcaceae UCG-005 sp.* |
| 2740cf2417c92847cc298cbd71dd1fcd | *D_0__Bacteria;D_1__Firmicutes;D_2__Negativicutes;D_3__Selenomonadales;D_4__Veillonellaceae;D_5__Veillonella;D_6__Veillonella parvula* | *Veillonella parvula* |
| 35ffcc3b809d667286737d79670b8de5 | *D_0__Bacteria;D_1__Actinobacteria;D_2__Actinobacteria;D_3__Bifidobacteriales;D_4__Bifidobacteriaceae;D_5__Bifidobacterium;D_6__Bifidobacterium longum* | 1 |
| 33518e48b174ac428cb7be1a216c6c6e | *D_0__Bacteria;D_1__Firmicutes;D_2__Clostridia;D_3__Clostridiales;D_4__Lachnospiraceae;D_5__Anaerostipes;D_6__Anaerostipes hadrus* | 2 |
| d114fb4c335125128be28401522dd41a | *D_0__Bacteria;D_1__Firmicutes;D_2__Bacilli;D_3__Lactobacillales;D_4__Streptococcaceae;D_5__Lactococcus;D_6__Lactococcus taiwanensis* | 3 |
| a7e28b3f4879359a929a0da8c8557757 | *D_0__Bacteria;D_1__Firmicutes;D_2__Clostridia;D_3__Clostridiales;D_4__Clostridiaceae;D_5__Clostridiaceae;D_6__Clostridiaceae* | 4 |
| 868be84b9daccd25a648f872c8d5e9ab | *D_0__Bacteria;D_1__Firmicutes;D_2__Negativicutes;D_3__Selenomonadales;D_4__Veillonellaceae;D_5__uncultured;D_6__uncultured organism* | 5 |
| 1eaa3cb7baaac3220b0606e3621afd16 | *D_0__Bacteria;D_1__Bacteroidetes;D_2__Bacteroidia;D_3__Bacteroidales;D_4__Rikenellaceae;D_5__Alistipes;D_6__Faecalibacterium prausnitzii* | 6 |
| da1d26d90a7443d34675778207d2c227 | *D_0__Bacteria;D_1__Firmicutes;D_2__Clostridia;D_3__Clostridiales;D_4__Ruminococcaceae;D_5__Ruminiclostridium 9* | 7 |
| e93b313352019b8113e7fc66bf757917 | *D_0__Bacteria;D_1__Firmicutes;D_2__Clostridia;D_3__Clostridiales;D_4__Family XI;D_5__Parvimonas;D_6__Parvimonas micra* | 8 |
| e4f195476bffa394e3048063f99360ea | *D_0__Bacteria;D_1__Bacteroidetes;D_2__Bacteroidia;D_3__Bacteroidales;D_4__Barnesiellaceae;D_5__Coprobacter;D_6__Coprobacter fastidiosus NSB1* | 9 |
| b204a17b1a7f29074d81303d5ca68464 | *D_0__Bacteria;D_1__Firmicutes;D_2__Clostridia;D_3__Clostridiales;D_4__Ruminococcaceae;D_5__Phocea;D_6__Phocea massiliensis* | 10 |
| 8ca0b9917d926bb6241ce6e23673fce4 | *D_0__Bacteria;D_1__Firmicutes;D_2__Clostridia;D_3__Clostridiales;D_4__Lachnospiraceae;D_5__Blautia;D_6__Blautia schinkii* | 11 |
| 74792adac452eb3544cdc17064d6bbee | *D_0__Bacteria;D_1__Proteobacteria;D_2__Deltaproteobacteria;D_3__Desulfovibrionales;D_4__Desulfovibrionaceae;D_5__Desulfovibrio;D_6__Desulfovibrio piger* | 12 |
| 2ef1e51ab1cf99a3c6417b05a060830e | *D_0__Bacteria;D_1__Firmicutes;D_2__Bacilli;D_3__Lactobacillales;D_4__Lactobacillaceae;D_5__Lactobacillus;D_6__Lactobacillus caviae* | 13 |
| ad03570a6623c3fe543fb52f514b234f | *D_0__Bacteria;D_1__Firmicutes;D_2__Clostridia;D_3__Clostridiales;D_4__Ruminococcaceae;D_5__Ruminococcus 1* | 14 |
| 3219af75ee8056ed06ba81e873b0d0cb | *D_0__Bacteria;D_1__Firmicutes;D_2__Clostridia;D_3__Clostridiales;D_4__Ruminococcaceae;D_5__Ruminococcaceae UCG-002;D_6__uncultured rumen bacterium* | 15 |
| 7427cc5ba277f690df113d56e5c6b538 | *D_0__Bacteria;D_1__Actinobacteria;D_2__Coriobacteriia;D_3__Coriobacteriales;D_4__Coriobacteriaceae;D_5__Olegusella;D_6__Olegusella massiliensis* | 16 |
| 1c719e69d668107c05bdba29829217c5 | *D_0__Bacteria;D_1__Proteobacteria;D_2__Deltaproteobacteria;D_3__Desulfovibrionales;D_4__Desulfovibrionaceae;D_5__Desulfovibrio;D_6__Desulfovibrio longreachensis* | 17 |
| cd604e575361dd5195144bf8e5007cb7 | *D_0__Bacteria;D_1__Firmicutes;D_2__Clostridia;D_3__Clostridiales;D_4__Ruminococcaceae;D_5__Fournierella;D_6__Fournierella massiliensis* | 18 |
| bdb6ddef008a6d31c019642fb28d7c1d | *D_0__Bacteria;D_1__Firmicutes;D_2__Clostridia;D_3__Clostridiales;D_4__Ruminococcaceae;D_5__Ruminiclostridium 9;D_6__Anaerostipes hadrus* | 19 |
| 74d519816d4cfb9cf4ec9605f76301d6 | *D_0__Bacteria;D_1__Firmicutes;D_2__Clostridia;D_3__Clostridiales;D_4__Family XIII;D_5__Family XIII UCG-001;D_6__uncultured bacterium* | 20 |
| a333ee090a52b582e1a0e2dd776eb937 | *D_0__Bacteria;D_1__Firmicutes;D_2__Erysipelotrichia;D_3__Erysipelotrichales;D_4__Erysipelotrichaceae;D_5__Coprobacillus;D_6__Coprobacillus cateniformis JCM 10604* | 21 |
| 9ea77e95fbfd1a99c9c28db3b03a01a8 | *D_0__Bacteria;D_1__Bacteroidetes;D_2__Bacteroidia;D_3__Bacteroidales;D_4__Barnesiellaceae;D_5__Barnesiella;D_6__Barnesiella intestinihominis* | 22 |
| b0e56c25ca193e27096e8fb4eb560eee | *D_0__Bacteria;D_1__Firmicutes;D_2__Clostridia;D_3__Clostridiales;D_4__Lachnospiraceae;D_5__Eisenbergiella;D_6__Eisenbergiella tayi* | 23 |
| 2de0f958e30d26f04cba8d2a920bbe46 | *D_0__Bacteria;D_1__Firmicutes;D_2__Clostridia;D_3__Clostridiales;D_4__Lachnospiraceae;D_5__Blautia;D_6__Blautia faecis* | 24 |
| fec2da4ccc40b9ad85a020553063366b | *D_0__Bacteria;D_1__Proteobacteria;D_2__Gammaproteobacteria;D_3__Betaproteobacteriales;D_4__Burkholderiaceae;D_5__Oxalobacteraceae;D_6__Noviherbaspirillum agri* | 25 |
| 54ec3041ed02d098b3c5e4395466c441 | *D_0__Bacteria;D_1__Firmicutes;D_2__Clostridia;D_3__Clostridiales;D_4__Ruminococcaceae;D_5__Ruminococcaceae UCG-002;D_6__uncultured organism* | 26 |
| ec6732c2e0d4cf64b3d0350e7fe3defb | *D_0__Bacteria;D_1__Firmicutes;D_2__Clostridia;D_3__Clostridiales;D_4__Lachnospiraceae;D_5__Roseburia;D_6__Roseburia inulinivorans DSM 16841* | 27 |
| c6b90711837508687841b1d5cbff2f65 | *D_0__Bacteria;D_1__Firmicutes;D_2__Clostridia;D_3__Clostridiales;D_4__Lachnospiraceae;D_5__uncultured;D_6__uncultured bacterium adhufec382* | 28 |
| 51d4f17b8dda9f7d972bf3dd0ecc4e59 | *D_0__Bacteria;D_1__Bacteroidetes;D_2__Bacteroidia;D_3__Bacteroidales;D_4__Rikenellaceae;D_5__Alistipes;D_6__Alistipes obesi* | 29 |
| eebc27bcf8bcb13a692001530892f5d5 | *D_0__Bacteria;D_1__Firmicutes;D_2__Negativicutes;D_3__Selenomonadales;D_4__Acidaminococcaceae;D_5__Phascolarctobacterium;D_6__uncultured Firmicutes bacterium* | 30 |
| c97ea25ba069877eac769cb05968724c | *D_0__Bacteria;D_1__Firmicutes;D_2__Clostridia;D_3__Clostridiales;D_4__Ruminococcaceae;D_5__Ruminococcaceae UCG-014* | 31 |
| fcffedae608fe4871be42936a1cb523a | *D_0__Bacteria;D_1__Firmicutes;D_2__Clostridia;D_3__Clostridiales;D_4__Lachnospiraceae;D_5__[Ruminococcus] torques group* | 32 |
| 91587a85e342f8dba27f54e15ac0ea77 | *D_0__Bacteria;D_1__Firmicutes;D_2__Clostridia;D_3__Clostridiales;D_4__Lachnospiraceae;D_5__Tyzzerella 3;D_6__unidentified* | 33 |
| aff3962bbd7607a2971161d9840ca409 | *D_0__Bacteria;D_1__Firmicutes;D_2__Clostridia;D_3__Clostridiales;D_4__Ruminococcaceae;D_5__Ruminiclostridium 5* | 34 |
| 44396f15b10f6577d61a11c6047c939f | *D_0__Archaea;D_1__Euryarchaeota;D_2__Methanobacteria;D_3__Methanobacteriales;D_4__Methanobacteriaceae;D_5__Methanobrevibacter;D_6__Methanobrevibacter millerae* | 35 |
| *** The ID of each amplicon sequence variant (ASV)** | | |
| **$ Biomarkers are denoted by ASVs annotated to species and other differential ASVs are denoted by node numbers accordingly** | | |
| **† Taxonomic information of each ASV** | | |

**Supplementary Table 7 | Node numbers of differential ASVs between adenoma and cancer**

| **ASV ID*** | **Taxon^†^** | **node^$^** |
| --- | --- | --- |
| 5149bdbf03de13bb38557d48d5141b87 | *D_0__Bacteria;D_1__Firmicutes;D_2__Clostridia;D_3__Clostridiales;D_4__Lachnospiraceae;D_5__Lachnoclostridium;D_6__[Clostridium] scindens* | *[Clostridium] scindens* |
| 8538d07b8c1e8676f9b9c8c242aab14a | *D_0__Bacteria;D_1__Firmicutes;D_2__Clostridia;D_3__Clostridiales;D_4__Ruminococcaceae;D_5__[Eubacterium] coprostanoligenes group;D_6__unidentified* | *[Eubacterium] coprostanoligenes group sp.* |
| a55a010c9525ce2943a553dca1421b1c | *D_0__Bacteria;D_1__Firmicutes;D_2__Clostridia;D_3__Clostridiales;D_4__Lachnospiraceae;D_5__[Eubacterium] ventriosum group;D_6__uncultured bacterium* | *[Eubacterium] ventriosum group sp.* |
| d1a75593a480570277f05ec729b865de | *D_0__Bacteria;D_1__Firmicutes;D_2__Clostridia;D_3__Clostridiales;D_4__Lachnospiraceae;D_5__[Ruminococcus] gnavus group* | *[Ruminococcus] gnavus group sp.* |
| fd44d4cb468fd7dc9b3227867714ed87 | *D_0__Bacteria;D_1__Bacteroidetes;D_2__Bacteroidia;D_3__Bacteroidales;D_4__Bacteroidaceae;D_5__Bacteroides;D_6__Bacteroides dorei* | *Bacteroides dorei* |
| 7ee3e4343242684ba3ce459672348ff7 | *D_0__Bacteria;D_1__Bacteroidetes;D_2__Bacteroidia;D_3__Bacteroidales;D_4__Bacteroidaceae;D_5__Bacteroides;D_6__Bacteroides nordii* | *Bacteroides nordii* |
| 2de0f958e30d26f04cba8d2a920bbe46 | *D_0__Bacteria;D_1__Firmicutes;D_2__Clostridia;D_3__Clostridiales;D_4__Lachnospiraceae;D_5__Blautia;D_6__Blautia faecis* | *Blautia faecis* |
| d49783e7800974b5f2bcce554e26072e | *D_0__Bacteria;D_1__Firmicutes;D_2__Clostridia;D_3__Clostridiales;D_4__Lachnospiraceae;D_5__Blautia;D_6__Blautia sp.* | *Blautia sp.* |
| e9f900358bb9297e10b9fbb1321b1e6e | *D_0__Bacteria;D_1__Firmicutes;D_2__Erysipelotrichia;D_3__Erysipelotrichales;D_4__Erysipelotrichaceae;D_5__Erysipelatoclostridium;D_6__Erysipelatoclostridium ramosum* | *Erysipelatoclostridium ramosum* |
| f50508546ae13143015f8c4cb976d0e4 | *D_0__Bacteria;D_1__Firmicutes;D_2__Clostridia;D_3__Clostridiales;D_4__Lachnospiraceae;D_5__[Eubacterium] ruminantium group;D_6__Eubacterium ruminantium* | *Eubacterium ruminantium* |
| 8646bffa210e3f061cc57f1b1cf2f751 | *D_0__Bacteria;D_1__Firmicutes;D_2__Clostridia;D_3__Clostridiales;D_4__Lachnospiraceae;D_5__Hungatella;D_6__Hungatella hathewayi WAL-18680* | *Hungatella hathewayi WAL-18680* |
| 6ec1023583647a401269990c4e8ad716 | *D_0__Bacteria;D_1__Firmicutes;D_2__Clostridia;D_3__Clostridiales;D_4__Lachnospiraceae;D_5__Lachnospira;D_6__Lachnospira pectinoschiza* | *Lachnospira pectinoschiza* |
| ab149525cad479f6599c7923acc49691 | *D_0__Bacteria;D_1__Firmicutes;D_2__Clostridia;D_3__Clostridiales;D_4__Lachnospiraceae;D_5__Lachnospiraceae UCG-010;D_6__uncultured organism* | *Lachnospiraceae UCG-010 sp.* |
| ee5b57a629da222ad236b5d4bc329d71 | *D_0__Bacteria;D_1__Firmicutes;D_2__Erysipelotrichia;D_3__Erysipelotrichales;D_4__Erysipelotrichaceae;D_5__Merdibacter;D_6__Merdibacter massiliensis* | *Merdibacter massiliensis* |
| e93b313352019b8113e7fc66bf757917 | *D_0__Bacteria;D_1__Firmicutes;D_2__Clostridia;D_3__Clostridiales;D_4__Family XI;D_5__Parvimonas;;D_6__Parvimonas micra* | *Parvimonas micra* |
| 31f567f59096ed2d57feecc6c3621a58 | *D_0__Bacteria;D_1__Bacteroidetes;D_2__Bacteroidia;D_3__Bacteroidales;D_4__Porphyromonadaceae;D_5__Porphyromonas;D_6__Porphyromonas sp. 2007b* | *Porphyromonas sp. 2007b* |
| 19da890972569b6859c360e9b4c8751e | *D_0__Bacteria;D_1__Bacteroidetes;D_2__Bacteroidia;D_3__Bacteroidales;D_4__Porphyromonadaceae;D_5__Porphyromonas;D_6__Porphyromonas sp. HMSC077F02* | *Porphyromonas sp. HMSC077F02* |
| 4924bfca291797ea1fca3e384f4143f9 | *D_0__Bacteria;D_1__Firmicutes;D_2__Clostridia;D_3__Clostridiales;D_4__Lachnospiraceae;D_6__Roseburia hominis A2-183* | *Roseburia hominis A2-183 sp.* |
| 96040114b5274bdb987f28c63b6b6b87 | *D_0__Bacteria;D_1__Firmicutes;D_2__Clostridia;D_3__Clostridiales;D_4__Lachnospiraceae;D_5_Roseburia;D_6__Roseburia intestinalis* | *Roseburia intestinalis* |
| 16530e5b50b780018210dac05a471715 | *D_0__Bacteria;D_1__Firmicutes;D_2__Clostridia;D_3__Clostridiales;D_4__Ruminococcaceae;D_5__Ruminococcaceae UCG-002* | *Ruminococcaceae UCG-002 sp.* |
| aff3962bbd7607a2971161d9840ca409 | *D_0__Bacteria;D_1__Firmicutes;D_2__Clostridia;D_3__Clostridiales;D_4__Ruminococcaceae;D_5__Ruminiclostridium 5;D_6__Ruminococcus bromii* | *Ruminococcus bromii* |
| 9fec7bdd6bd88e710bd69b15692e54a0 | *D_0__Bacteria;D_1__Firmicutes;D_2__Bacilli;D_3__Lactobacillales;D_4__Streptococcaceae;D_5__Streptococcus;D_6__Streptococcus infantarius* | *Streptococcus infantarius* |
| fd496fd32dc8c08ade2e8b6c9d8ee13d | *D_0__Bacteria;D_1__Firmicutes;D_2__Bacilli;D_3__Lactobacillales;D_4__Streptococcaceae;D_5__Streptococcus;D_6__Streptococcus thermophilus TH1435* | *Streptococcus thermophilus TH1435* |
| 91587a85e342f8dba27f54e15ac0ea77 | *D_0__Bacteria;D_1__Firmicutes;D_2__Clostridia;D_3__Clostridiales;D_4__Lachnospiraceae;D_5__Tyzzerella 3;D_6__unidentified* | *Tyzzerella 3 sp.* |
| 00a96fbd0ac34bfd245f9c24f8737f7d | *D_0__Bacteria;D_1__Firmicutes;D_2__Clostridia;D_3__Clostridiales;D_4__Lachnospiraceae;D_5__Blautia;D_6__Blautia obeum* | 1 |
| a18c0c17fce3d52dc60506e8730bc996 | *D_0__Bacteria;D_1__Firmicutes;D_2__Clostridia;D_3__Clostridiales;D_4__Ruminococcaceae;D_5__Butyricicoccus;D_6__Butyricicoccus faecihominis* | 2 |
| e865f29a716e8d51f83931befa951240 | *D_0__Bacteria;D_1__Firmicutes;D_2__Erysipelotrichia;D_3__Erysipelotrichales;D_4__Erysipelotrichaceae;D_5__Erysipelotrichaceae UCG-003* | 3 |
| 2e4f2b53b856c4def6d021d01f5abb70 | *D_0__Bacteria;D_1__Firmicutes;D_2__Clostridia;D_3__Clostridiales;D_4__Lachnospiraceae;D_5__Dorea;D_6__Dorea longicatena* | 4 |
| 69b90707f6f3da7941026fbe6a1597ce | *D_0__Bacteria;D_1__Firmicutes;D_2__Clostridia;D_3__Clostridiales;D_4__Lachnospiraceae;D_5__Lachnospiraceae FCS020 group;D_6__uncultured organism* | 5 |
| c48070f3061b086b60ff32f77e0002fa | *D_0__Bacteria;D_1__Firmicutes;D_2__Clostridia;D_3__Clostridiales;D_4__Ruminococcaceae;D_5__Subdoligranulum;D_6__Subdoligranulum variabile* | 6 |
| d76d59ec71de0e3b22da0c9cd564d41a | *D_0__Bacteria;D_1__Firmicutes;D_2__Clostridia;D_3__Clostridiales;D_4__Lachnospiraceae;D_5__Fusicatenibacter;D_6__Fusicatenibacter saccharivorans* | 7 |
| 1a9df3fbbc7fc8cecec4239cf801d2fc | *D_0__Bacteria;D_1__Firmicutes;D_2__Clostridia;D_3__Clostridiales;D_4__Ruminococcaceae;D_5__Ruminococcaceae UCG-013* | 8 |
| 9639a3291729a3758207b47715d9205f | *D_0__Bacteria;D_1__Firmicutes;D_2__Clostridia;D_3__Clostridiales;D_4__Ruminococcaceae;D_5__SubdoligranulumD_6__Subdoligranulum variabile* | 9 |
| caea322863a57bb2b14c7317fa120dc4 | *D_0__Bacteria;D_1__Firmicutes;D_2__Clostridia;D_3__Clostridiales;D_4__Lachnospiraceae;D_5__Lachnospiraceae NK4A136 group* | 10 |
| 394eda29c886632f514dd94b58381186 | *D_0__Bacteria;D_1__Proteobacteria;D_2__Gammaproteobacteria;D_3__Pasteurellales;D_4__Pasteurellaceae;D_5__Haemophilus;D_6__Haemophilus parainfluenzae ATCC 33392* | 11 |
| 2c0c62ea09b2efe01bacdcbcf558d2b8 | *D_0__Bacteria;D_1__Firmicutes;D_2__Clostridia;D_3__Clostridiales;D_4__Lachnospiraceae;D_5__Lachnoclostridium;D_6__uncultured Firmicutes bacterium* | 12 |
| 0a035a7ec1f4a194dd7fbda3c5660db0 | *D_0__Bacteria;D_1__Firmicutes;D_2__Clostridia;D_3__Clostridiales;D_4__Lachnospiraceae;D_5__Lachnoclostridium;D_6__human gut metagenome* | 13 |
| 6851a4ee264b56be2fff4686ce269907 | *D_0__Bacteria;D_1__Firmicutes;D_2__Clostridia;D_3__Clostridiales;D_4__Lachnospiraceae;D_5__[Eubacterium] hallii group* | 14 |
| 421cbd1d71d34704c6d53377261e213c | *D_0__Bacteria;D_1__Firmicutes;D_2__Clostridia;D_3__Clostridiales;D_4__Lachnospiraceae;D_5__[Eubacterium] ventriosum group* | 15 |
| 55beed6d1a19ce34788b66488ffe3917 | *D_0__Bacteria;D_1__Firmicutes;D_2__Clostridia;D_3__Clostridiales;D_4__Ruminococcaceae;D_5__Acetanaerobacterium;D_6__Acetanaerobacterium elongatum* | 16 |
| 4e2e6735331ad01a30c94d2f4c285dc2 | *D_0__Bacteria;D_1__Firmicutes;D_2__Clostridia;D_3__Clostridiales;D_4__Lachnospiraceae;D_5__CAG-56;D_6__uncultured bacterium* | 17 |
| a891fb44241e433fa8acf251ef4d328f | *D_0__Bacteria;D_1__Firmicutes;D_2__Clostridia;D_3__Clostridiales;D_4__Lachnospiraceae;D_5__Lacrimispora;D_6__Lacrimispora amygdalina* | 18 |
| 208c0c43c7c5d4a6bb549e0c8365cd21 | *D_0__Bacteria;D_1__Firmicutes;D_2__Clostridia;D_3__Clostridiales;D_4__Lachnospiraceae;D_5__[Eubacterium] eligens group* | 19 |
| e335f74033bc634af43ee6baa84fa247 | *D_0__Bacteria;D_1__Firmicutes;D_2__Clostridia;D_3__Clostridiales;D_4__Peptostreptococcaceae;D_5__Romboutsia;D_6__Romboutsia timonensis* | 20 |
| c97ea25ba069877eac769cb05968724c | *D_0__Bacteria;D_1__Firmicutes;D_2__Clostridia;D_3__Clostridiales;D_4__Ruminococcaceae;D_5__Ruminococcaceae UCG-014* | 21 |
| ee72ee97db064e5d745356f479b9e576 | *D_0__Bacteria;D_1__Firmicutes;D_2__Clostridia;D_3__Clostridiales;D_4__Ruminococcaceae;D_5__Ruminococcus 2* | 22 |
| df16a09f3e448ddd3b1ec54066d3081d | *D_0__Bacteria;D_1__Firmicutes;D_2__Clostridia;D_3__Clostridiales;D_4__Ruminococcaceae;D_5__Butyricicoccus;D_6__Butyricicoccus faecihominis* | 23 |
| f5f5e0da89730462abaf6301a9557193 | *D_0__Bacteria;D_1__Firmicutes;D_2__Clostridia;D_3__Clostridiales;D_4__Ruminococcaceae;D_5__Faecalibacterium;D_6__Faecalibacterium prausnitzii A2-165* | 24 |
| 86ff0efd7fd88b82809500c90c4a7832 | *D_0__Bacteria;D_1__Firmicutes;D_2__Clostridia;D_3__Clostridiales;D_4__Ruminococcaceae;D_5__Faecalibacterium;D_6__Faecalibacterium prausnitzii* | 25 |
| 33518e48b174ac428cb7be1a216c6c6e | *D_0__Bacteria;D_1__Firmicutes;D_2__Clostridia;D_3__Clostridiales;D_4__Lachnospiraceae;D_5__Anaerostipes;D_6__Anaerostipes hadrus* | 26 |
| c66ec67719e54d2cb0afc3154b3c23af | *D_0__Bacteria;D_1__Firmicutes;D_2__Clostridia;D_3__Clostridiales;D_4__Christensenellaceae;D_5__Christensenellaceae R-7 group* | 27 |
| eb51a6482b09f0c5962e362d2d23abb9 | *D_0__Bacteria;D_1__Firmicutes;D_2__Clostridia;D_3__Clostridiales;D_4__Ruminococcaceae;D_5__Ruminococcaceae UCG-014* | 28 |
| aae4c4ed528ae16ec417df816699cfdb | *D_0__Bacteria;D_1__Firmicutes;D_2__Clostridia;D_3__Clostridiales;D_4__Ruminococcaceae;D_5__Ruminococcaceae UCG-005* | 29 |
| 15b5966398a85e5c75501ad243691dfc | *D_0__Bacteria;D_1__Firmicutes;D_2__Clostridia;D_3__Clostridiales;D_4__Lachnospiraceae;D_5__Lachnoclostridium* | 30 |
| 54a2bfb0a96adc52c4ccca239ad01466 | *D_0__Bacteria;D_1__Firmicutes;D_2__Clostridia;D_3__Clostridiales;D_4__Ruminococcaceae;D_5__Ruminiclostridium 6;D_6__Ruminiclostridium hungatei* | 31 |
| 6c324279c1fc2965e594486ac26b0548 | *D_0__Bacteria;D_1__Firmicutes;D_2__Clostridia;D_3__Clostridiales;D_4__Lachnospiraceae;D_5__Coprococcus 2* | 32 |
| 43202fe954709a694e31c3a1be7e889b | *D_0__Bacteria;D_1__Firmicutes;D_2__Clostridia;D_3__Clostridiales;D_4__Ruminococcaceae;D_5__Ruminococcaceae UCG-014* | 33 |
| cd9401a6bce4a63af516d06d2a843f9d | *D_0__Bacteria;D_1__Firmicutes;D_2__Negativicutes;D_3__Selenomonadales;D_4__Veillonellaceae;D_5__Veillonella;D_6__Veillonella parvula DSM 2008* | 34 |
| fb428d07d03e32be0b85eacbed10df0a | *D_0__Bacteria;D_1__Firmicutes;D_2__Clostridia;D_3__Clostridiales;D_4__Lachnospiraceae;D_5__Coprococcus 2;D_6__Coprococcus eutactus* | 35 |
| 369bbf86b13d5924c095aad30bba9af1 | *D_0__Bacteria;D_1__Firmicutes;D_2__Clostridia;D_3__Clostridiales;D_4__Lachnospiraceae;D_5__Blautia;D_6__Blautia schinkii* | 36 |
| c0044f57ec04cd70905d690c5b7bd147 | *D_0__Bacteria;D_1__Firmicutes;D_2__Clostridia;D_3__Clostridiales;D_4__Lachnospiraceae;D_5__Coprococcus 2* | 37 |
| 366bb3ddd63d135bec172051cf2cfe0a | *D_0__Bacteria;D_1__Actinobacteria;D_2__Actinobacteria;D_3__Actinomycetales;D_4__Actinomycetaceae;D_5__Actinomyces;D_6__Actinomyces graevenitzii F0530* | 38 |
| 25bf7b31746631bbb8582ebe892fe837 | *D_0__Bacteria;D_1__Firmicutes;D_2__Clostridia;D_3__Clostridiales;D_4__Ruminococcaceae;D_5__Ruminococcaceae UCG-014* | 39 |
| 0b53415990b1e4435ba981cc617f6be0 | *D_0__Bacteria;D_1__Firmicutes;D_2__Clostridia;D_3__Clostridiales;D_4__Lachnospiraceae;D_5__Lachnospiraceae UCG-003;D_6__uncultured bacterium* | 40 |
| cd5b7f7211526a4109099a6fd4b01aab | *D_0__Bacteria;D_1__Firmicutes;D_2__Clostridia;D_3__Clostridiales;D_4__Defluviitaleaceae;D_5__Defluviitaleaceae UCG-011;D_6__uncultured bacterium* | 41 |
| 54886b8e1a829f2bf0f874ea7a002750 | *D_0__Bacteria;D_1__Firmicutes;D_2__Clostridia;D_3__Clostridiales;D_4__Lachnospiraceae;D_5__Acetitomaculum;D_6__Firmicutes bacterium CAG_194_44_15* | 42 |
| 4e0a8ac6434b87de6ed53d016fd7c87c | *D_0__Bacteria;D_1__Firmicutes;D_2__Clostridia;D_3__Clostridiales;D_4__Family XIII;D_5__Family XIII AD3011 group* | 43 |
| e35e7d4cf6b6c4837a7bf77ca88c05ca | *D_0__Bacteria;D_1__Firmicutes;D_2__Clostridia;D_3__Clostridiales;D_4__Ruminococcaceae;D_5__Ruminococcaceae UCG-010;D_6__uncultured bacterium* | 44 |
| 989f2970e8c86c75bf4b205be62fd345 | *D_0__Bacteria;D_1__Firmicutes;D_2__Clostridia;D_3__Clostridiales;D_4__Ruminococcaceae;D_5__Ruminiclostridium 5;D_6__uncultured bacterium* | 45 |
| 8ee6f70260b29cb40fdff8d4500ae32a | *D_0__Bacteria;D_1__Firmicutes;D_2__Clostridia;D_3__Clostridiales;D_4__Lachnospiraceae;D_5__Roseburia;D_6__Roseburia hominis A2-183* | 46 |
| 8d5153b41e59c5ffb5a1fc080eb784cf | *D_0__Bacteria;D_1__Firmicutes;D_2__Clostridia;D_3__Clostridiales;D_4__Ruminococcaceae;D_5__Ruminococcaceae UCG-004;D_6__uncultured bacterium* | 47 |
| 003e253932cf83d92de80661474011ca | *D_0__Bacteria;D_1__Firmicutes;D_2__Clostridia;D_3__Clostridiales;D_4__Christensenellaceae;D_5__Catabacter;D_6__Christensenella massiliensis* | 48 |
| 1d49df56b83a383d8b998af9d148f52c | *D_0__Bacteria;D_1__Firmicutes;D_2__Clostridia;D_3__Clostridiales;D_4__Clostridiaceae 1;D_5__Clostridium sensu stricto 1* | 49 |
| 0c410683e68edf42337559941e5d3018 | *D_0__Bacteria;D_1__Firmicutes;D_2__Clostridia;D_3__Clostridiales;D_4__Ruminococcaceae;D_5__Ruminococcaceae UCG-013* | 50 |
| b5a69fb7a66fdb2bf437bd286b439706 | *D_0__Bacteria;D_1__Bacteroidetes;D_2__Bacteroidia;D_3__Bacteroidales;D_4__Rikenellaceae;D_5__Alistipes;D_6__Alistipes shahii WAL 8301* | 51 |
| bc7f6c40d5f8e134763db884ca78cb1d | *D_0__Bacteria;D_1__Firmicutes;D_2__Clostridia;D_3__Clostridiales;D_4__Lachnospiraceae;D_5__Blautia;D_6__Blautia producta ATCC 27340* | 52 |
| 9ee9d7f1e7a96963f1ee2668f858fced | *D_0__Bacteria;D_1__Firmicutes;D_2__Clostridia;D_3__Clostridiales;D_4__Family XI;D_5__Anaerococcus;D_6__Anaerococcus vaginalis ATCC 51170* | 53 |
| 868be84b9daccd25a648f872c8d5e9ab | *D_0__Bacteria;D_1__Firmicutes;D_2__Negativicutes;D_3__Selenomonadales;D_4__Veillonellaceae;D_5__uncultured;D_6__uncultured organism* | 54 |
| 12cc4c8a5d46fec0b6738bfe7c6c8a08 | *D_0__Bacteria;D_1__Bacteroidetes;D_2__Bacteroidia;D_3__Bacteroidales;D_4__Bacteroidaceae;D_5__Bacteroides;D_6__Bacteroides intestinalis* | 55 |
| 25394a604bbcd078c80186b300dab8ee | *D_0__Bacteria;D_1__Firmicutes;D_2__Erysipelotrichia;D_3__Erysipelotrichales;D_4__Erysipelotrichaceae;D_5__Holdemanella;D_6__Holdemanella biformis* | 56 |
| 8e2daae0f076acb2d186378cb883210a | *D_0__Bacteria;D_1__Firmicutes;D_2__Clostridia;D_3__Clostridiales;D_4__Ruminococcaceae;D_5__UBA1819* | 57 |
| 5e581b06024ab980e25ee550e0b97247 | *D_0__Bacteria;D_1__Firmicutes;D_2__Clostridia;D_3__Clostridiales;D_4__Christensenellaceae;D_5__Catabacter;D_6__Catabacter hongkongensis* | 58 |
| 6ba98911efd5b1fa1cbba9f869562cce | *D_0__Bacteria;D_1__Synergistetes;D_2__Synergistia;D_3__Synergistales;D_4__Synergistaceae;D_5__Cloacibacillus;D_6__Cloacibacillus evryensis* | 59 |
| 33ba975cdb88b5073a2355ffdf01e9cb | *D_0__Bacteria;D_1__Actinobacteria;D_2__Actinobacteria;D_3__Bifidobacteriales;D_4__Bifidobacteriaceae;D_5__Bifidobacterium;D_6__Bifidobacterium dentium* | 60 |
| 08bd43071a06abf56df235fca6655bea | *D_0__Bacteria;D_1__Firmicutes;D_2__Clostridia;D_3__Clostridiales;D_4__Lachnospiraceae;D_5__[Ruminococcus] torques group* | 61 |
| 24cccadb274fef1f88113e8f019e2a63 | *D_0__Bacteria;D_1__Firmicutes;D_2__Clostridia;D_3__Clostridiales;D_4__Family XIII;D_5__[Eubacterium] nodatum group* | 62 |
| edd418053a744b62aa296a5a85039b3a | *D_0__Bacteria;D_1__Firmicutes;D_2__Clostridia;D_3__Clostridiales;D_4__Ruminococcaceae;D_5__Hydrogenoanaerobacterium;D_6__[Clostridium] methylpentosum* | 63 |
| ef1243f7818d1e35c1e968e8539173cf | *D_0__Bacteria;D_1__Firmicutes;D_2__Clostridia;D_3__Clostridiales;D_4__Ruminococcaceae;D_5__Anaerotruncus;D_6__Anaerotruncus sp. AT3* | 64 |
| 11b7d5a0dd0446562be236a4a9144c75 | *D_0__Bacteria;D_1__Firmicutes;D_2__Clostridia;D_3__Clostridiales;D_4__Ruminococcaceae;D_5__Ruminiclostridium 9;D_6__uncultured Flavonifractor sp.* | 65 |
| c532c2c02cdb1cec71b6500c5d6c2edf | *D_0__Bacteria;D_1__Bacteroidetes;D_2__Bacteroidia;D_3__Bacteroidales;D_4__Porphyromonadaceae;D_5__Porphyromonas;D_6__Porphyromonas asaccharolytica* | 66 |
| d515ae0a0b58aa13869a5966a80c98a9 | *D_0__Bacteria;D_1__Firmicutes;D_2__Clostridia;D_3__Clostridiales;D_4__Lachnospiraceae* | 67 |
| c84d13308b8608a006759b82fcb5e4ac | *D_0__Bacteria;D_1__Firmicutes;D_2__Clostridia;D_3__Clostridiales;D_4__Lachnospiraceae;D_5__Blautia;D_6__Blautia hydrogenotrophica* | 68 |
| 7a6dba47692f821cabeaf266b6b36593 | *D_0__Bacteria;D_1__Firmicutes;D_2__Negativicutes;D_3__Selenomonadales;D_4__Veillonellaceae;D_5__Dialister;D_6__Dialister pneumosintes* | 69 |
| 4bdbaae586dedec92168f93266de0979 | *D_0__Bacteria;D_1__Firmicutes;D_2__Clostridia;D_3__Clostridiales;D_4__Christensenellaceae;D_5__Christensenellaceae R-7 group* | 70 |
| e502ebea2a54a6f839d310d6d57e7d47 | *D_0__Bacteria;D_1__Fusobacteria;D_2__Fusobacteriia;D_3__Fusobacteriales;D_4__Fusobacteriaceae;D_5__Fusobacterium;D_6__Fusobacterium nucleatum subsp.animalis* | 71 |
| 0df6c802966e8670279671824da4f10a | *D_0__Bacteria;D_1__Firmicutes;D_2__Bacilli;D_3__Lactobacillales;D_4__Lactobacillaceae;D_5__Lactobacillus;D_6__Lactobacillus gasseri ATCC 33323* | 72 |
| 6b5c5532185593c41966b2173ea9df1c | *D_0__Bacteria;D_1__Firmicutes;D_2__Clostridia;D_3__Clostridiales;D_4__Lachnospiraceae;D_5__Enterocloster;D_6__Enterocloster lavalensis* | 73 |
| 07699fa14cb8970eee5e8dda19dd5dbe | *D_0__Bacteria;D_1__Firmicutes;D_2__Clostridia;D_3__Clostridiales;D_4__Lachnospiraceae;D_5__Eisenbergiella;D_6__Eisenbergiella massiliensis* | 74 |
| 05dbbf0301a9625e1f1a41470a92f7fe | *D_0__Bacteria;D_1__Firmicutes;D_2__Clostridia;D_3__Clostridiales;D_4__Lachnospiraceae;D_5__Anaerostipes;D_6__Anaerostipes caccae* | 75 |
| f479c23321346918723839e33a3544f4 | *D_0__Bacteria;D_1__Firmicutes;D_2__Clostridia;D_3__Clostridiales;D_4__Ruminococcaceae;D_5__UBA1819* | 76 |
| 4912cd4c1f9a773f88a73c258c25c825 | *D_0__Bacteria;D_1__Firmicutes;D_2__Clostridia;D_3__Clostridiales;D_4__Lachnospiraceae;D_5__[Eubacterium] fissicatena group* | 77 |
| 58bd181f757283f5a4ceebdf90ff6243 | *D_0__Bacteria;D_1__Firmicutes;D_2__Clostridia;D_3__Clostridiales;D_4__Lachnospiraceae;D_5__Sellimonas;D_6__Sellimonas intestinalis* | 78 |
| da8644094f1d480f3b3a6a03a7027ab4 | *D_0__Bacteria;D_1__Firmicutes;D_2__Clostridia;D_3__Clostridiales;D_4__Lachnospiraceae;D_5__Hungatella;D_6__Hungatella effluvii* | 79 |
| a248d70e41d75f4773ef7c94e831a21a | *D_0__Bacteria;D_1__Firmicutes;D_2__Clostridia;D_3__Clostridiales;D_4__Lachnospiraceae;D_5__Faecalimonas;D_6__Faecalimonas umbilicata* | 80 |
| 47f770032218f91b352999ff1f663807 | *D_0__Bacteria;D_1__Firmicutes;D_2__Clostridia;D_3__Clostridiales;D_4__Lachnospiraceae;D_5__Lachnoclostridium;D_6__Lachnoclostridium pacaense* | 81 |
| 52573485a4521e007e4c5217fc5d7c68 | *D_0__Bacteria;D_1__Firmicutes;D_2__Clostridia;D_3__Clostridiales;D_4__Lachnospiraceae;D_5__Anaerotignum;D_6__Anaerotignum aminivorans* | 82 |
| 1f0532e025d966a685fd0d1f3144bc8a | *D_0__Bacteria;D_1__Firmicutes;D_2__Clostridia;D_3__Clostridiales;D_4__Peptostreptococcaceae;D_5__Peptostreptococcus;D_6__Peptostreptococcus stomatis* | 83 |
| 5fbc483f79cd97d3a7253931f37c33e8 | *D_0__Bacteria;D_1__Firmicutes;D_2__Clostridia;D_3__Clostridiales;D_4__Ruminococcaceae;D_5__Flavonifractor;D_6__Flavonifractor plautii* | 84 |
| 7af6fc12809fdafab918881f836460f8 | *D_0__Bacteria;D_1__Firmicutes;D_2__Clostridia;D_3__Clostridiales;D_4__Lachnospiraceae;D_5__Tyzzerella 4* | 85 |
| a663a01aeb4dbb88bbe0a9840e8f64dc | *D_0__Bacteria;D_1__Firmicutes;D_2__Erysipelotrichia;D_3__Erysipelotrichales;D_4__Erysipelotrichaceae;D_5__Erysipelatoclostridium;D_6__Erysipelatoclostridium ramosum* | 86 |
| d46e2205f0c6ecf67b51f83d111c509c | *D_0__Bacteria;D_1__Proteobacteria;D_2__Gammaproteobacteria;D_3__Enterobacteriales;D_4__Enterobacteriaceae;D_5__Escherichia-Shigella* | 87 |
| c24e0e391aa836b5eae25567c7eb89ee | *D_0__Bacteria;D_1__Firmicutes;D_2__Clostridia;D_3__Clostridiales;D_4__Lachnospiraceae;D_5__[Ruminococcus] gnavus group* | 88 |
| 67e8859e108281ee7971084bfa759522 | *D_0__Bacteria;D_1__Firmicutes;D_2__Clostridia;D_3__Clostridiales;D_4__Lachnospiraceae;D_5__Lachnoclostridium* | 89 |
| c67445ed68d61f49e41b7bdd1de019fb | *D_0__Bacteria;D_1__Firmicutes;D_2__Clostridia;D_3__Clostridiales;D_4__Lachnospiraceae;D_5__Lachnoclostridium* | 90 |
| *** The ID of each amplicon sequence variant (ASV)** | | |
| **$ Biomarkers are denoted by ASVs annotated to species and other differential ASVs are denoted by node numbers accordingly** | | |
| **† Taxonomic information of each ASV** | | |

**Supplementary Table 8 | Characteristics of the independent cohorts and non-CRC disease studies**

| **Study** | **Group(N*)** | **Age (average±s.d.)^#^** | **BMI (average±s.d.)** | **Aex F(%)/M(%)^†^** | **country** |
| --- | --- | --- | --- | --- | --- |
| **Independent validation cohorts** | | | | | |
| Validation cohort1 | control(70) | 63.12±8.05 | 27.41±5.52 | 36.27/63.73 | American |
|  | Adenoma(102) | 61.46±9.03 | 26.81±4.41 | 40.00/60.00 |  |
| Validation cohort2 | Adenoma(57) | NA | NA | NA | Chinese |
|  | Cancer(52) |  |  |  |  |
| **Non-CRC disease studies** | | | | | |
| NAFLD^$^ | control(51) | 45.85±19.86 | 26.07±6.83 | 70.59/29.41 | American |
|  | case(18) | 54.00±14.86 | 31.08±6.65 | 66.67/33.33 |  |
| IBS^$^ | control(44) | 39.05±12.92 | 23.82±3.72 | 43.18/56.82 | Chinese |
|  | case(84) | 42.01±11.96 | 23.47±3.52 | 34.52/65.48 |  |
| T2D^$^ | control(214) | 36.34±13.99 | 26.38±5.47 | 78.50/21.5 | American |
|  | case(48) | 51.44±9.26 | 32.47±6.95 | 64.58/35.42 |  |
| IBD^$^ | control(18) | 25±2.74 | NA | 33.33/66.67 | American |
|  | CD(61) | 33.51±19.76 |  | 36.07/63.93 |  |
|  | UC(47) | 41.81±18.41 |  | 48.94/51.06 |  |

*** number of samples**

**# standard deviation**

**† the ratio of percentage of female and male**

**$ The accession numbers of NAFLD, IBS,T2D and IBD studies were PRJEB28350, PRJNA544721, PRJNA541332 and PRJNA82111**

**Supplementary Table 9 | Differentially abundant pathways between control and adenoma (P < 0.05)**

| **Pathway** | **Annotation** | **Surperclass^§^** | **pvalue-FR^‡^** | **pvalue-US2^‡^** | **pvalue-CA^‡^** | **pvalue-meta^#^** | **GFOLD-meta^$^** |
| --- | --- | --- | --- | --- | --- | --- | --- |
| PWY-5677 | succinate fermentation to butanoate | Fermentation | 0.5030 | 0.1823 | 0.7172 | 0.0216 | 0.0355 |
| P461-PWY | hexitol fermentation to lactate, formate, ethanol and acetate | Fermentation | 0.0052 | 0.0676 | 0.3953 | 0.0238 | -0.0801 |
| PWY-5022 | 4-aminobutanoate degradation V | Amine and Polyamine Degradation | 0.0695 | 0.1376 | 0.7958 | 0.0088 | 0.0684 |
| LACTOSECAT-PWY | lactose and galactose degradation I | Carbohydrate Biosynthesis/Degradation | 0.0721 | 0.0737 | 0.7283 | 0.0008 | -0.1184 |
| PWY-7371 | 1,4-dihydroxy-6-naphthoate biosynthesis II | Cofactor, Prosthetic Group, Electron Carrier, and Vitamin Biosynthesis | 0.6103 | 0.2113 | 0.3042 | 0.0491 | -0.0068 |
| HEXITOLDEGSUPER-PWY | superpathway of hexitol degradation (bacteria) | Secondary Metabolite Biosynthesis/Degradation | 0.0048 | 0.1908 | 0.2838 | 0.0186 | -0.0560 |
| PWY0-1241 | ADP-L-glycero-beta-D-manno-heptose biosynthesis | Carbohydrate Biosynthesis/Degradation | 0.1393 | 0.1757 | 0.8073 | 0.0030 | 0.0802 |
| SO4ASSIM-PWY | assimilatory sulfate | Inorganic Nutrient Metabolism | 0.0064 | 0.8765 | 0.9705 | 0.0482 | 0.0436 |
| PWY-6263 | superpathway of menaquinol-8 biosynthesis II | Cofactor, Prosthetic Group, Electron Carrier, and Vitamin Biosynthesis | 0.5811 | 0.2082 | 0.3478 | 0.0483 | -0.0041 |
| PWY-6478 | GDP-D-glycero-alpha-D-manno-heptose biosynthesis | Carbohydrate Biosynthesis/Degradation | 0.0482 | 0.8094 | 0.8650 | 0.0017 | 0.0420 |
| PWY-7220 | adenosine deoxyribonucleotides de novo biosynthesis II | Nucleoside and Nucleotide Biosynthesis/Degradation | 0.0762 | 0.5558 | 0.4825 | 0.0080 | 0.0054 |
| PWY-6609 | adenine and adenosine salvage III | Nucleoside and Nucleotide Biosynthesis/Degradation | 0.2140 | 0.1405 | 0.3711 | 0.0208 | -0.0098 |
| ASPASN-PWY | superpathway of L-aspartate and L-asparagine biosynthesis | Amino Acid Biosynthesis/Degradation | 0.3390 | 0.1407 | 0.0024 | 0.0235 | -0.0135 |
| GLYCOLYSIS-E-D | superpathway of glycolysis and the Entner-Doudoroff pathway | Others | 0.0116 | 0.0772 | 0.5997 | 0.0187 | 0.0305 |
| PRPP-PWY | superpathway of histidine, purine, and pyrimidine biosynthesis | Others | 0.0248 | 0.1218 | 0.6627 | 0.0265 | 0.0308 |
| PWY-6126 | superpathway of adenosine nucleotides de novo biosynthesis II | Nucleoside and Nucleotide Biosynthesis/Degradation | 0.1989 | 0.2244 | 0.5298 | 0.0186 | 0.0030 |
| PWY-4984 | urea cycle | Inorganic Nutrient Metabolism | 0.3476 | 0.4055 | 0.9705 | 0.0127 | 0.0442 |
| P162-PWY | L-glutamate degradation V (via hydroxyglutarate) | Amino Acid Biosynthesis/Degradation | 0.0895 | 0.4944 | 0.6520 | 0.0031 | 0.0420 |
| PWY-5345 | superpathway of L-methionine biosynthesis (by sulfhydrylation) | Amino Acid Biosynthesis/Degradation | 0.0085 | 0.2110 | 0.2643 | 0.0075 | 0.0761 |
| PWY0-1261 | anhydromuropeptides recycling I | Secondary Metabolite Biosynthesis/Degradation | 0.0098 | 0.5217 | 0.1188 | 0.0018 | -0.0131 |
| PWY-7392 | taxadiene biosynthesis (engineered) | Secondary Metabolite Biosynthesis/Degradation | 0.4211 | 0.0201 | 0.6520 | 0.0317 | 0.0407 |
| PWY-7222 | guanosine deoxyribonucleotides de novo biosynthesis II | Nucleoside and Nucleotide Biosynthesis/Degradation | 0.0762 | 0.5558 | 0.4825 | 0.0080 | 0.0054 |
| PWY-3781 | aerobic respiration I (cytochrome c) | Electron Transfer | 0.0644 | 0.9351 | 0.9234 | 0.0071 | 0.0417 |
| PWY-7374 | 1,4-dihydroxy-6-naphthoate biosynthesis I | Cofactor, Prosthetic Group, Electron Carrier, and Vitamin Biosynthesis | 0.4067 | 0.2103 | NA | 0.0002 | 0.0632 |
| PWY-5507 | adenosylcobalamin biosynthesis I (anaerobic) | Cofactor, Prosthetic Group, Electron Carrier, and Vitamin Biosynthesis | 0.3696 | 0.9024 | 0.4917 | 0.0457 | 0.0385 |
| PWY-5182 | toluene degradation II (aerobic) (via 4-methylcatechol) | Aromatic Compound Degradation | 0.3222 | 0.9607 | 0.9587 | 0.0012 | -0.0324 |
| PWY-5180 | toluene degradation I (aerobic) (via o-cresol) | Aromatic Compound Degradation | 0.3222 | 0.9607 | 0.9587 | 0.0012 | -0.0324 |

**§ surperclass of pathway**

**‡ Single-study P-value calculated by a two-sided Wilcoxon test**

**# Meta-analysis P-value calculated by two-sided a blocked Wilcoxon test (n=559 independent observations)**

**$ Mean of generalized fold changes across studies, GFOLD-meta >0: pathways enriched in adenoma compared with control; <0: pathways enriched in control compared with adenoma**

**Supplementary Table 10 | Differentially abundant pathways between adenoma and cancer (P < 0.05)**

| **Pathway** | **Annotation** | **Surperclass^§^** | **pvalue-FR^‡^** | **pvalue-US1^‡^** | **pvalue-US2^‡^** | **pvalue-CA^‡^** | **pvalue-meta^#^** | **GFOLD-meta^$^** |
| --- | --- | --- | --- | --- | --- | --- | --- | --- |
| P164-PWY | purine nucleobases degradation I (anaerobic) | Nucleoside and Nucleotide Biosynthesis/Degradation | 0.0869 | 0.1168 | 0.0169 | 0.3255 | 0.0078 | 0.0305 |
| PWY-7013 | (S)-propane-1,2-diol degradation | Fermentation | 0.0020 | 0.0341 | 0.0002 | 0.0537 | 0.0000 | 0.2501 |
| PWY-7377 | cob(II)yrinate a,c-diamide biosynthesis I (early cobalt insertion) | Cofactor, Prosthetic Group, Electron Carrier, and Vitamin Biosynthesis | 0.1451 | 0.7678 | 0.0702 | 0.6952 | 0.0220 | 0.1194 |
| TEICHOICACID-PWY | poly(glycerol phosphate) wall teichoic acid biosynthesis | Cell Structure Biosynthesis | 0.5593 | 0.2007 | 0.3693 | 0.0406 | 0.0410 | -0.0647 |
| PWY-6588 | pyruvate fermentation to acetone | Fermentation | 0.0362 | 0.1199 | 0.0614 | 0.5997 | 0.0460 | 0.0233 |
| PWY-5677 | succinate fermentation to butanoate | Fermentation | 0.0320 | 0.7280 | 0.0276 | 0.1907 | 0.0012 | 0.0926 |
| PWY-5845 | superpathway of menaquinol-9 biosynthesis | Cofactor, Prosthetic Group, Electron Carrier, and Vitamin Biosynthesis | 0.1271 | 0.0051 | 0.0549 | 0.9117 | 0.0025 | 0.1585 |
| PWY-5705 | allantoin degradation to glyoxylate III | Amine and Polyamine Degradation | 0.0469 | 0.0798 | 0.0689 | 0.9352 | 0.0040 | 0.0883 |
| CENTFERM-PWY | pyruvate fermentation to butanoate | Fermentation | 0.1562 | 0.0095 | 0.6187 | 0.4376 | 0.0357 | 0.1279 |
| PWY-5097 | L-lysine biosynthesis VI | Amino Acid Biosynthesis/Degradation | 0.0290 | 0.0513 | 0.9804 | 0.0657 | 0.0307 | -0.0143 |
| GLUCARDEG-PWY | D-glucarate degradation I | Carboxylate Degradation | 0.0136 | 0.3617 | 0.1071 | 0.2905 | 0.0063 | 0.1363 |
| ALL-CHORISMATE-PWY | superpathway of chorismate metabolism | Others | 0.1271 | 0.0029 | 0.3345 | 0.7845 | 0.0159 | 0.2003 |
| PWY-5913 | partial TCA cycle (obligate autotrophs) | TCA cycle | 0.0551 | 0.0868 | 0.3418 | 0.9470 | 0.0357 | 0.0470 |
| PWY-5022 | 4-aminobutanoate degradation V | Amine and Polyamine Degradation | 0.3193 | 0.0363 | 0.0105 | 0.2973 | 0.0004 | 0.1111 |
| PWY490-3 | nitrate reduction VI (assimilatory) | Inorganic Nutrient Metabolism | 0.0276 | 0.0067 | 0.2045 | 0.6735 | 0.0031 | -0.0864 |
| PWY-5918 | superpathway of heme b biosynthesis from glutamate | Cofactor, Prosthetic Group, Electron Carrier, and Vitamin Biosynthesis | 0.7129 | 0.1364 | 0.0014 | 0.0905 | 0.0000 | 0.1504 |
| PWY-6895 | superpathway of thiamine diphosphate biosynthesis II | Cofactor, Prosthetic Group, Electron Carrier, and Vitamin Biosynthesis | 0.0063 | 0.1961 | 0.3405 | 0.4553 | 0.0055 | 0.0567 |
| PWY-6478 | GDP-D-glycero-alpha-D-manno-heptose biosynthesis | Carbohydrate Biosynthesis/Degradation | 0.7873 | 0.0026 | 0.0886 | 0.7394 | 0.0175 | -0.0521 |
| HISDEG-PWY | L-histidine degradation I | Amino Acid Biosynthesis/Degradation | 0.1176 | 0.1829 | 0.9754 | 0.0033 | 0.0365 | 0.1319 |
| PWY-5850 | superpathway of menaquinol-6 biosynthesis I | Cofactor, Prosthetic Group, Electron Carrier, and Vitamin Biosynthesis | 0.1271 | 0.0055 | 0.0524 | 0.9117 | 0.0025 | 0.1580 |
| PWY-6471 | peptidoglycan biosynthesis IV (Enterococcus faecium) | Cell Structure Biosynthesis | 0.0030 | 0.1703 | 0.8775 | 0.1260 | 0.0228 | -0.0555 |
| HEMESYN2-PWY | heme b biosynthesis II (oxygen-independent) | Cofactor, Prosthetic Group, Electron Carrier, and Vitamin Biosynthesis | 0.4588 | 0.1399 | 0.0176 | 0.2905 | 0.0012 | 0.1126 |
| REDCITCYC | TCA cycle VI (Helicobacter) | TCA cycle | 0.8253 | 0.0088 | 0.6528 | 0.2116 | 0.0491 | 0.1069 |
| PWY-5667 | CDP-diacylglycerol biosynthesis I | Fatty Acid and Lipid Biosynthesis/Degradation | 0.0148 | 0.0652 | 0.7478 | 0.0351 | 0.0492 | -0.0164 |
| PWY-7003 | glycerol degradation to butanol | Fermentation | 0.0887 | 0.1961 | 0.1446 | 0.9823 | 0.0293 | 0.1064 |
| PWY-5896 | superpathway of menaquinol-10 biosynthesis | Cofactor, Prosthetic Group, Electron Carrier, and Vitamin Biosynthesis | 0.1271 | 0.0055 | 0.0524 | 0.9117 | 0.0025 | 0.1580 |
| HEME-BIOSYNTHESIS-II | heme b biosynthesis I (aerobic) | Cofactor, Prosthetic Group, Electron Carrier, and Vitamin Biosynthesis | 0.6065 | 0.1231 | 0.0014 | 0.0824 | 0.0000 | 0.1606 |
| PWY-6891 | thiazole biosynthesis II (aerobic bacteria) | Cofactor, Prosthetic Group, Electron Carrier, and Vitamin Biosynthesis | 0.0048 | 0.3414 | 0.0144 | 0.5592 | 0.0006 | 0.0846 |
| PWY0-1415 | superpathway of heme b biosynthesis from uroporphyrinogen-III | Cofactor, Prosthetic Group, Electron Carrier, and Vitamin Biosynthesis | 0.3589 | 0.1961 | 0.1825 | 0.1494 | 0.0103 | 0.1331 |
| PWY-6749 | CMP-legionaminate biosynthesis I | Carbohydrate Biosynthesis/Degradation | 0.0313 | 0.0513 | 0.1896 | 0.9117 | 0.0116 | -0.0805 |
| PWY-5971 | palmitate biosynthesis II (bacteria and plant cytoplasm) | Fatty Acid and Lipid Biosynthesis/Degradation | 0.2294 | 0.9634 | 0.0144 | 0.3790 | 0.0244 | -0.0239 |
| FAO-PWY | fatty acid beta-oxidation I (generic) | Fatty Acid and Lipid Biosynthesis/Degradation | 0.3745 | 0.0060 | 0.4698 | 0.5493 | 0.0176 | 0.1527 |
| P163-PWY | L-lysine fermentation to acetate and butanoate | Amino Acid Biosynthesis/Degradation | 0.0000 | 0.9216 | 0.0042 | 0.0261 | 0.0000 | 0.1882 |
| PWY-6590 | superpathway of Clostridium acetobutylicum acidogenic fermentation | Fermentation | 0.1680 | 0.0095 | 0.6143 | 0.4290 | 0.0363 | 0.1259 |
| PWY0-1319 | CDP-diacylglycerol biosynthesis II | Fatty Acid and Lipid Biosynthesis/Degradation | 0.0148 | 0.0652 | 0.7478 | 0.0351 | 0.0492 | -0.0164 |
| P162-PWY | L-glutamate degradation V (via hydroxyglutarate) | Amino Acid Biosynthesis/Degradation | 0.0017 | 0.4427 | 0.0724 | 0.0067 | 0.0000 | 0.1824 |
| PWY-6545 | pyrimidine deoxyribonucleotides de novo biosynthesis III | Nucleoside and Nucleotide Biosynthesis/Degradation | 0.6839 | 0.8184 | 0.0108 | 0.9470 | 0.0119 | -0.0063 |
| PWY-6151 | S-adenosyl-L-methionine cycle I | Amino Acid Biosynthesis/Degradation | 0.0103 | 0.2968 | 0.7072 | 0.1373 | 0.0222 | -0.0293 |
| PWY-5862 | superpathway of demethylmenaquinol-9 biosynthesis | Cofactor, Prosthetic Group, Electron Carrier, and Vitamin Biosynthesis | NA | 0.0051 | 0.0486 | 0.9352 | 0.0061 | 0.1106 |
| PWY0-1533 | methylphosphonate degradation I | Inorganic Nutrient Metabolism | 0.2960 | 0.0483 | NA | 0.1958 | 0.0087 | 0.1217 |
| PWY-5860 | superpathway of demethylmenaquinol-6 biosynthesis I | Cofactor, Prosthetic Group, Electron Carrier, and Vitamin Biosynthesis | NA | 0.0057 | 0.0460 | 0.9352 | 0.0060 | 0.1103 |

**§ surperclass of pathway**

**‡ Single-study P-value calculated by a two-sided Wilcoxon test**

**# Meta-analysis P-value calculated by a two-sided blocked Wilcoxon test (n=559 independent observations)**

**$ Mean of generalized fold changes across studies, GFOLD-meta >0: pathways enriched in cancer compared with adenoma; <0: pathways enriched in adenoma compared with cancer**

**Supplementary Table 11 | Average distribution of each ASV of healthy subjects and adenoma patients.**

| **ASV ID** | |  |  | **2740cf2417c92847cc298cbd71dd1fcd^‡^** | | **410e1eaa1468a3d40595898f671ba0b3** | | **94928715987dcb9638dd5c0e8f9b20f4** | | **aae4c4ed528ae16ec417df816699cfdb** | | **f50508546ae13143015f8c4cb976d0e4** | |
| --- | --- | --- | --- | --- | --- | --- | --- | --- | --- | --- | --- | --- | --- |
|  | **Function** | **Annotation** |  | **Control** | **Adenoma** | **Control** | **Adenoma** | **Control** | **Adenoma** | **Control** | **Adenoma** | **Control** | **Adenoma** |
|  | ASPASN-PWY | superpathway of L-aspartate and L-asparagine biosynthesis | contribution^†^ | 0.000593277 | 0.001045595 | 0.011903603 | 0.023271737 | 0.000203276 | 0.000240273 | 0.001952301 | 0.001687822 | 0.00345106 | 0.004987388 |
|  |  |  | rank^#^ | 174 | 131 | 24 | 7 | 220 | 219 | 97 | 103 | 65 | 51 |
|  | GLYCOLYSIS-E-D | superpathway of glycolysis and the Entner-Doudoroff pathway | contribution | 0.000403193 | 0.001147156 | 0.008631485 | 0.020062148 | 0.000596136 | 0.000493945 | 0.001387401 | 0.001289289 | 0.004273701 | 0.006154488 |
|  |  |  | rank | 200 | 133 | 32 | 11 | 182 | 184 | 126 | 121 | 58 | 43 |
|  | HEXITOLDEGSUPER-PWY | superpathway of hexitol degradation (bacteria) | contribution | 0.000388314 | 0.001251032 | 0.008232877 | 0.010564415 | 0.000523762 | 0.000456946 | 0.001045323 | 0.000743858 | 0.003304705 | 0.003134257 |
|  |  |  | rank | 196 | 117 | 29 | 26 | 183 | 185 | 134 | 156 | 65 | 65 |
|  | LACTOSECAT-PWY | lactose and galactose degradation I | contribution | 0 | 0 | 0 | 0 | 0 | 0 | 0 | 0 | 0 | 0 |
|  |  |  | rank | 38 | 38 | 38 | 38 | 38 | 38 | 38 | 38 | 38 | 38 |
|  | P162-PWY | L-glutamate degradation V (via hydroxyglutarate) | contribution | 0.001985712 | 0.010342365 | 0 | 0 | 0.000637255 | 0.000659999 | 0 | 0 | 0.003663262 | 0.004398685 |
|  |  |  | rank | 69 | 22 | 134 | 134 | 105 | 113 | 134 | 134 | 49 | 48 |
|  | P461-PWY | hexitol fermentation to lactate, formate, ethanol and acetate | contribution | 0.000312109 | 0.000910459 | 0.008811259 | 0.011331728 | 0.000446214 | 0.000417183 | 0.000971773 | 0.000742398 | 0.002469307 | 0.002580792 |
|  |  |  | rank | 203 | 139 | 28 | 23 | 189 | 185 | 141 | 156 | 77 | 71 |
|  | PRPP-PWY | superpathway of histidine, purine, and pyrimidine biosynthesis | contribution | 0.000658088 | 0.001293567 | 0.010997153 | 0.020856466 | 0.000702521 | 0.000578328 | 0.00143049 | 0.001302287 | 0.003010133 | 0.005278749 |
|  |  |  | rank | 175 | 127 | 26 | 10 | 169 | 178 | 124 | 126 | 74 | 52 |
|  | **PWY0-1241** | **ADP-L-glycero-beta-D-manno-heptose biosynthesis** | contribution | **0.011182423** | **0.022622036** | 0 | 0 | 0.005228543 | 0.003221304 | 0 | 0 | 0 | 0 |
|  |  |  | rank | **16** | **9** | 59 | 59 | 28 | 32 | 59 | 59 | 59 | 59 |
|  | PWY0-1261 | anhydromuropeptides recycling I | contribution | 0.000602215 | 0.001273287 | 0.01615139 | 0.035079922 | 0.000335688 | 0.000423438 | 0.002046811 | 0.002130957 | 0.002552123 | 0.006461994 |
|  |  |  | rank | 173 | 126 | 13 | 4 | 201 | 191 | 100 | 93 | 93 | 45 |
|  | PWY-3781 | aerobic respiration I (cytochrome c) | contribution | 0.000511039 | 0.000247148 | 0.013622996 | 0.028707053 | 0.002617833 | 0.001208936 | 0.001620326 | 0.001715191 | 0.001688841 | 0.00431815 |
|  |  |  | rank | 139 | 173 | 15 | 6 | 65 | 98 | 89 | 77 | 85 | 45 |
|  | PWY-4984 | urea cycle | contribution | 0.000770918 | 0.002125625 | 0.013618288 | 0.019351871 | 0.001265471 | 0.000630042 | 0.001604179 | 0.001497826 | 0.004289992 | 0.005158358 |
|  |  |  | rank | 165 | 98 | 18 | 12 | 131 | 170 | 117 | 121 | 62 | 53 |
|  | PWY-5022 | 4-aminobutanoate degradation V | contribution | 0.001427503 | 0.002976295 | 0 | 0 | 0 | 0 | 0 | 0 | 0 | 0 |
|  |  |  | rank | 61 | 50 | 89 | 89 | 89 | 89 | 89 | 89 | 89 | 89 |
|  | PWY-5180 | toluene degradation I (aerobic) (via o-cresol) | contribution | 0 | 0 | 0.013783622 | 0.027360165 | 0 | 0 | 0.001584038 | 0.001509357 | 0.005815304 | 0.00608798 |
|  |  |  | rank | 147 | 147 | 17 | 10 | 147 | 147 | 81 | 74 | 36 | 34 |
|  | PWY-5182 | toluene degradation II (aerobic) (via 4-methylcatechol) | contribution | 0 | 0 | 0.013783622 | 0.027360165 | 0 | 0 | 0.001584038 | 0.001509357 | 0.005815304 | 0.00608798 |
|  |  |  | rank | 147 | 147 | 17 | 10 | 147 | 147 | 81 | 74 | 36 | 34 |
|  | PWY-5345 | superpathway of L-methionine biosynthesis (by sulfhydrylation) | contribution | 0.000356504 | 0.000814403 | 0.011235588 | 0.020933173 | 0.000459579 | 0.000450592 | 0.001732152 | 0.001455467 | 0.002734861 | 0.00518269 |
|  |  |  | rank | 195 | 147 | 25 | 11 | 179 | 182 | 105 | 113 | 79 | 48 |
|  | PWY-5507 | adenosylcobalamin biosynthesis I (anaerobic) | contribution | 0.000956215 | 0.001505569 | 0.01330191 | 0.045923223 | 0.000327901 | 0.000244687 | 0.002082131 | 0.002483651 | 0.004936982 | 0.006724306 |
|  |  |  | rank | 133 | 108 | 20 | 3 | 181 | 191 | 96 | 82 | 47 | 36 |
|  | PWY-5677 | succinate fermentation to butanoate | contribution | 0.001242452 | 0.002847445 | 0 | 0 | 0 | 0 | 0 | 0 | 0 | 0 |
|  |  |  | rank | 67 | 48 | 88 | 88 | 88 | 88 | 88 | 88 | 88 | 88 |
|  | PWY-6126 | superpathway of adenosine nucleotides de novo biosynthesis II | contribution | 0.000726472 | 0.001229869 | 0.009540959 | 0.017868807 | 0.00064867 | 0.000626097 | 0.001494552 | 0.001268014 | 0.003049205 | 0.004370994 |
|  |  |  | rank | 170 | 131 | 28 | 13 | 179 | 170 | 122 | 126 | 75 | 57 |
|  | PWY-6263 | superpathway of menaquinol-8 biosynthesis II | contribution | 0.000716662 | 0.002089544 | 0.012315818 | 0.025144994 | 0.000728901 | 0.00122203 | 0.001190845 | 0.001336784 | 0.001979978 | 0.003666437 |
|  |  |  | rank | 153 | 91 | 21 | 6 | 150 | 128 | 124 | 120 | 95 | 62 |
|  | PWY-6478 | GDP-D-glycero-alpha-D-manno-heptose biosynthesis | contribution | 0.005441458 | 0.007098287 | 0 | 0 | 0.005362753 | 0.004217603 | 0 | 0 | 0 | 0 |
|  |  |  | rank | 38 | 32 | 58 | 58 | 39 | 43 | 58 | 58 | 58 | 58 |
|  | PWY-6609 | adenine and adenosine salvage III | contribution | 0.000188137 | 0.000340846 | 0.009170758 | 0.017081053 | 0.000630593 | 0.000592024 | 0.001411472 | 0.001203893 | 0.003868985 | 0.006162149 |
|  |  |  | rank | 228 | 203 | 27 | 16 | 172 | 171 | 125 | 122 | 58 | 38 |
|  | PWY-7220 | adenosine deoxyribonucleotides de novo biosynthesis II | contribution | 0.001165782 | 0.001863479 | 0.009344541 | 0.017025683 | 0.000624165 | 0.000614731 | 0.001409972 | 0.001209334 | 0.001917465 | 0.003166209 |
|  |  |  | rank | 132 | 100 | 30 | 12 | 172 | 165 | 124 | 122 | 98 | 68 |
|  | PWY-7222 | guanosine deoxyribonucleotides de novo biosynthesis II | contribution | 0.001165782 | 0.001863479 | 0.009344541 | 0.017025683 | 0.000624165 | 0.000614731 | 0.001409972 | 0.001209334 | 0.001917465 | 0.003166209 |
|  |  |  | rank | 132 | 100 | 30 | 12 | 172 | 165 | 124 | 122 | 98 | 68 |
|  | PWY-7371 | 1,4-dihydroxy-6-naphthoate biosynthesis II | contribution | 0 | 0 | 0 | 0 | 0 | 0 | 0 | 0 | 0 | 0 |
|  |  |  | rank | 12 | 12 | 12 | 12 | 12 | 12 | 12 | 12 | 12 | 12 |
|  | PWY-7374 | 1,4-dihydroxy-6-naphthoate biosynthesis I | contribution | 0 | 0 | 0 | 0 | 0 | 0 | 0 | 0 | 0 | 0 |
|  |  |  | rank | 12 | 12 | 12 | 12 | 12 | 12 | 12 | 12 | 12 | 12 |
|  | PWY-7392 | taxadiene biosynthesis (engineered) | contribution | 0.000533428 | 0.001389394 | 0.009476037 | 0.023645896 | 0.000916772 | 0.00073712 | 0.001785576 | 0.001482304 | 0.003420451 | 0.004350249 |
|  |  |  | rank | 189 | 120 | 26 | 8 | 148 | 161 | 112 | 116 | 74 | 58 |
|  | SO4ASSIM-PWY | assimilatory sulfate | contribution | 0 | 0 | 0 | 0 | 0 | 0 | 0 | 0 | 0 | 0 |
|  |  |  | rank | 72 | 72 | 72 | 72 | 72 | 72 | 72 | 72 | 72 | 72 |

**‡ The ASV ID asigned as Veillonella.**

**† Defined as the proportion of a per ASV function abundance compared to the total function abundance of all ASVs corresponding to a given pathway.**

**# The rank of the contribution in control and adenoma.**

**Supplementary Table 12 | Candidated genes of ADP-heptose and MK-10 biosynthesis**

| **Control versus Adenoma** | | | |
| --- | --- | --- | --- |
| **ADP-L-glycero-beta-D-manno-heptose biosynthesis (ADP-heptose)** | | | |
| **Enzyme** | **GFOLD-meta$** | **pvalue-meta#** | **Gene name** |
| EC: 5.3.1.28 | 0.0712 | 0.0046 | *gmhA* |
| EC: 2.7.1.167 | 0.0838 | 0.0042 | *hldE* |
| EC: 3.1.3.82 | 0.0706 | 0.0014 | *gmhB* |
| EC: 2.7.7.70 | 0.0838 | 0.0042 | *hldE* |
| EC: 5.1.3.20 | 0.0675 | 0.0185 | *rfaD* |
| **Adenoma versus Cancer** | | |  |
| **menaquinol-10 biosynthesis (MK-10)** | | |  |
| **Enzyme** | **GFOLD-meta†** | **pvalue-meta#** | **Gene name** |
| EC: 4.2.1.113 | 0.0798 | 0.0259 | *menC* |
| EC: 4.2.99.20 | 0.0870 | 0.0459 | *menH* |
| EC: 5.4.4.2 | 0.0967 | 0.0491 | *menF* |

**# Meta-analysis P-value calculated by two-sided a blocked Wilcoxon test**

**$ Mean of generalized fold changes across studies, GFOLD-meta >0: pathways enriched in adenoma compared with control; <0: pathways enriched in control compared with adenoma**

**† Mean of generalized fold changes across studies, GFOLD-meta >0: pathways enriched in cancer compared with adenoma; <0: pathways enriched in adenoma compared with cancer**

**Supplementary Table 13 | Average distribution of each ASV of adenoma patients and CRC.**

| **ASV ID** |  |  | **fd44d4cb468fd7dc9b3227867714ed87^§^** | | **5149bdbf03de13bb38557d48d5141b87** | | **d49783e7800974b5f2bcce554e26072e** | | **8538d07b8c1e8676f9b9c8c242aab14a** | | **7ee3e4343242684ba3ce459672348ff7** | |
| --- | --- | --- | --- | --- | --- | --- | --- | --- | --- | --- | --- | --- |
| **Function** | **Annotation** |  | **Adenoma** | **Cancer** | **Adenoma** | **Cancer** | **Adenoma** | **Cancer** | **Adenoma** | **Cancer** | **Adenoma** | **Cancer** |
| ALL-CHORISMATE-PWY | superpathway of chorismate metabolism | contribution^†^ | 0.023689928 | 0.034035749 | 0.000398435 | 0.000693012 | 0.002378832 | 0.003983944 | 0.001242809 | 0.0017666 | 0 | 0 |
|  |  | rank^#^ | 8 | 3 | 182 | 168 | 80 | 56 | 121 | 101 | 246 | 246 |
| CENTFERM-PWY | pyruvate fermentation to butanoate | contribution | 0.031513165 | 0.031019502 | 0.00010402 | 0.000231819 | 0.004295877 | 0.004227672 | 0.000328045 | 0.000481385 | 0.000420556 | 0.001138968 |
|  |  | rank | 7 | 6 | 234 | 203 | 57 | 55 | 195 | 177 | 185 | 120 |
| FAO-PWY | fatty acid beta-oxidation I (generic) | contribution | 0.06781719 | 0.049320517 | 0.00054727 | 0.000783752 | 0.00177546 | 0.001651262 | 0 | 0 | 0.000740283 | 0.002236989 |
|  |  | rank | 3 | 3 | 128 | 110 | 81 | 77 | 165 | 165 | 115 | 64 |
| GLUCARDEG-PWY | D-glucarate degradation I | contribution | 0 | 0 | 0 | 0 | 0.011995189 | 0.006777061 | 0 | 0 | 0 | 0 |
|  |  | rank | 46 | 46 | 46 | 46 | 17 | 18 | 46 | 46 | 46 | 46 |
| HEME-BIOSYNTHESIS-II | heme b biosynthesis I (aerobic) | contribution | 0.042894099 | 0.024703493 | 0.001166784 | 0.002957835 | 0.005011703 | 0.006775788 | 0 | 0 | 0 | 0 |
|  |  | rank | 7 | 9 | 70 | 49 | 42 | 34 | 99 | 99 | 99 | 99 |
| HEMESYN2-PWY | heme b biosynthesis II (oxygen-independent) | contribution | 0.017220372 | 0.017019092 | 0.000513216 | 0.001392312 | 0.004881156 | 0.00514924 | 0.00070733 | 0.001305806 | 0.000131906 | 0.000422847 |
|  |  | rank | 15 | 11 | 174 | 124 | 53 | 52 | 156 | 131 | 237 | 198 |
| HISDEG-PWY | L-histidine degradation I | contribution | 0.118686936 | 0.088265721 | 0 | 0 | 0.027861103 | 0.024971203 | 0 | 0 | 0.002246194 | 0.004330958 |
|  |  | rank | 2 | 2 | 38 | 38 | 11 | 13 | 38 | 38 | 31 | 26 |
| P162-PWY | L-glutamate degradation V (via hydroxyglutarate) | contribution | 0 | 0 | 0 | 0 | 0 | 0 | 0 | 0 | 0 | 0 |
|  |  | rank | 134 | 134 | 134 | 134 | 134 | 134 | 134 | 134 | 134 | 134 |
| P163-PWY | L-lysine fermentation to acetate and butanoate | contribution | 0.018129344 | 0.014579515 | 0.000272532 | 0.001119317 | 0.000493563 | 0.000476418 | 0.001379973 | 0.001819781 | 0.000760502 | 0.001773039 |
|  |  | rank | 14 | 15 | 216 | 140 | 193 | 195 | 118 | 107 | 164 | 109 |
| P164-PWY | purine nucleobases degradation I (anaerobic) | contribution | 0.015107292 | 0.017550204 | 0.001272697 | 0.002808837 | 0.014224494 | 0.013107167 | 0.000795403 | 0.001410989 | 0.00036987 | 0.00103727 |
|  |  | rank | 11 | 8 | 112 | 72 | 14 | 16 | 144 | 110 | 187 | 133 |
| PWY0-1319 | CDP-diacylglycerol biosynthesis II | contribution | 0.011458406 | 0.010957328 | 0.000440629 | 0.001083098 | 0.002829319 | 0.002774998 | 0.001305202 | 0.002678607 | 0.000279758 | 0.000692743 |
|  |  | rank | 21 | 23 | 188 | 155 | 79 | 84 | 125 | 87 | 214 | 177 |
| PWY0-1415 | superpathway of heme b biosynthesis from uroporphyrinogen-III | contribution | 0.0171524 | 0.018184519 | 0.000804633 | 0.00147359 | 0.004151631 | 0.007613577 | 0.000878787 | 0.001398246 | 0.000181949 | 0.000390363 |
|  |  | rank | 17 | 8 | 152 | 117 | 62 | 33 | 150 | 126 | 230 | 201 |
| PWY0-1533 | methylphosphonate degradation I | contribution | 0.0779301 | 0.040643247 | 0 | 0 | 0.009198302 | 0.008094783 | 0 | 0 | 0 | 0 |
|  |  | rank | 3 | 5 | 40 | 40 | 21 | 19 | 40 | 40 | 40 | 40 |
| PWY490-3 | nitrate reduction VI (assimilatory) | contribution | 0.024028139 | 0.015729783 | 0.000187504 | 0.000729036 | 0.004390424 | 0.004719571 | 0.000883356 | 0.001767954 | 0.00049411 | 0.001112852 |
|  |  | rank | 10 | 17 | 221 | 161 | 51 | 48 | 141 | 104 | 175 | 141 |
| PWY-5022 | 4-aminobutanoate degradation V | contribution | 0 | 0 | 0 | 0 | 0 | 0 | 0 | 0 | 0 | 0 |
|  |  | rank | 89 | 89 | 89 | 89 | 89 | 89 | 89 | 89 | 89 | 89 |
| PWY-5097 | L-lysine biosynthesis VI | contribution | 0.022752592 | 0.020657808 | 0.00042255 | 0.00102518 | 0.00238254 | 0.00223923 | 0.00110982 | 0.002206704 | 0.000493814 | 0.00122084 |
|  |  | rank | 10 | 6 | 191 | 153 | 87 | 94 | 133 | 95 | 183 | 137 |
| PWY-5667 | CDP-diacylglycerol biosynthesis I | contribution | 0.011458406 | 0.010957328 | 0.000440629 | 0.001083098 | 0.002829319 | 0.002774998 | 0.001305202 | 0.002678607 | 0.000279758 | 0.000692743 |
|  |  | rank | 21 | 23 | 188 | 155 | 79 | 84 | 125 | 87 | 214 | 177 |
| PWY-5677 | succinate fermentation to butanoate | contribution | 0 | 0 | 0 | 0 | 0 | 0 | 0 | 0 | 0 | 0 |
|  |  | rank | 88 | 88 | 88 | 88 | 88 | 88 | 88 | 88 | 88 | 88 |
| PWY-5705 | allantoin degradation to glyoxylate III | contribution | 0 | 0 | 0 | 0 | 0 | 0 | 0 | 0 | 0 | 0 |
|  |  | rank | 17 | 17 | 17 | 17 | 17 | 17 | 17 | 17 | 17 | 17 |
| PWY-5845 | superpathway of menaquinol-9 biosynthesis | contribution | 0.084708351 | 0.08229146 | 0 | 0 | 0 | 0 | 0 | 0 | 0.001553714 | 0.002720819 |
|  |  | rank | 3 | 4 | 69 | 69 | 69 | 69 | 69 | 69 | 48 | 37 |
| PWY-5850 | superpathway of menaquinol-6 biosynthesis I | contribution | 0.08472472 | 0.082320479 | 0 | 0 | 0 | 0 | 0 | 0 | 0.001553804 | 0.002721414 |
|  |  | rank | 3 | 4 | 69 | 69 | 69 | 69 | 69 | 69 | 48 | 37 |
| PWY-5860 | superpathway of demethylmenaquinol-6 biosynthesis I | contribution | 0.086121122 | 0.080338709 | 0 | 0 | 0 | 0 | 0 | 0 | 0.00161686 | 0.002554539 |
|  |  | rank | 3 | 4 | 53 | 53 | 53 | 53 | 53 | 53 | 41 | 34 |
| PWY-5862 | superpathway of demethylmenaquinol-9 biosynthesis | contribution | 0.086108983 | 0.080312107 | 0 | 0 | 0 | 0 | 0 | 0 | 0.001616895 | 0.002554091 |
|  |  | rank | 3 | 4 | 53 | 53 | 53 | 53 | 53 | 53 | 41 | 34 |
| **PWY-5896** | **superpathway of menaquinol-10 biosynthesis** | contribution | **0.08472472** | **0.082320479** | 0 | 0 | 0 | 0 | 0 | 0 | 0.001553804 | 0.002721414 |
|  |  | rank | **3** | **4** | 69 | 69 | 69 | 69 | 69 | 69 | 48 | 37 |
| PWY-5913 | partial TCA cycle (obligate autotrophs) | contribution | 0.030814844 | 0.029522449 | 0.000210326 | 0.000549783 | 0.002104563 | 0.002130734 | 0.001149773 | 0.002500275 | 0.000923726 | 0.002014475 |
|  |  | rank | 4 | 6 | 222 | 171 | 88 | 83 | 121 | 73 | 133 | 88 |
| PWY-5918 | superpathway of heme b biosynthesis from glutamate | contribution | 0.017067117 | 0.014264791 | 0.000320477 | 0.000890349 | 0.002110291 | 0.003268183 | 0.000596403 | 0.000949371 | 0.000176158 | 0.000526401 |
|  |  | rank | 14 | 16 | 192 | 145 | 90 | 70 | 163 | 138 | 218 | 175 |
| PWY-5971 | palmitate biosynthesis II (bacteria and plant cytoplasm) | contribution | 0.041258355 | 0.026649031 | 0.000246285 | 0.000971162 | 0.00165064 | 0.002012301 | 0.000777945 | 0.001596763 | 0 | 0 |
|  |  | rank | 3 | 4 | 213 | 147 | 104 | 94 | 146 | 113 | 251 | 251 |
| PWY-6151 | S-adenosyl-L-methionine cycle I | contribution | 0.019761316 | 0.01928943 | 0.000445736 | 0.001117012 | 0.00302796 | 0.003004274 | 0.00058247 | 0.001123652 | 0.000312522 | 0.000814187 |
|  |  | rank | 13 | 9 | 172 | 143 | 72 | 73 | 159 | 142 | 196 | 154 |
| PWY-6471 | peptidoglycan biosynthesis IV (Enterococcus faecium) | contribution | 0.012421978 | 0.010614833 | 0.000282308 | 0.000864491 | 0.002659456 | 0.002951477 | 0.001232161 | 0.003083627 | 0.000266428 | 0.00074787 |
|  |  | rank | 20 | 23 | 215 | 162 | 81 | 79 | 123 | 70 | 217 | 174 |
| PWY-6478 | GDP-D-glycero-alpha-D-manno-heptose biosynthesis | contribution | 0 | 0 | 0 | 0 | 0.011623692 | 0.009280731 | 0 | 0 | 0 | 0 |
|  |  | rank | 58 | 58 | 58 | 58 | 18 | 27 | 58 | 58 | 58 | 58 |
| PWY-6545 | pyrimidine deoxyribonucleotides de novo biosynthesis III | contribution | 0.031940459 | 0.026808733 | 0.000343489 | 0.000772179 | 0.004698932 | 0.004382307 | 0.001347113 | 0.00260641 | 0.000716471 | 0.001707064 |
|  |  | rank | 7 | 6 | 189 | 146 | 48 | 54 | 110 | 88 | 146 | 109 |
| PWY-6588 | pyruvate fermentation to acetone | contribution | 0.009000456 | 0.009672658 | 0.000144371 | 0.000395227 | 0.004336621 | 0.003633529 | 0.000694997 | 0.001347689 | 0.000271926 | 0.000587132 |
|  |  | rank | 24 | 26 | 229 | 193 | 52 | 65 | 167 | 119 | 214 | 174 |
| PWY-6590 | superpathway of Clostridium acetobutylicum acidogenic fermentation | contribution | 0.02856611 | 0.03006335 | 0.000208195 | 0.000512888 | 0.002768296 | 0.002863341 | 0.000558826 | 0.00080756 | 0.000454907 | 0.001331315 |
|  |  | rank | 6 | 4 | 225 | 187 | 74 | 74 | 175 | 155 | 185 | 120 |
| PWY-6749 | CMP-legionaminate biosynthesis I | contribution | 0.009382432 | 0.006354398 | 0.000171413 | 0.000569323 | 0.002485165 | 0.002807945 | 0.000976584 | 0.002585892 | 0.000275953 | 0.000565227 |
|  |  | rank | 23 | 37 | 222 | 177 | 79 | 75 | 133 | 82 | 210 | 179 |
| PWY-6891 | thiazole biosynthesis II (aerobic bacteria) | contribution | 0.015583075 | 0.026018422 | 0.000600339 | 0.001190493 | 0.004955714 | 0.00448513 | 0.00086547 | 0.001655042 | 0.00028798 | 0.000626296 |
|  |  | rank | 17 | 8 | 168 | 137 | 52 | 59 | 147 | 113 | 206 | 172 |
| PWY-6895 | superpathway of thiamine diphosphate biosynthesis II | contribution | 0.023970193 | 0.030930387 | 0.000347518 | 0.000808847 | 0.00272314 | 0.002990505 | 0.000891354 | 0.001793605 | 0.000553948 | 0.001119188 |
|  |  | rank | 7 | 3 | 194 | 158 | 82 | 79 | 144 | 108 | 174 | 144 |
| PWY-7003 | glycerol degradation to butanol | contribution | 0.010025988 | 0.02645448 | 0.000592627 | 0.0012821 | 0.012282452 | 0.011130307 | 0.00119319 | 0.001959244 | 0 | 0 |
|  |  | rank | 28 | 6 | 169 | 131 | 22 | 27 | 124 | 97 | 252 | 252 |
| PWY-7013 | (S)-propane-1,2-diol degradation | contribution | 0.013534491 | 0.01887056 | 0.000564382 | 0.000983825 | 0.005954329 | 0.006515241 | 0.001305666 | 0.001765226 | 0 | 0 |
|  |  | rank | 17 | 8 | 159 | 133 | 38 | 39 | 115 | 101 | 221 | 221 |
| PWY-7377 | cob(II)yrinate a,c-diamide biosynthesis I (early cobalt insertion) | contribution | 0.012229597 | 0.010951351 | 0.000467085 | 0.001217542 | 0.002232903 | 0.002489223 | 0.001303189 | 0.00232439 | 3.90E-05 | 5.70E-05 |
|  |  | rank | 21 | 27 | 163 | 129 | 90 | 87 | 120 | 94 | 228 | 226 |
| REDCITCYC | TCA cycle VI (Helicobacter) | contribution | 0.041416436 | 0.035287776 | 0.000120724 | 0.000319356 | 0.00419306 | 0.003583641 | 0.001031 | 0.001169719 | 0.000446942 | 0.002136079 |
|  |  | rank | 5 | 5 | 232 | 196 | 54 | 58 | 129 | 125 | 173 | 84 |
| TEICHOICACID-PWY | poly(glycerol phosphate) wall teichoic acid biosynthesis | contribution | 0.031911929 | 0.024916938 | 0.000190498 | 0.000516154 | 0.001918634 | 0.001666864 | 0.0022493 | 0.004078157 | 0.001662682 | 0.003411148 |
|  |  | rank | 4 | 6 | 172 | 140 | 78 | 93 | 70 | 49 | 84 | 58 |

**(Supplementary Table 13 continued)**

| **ASV ID** |  |  | **fd496fd32dc8c08ade2e8b6c9d8ee13d** | | **aff3962bbd7607a2971161d9840ca409** | | **a55a010c9525ce2943a553dca1421b1c** | | **2de0f958e30d26f04cba8d2a920bbe46** | | **6ec1023583647a401269990c4e8ad716** | |
| --- | --- | --- | --- | --- | --- | --- | --- | --- | --- | --- | --- | --- |
| **Function** | **Annotation** |  | **Adenoma** | **Cancer** | **Adenoma** | **Cancer** | **Adenoma** | **Cancer** | **Adenoma** | **Cancer** | **Adenoma** | **Cancer** |
| ALL-CHORISMATE-PWY | superpathway of chorismate metabolism | contribution^†^ | 0.019772827 | 0.008926234 | 0.000310891 | 0.000715668 | 0.000917394 | 0.000373338 | 0.000887756 | 0.00077825 | 0.001366371 | 0.000721864 |
|  |  | rank^#^ | 11 | 30 | 193 | 166 | 144 | 190 | 145 | 157 | 117 | 165 |
| CENTFERM-PWY | pyruvate fermentation to butanoate | contribution | 0.006138843 | 0.002121729 | 0.00051031 | 0.000678796 | 0.000863126 | 0.000396481 | 0.001111421 | 0.000875872 | 0.001625936 | 0.000755939 |
|  |  | rank | 42 | 87 | 172 | 157 | 142 | 189 | 123 | 139 | 104 | 150 |
| FAO-PWY | fatty acid beta-oxidation I (generic) | contribution | 0 | 0 | 0.000399296 | 0.000476269 | 0 | 0 | 0.001234266 | 0.000747367 | 0 | 0 |
|  |  | rank | 165 | 165 | 139 | 129 | 165 | 165 | 96 | 111 | 165 | 165 |
| GLUCARDEG-PWY | D-glucarate degradation I | contribution | 0 | 0 | 0 | 0 | 0 | 0 | 0 | 0 | 0 | 0 |
|  |  | rank | 46 | 46 | 46 | 46 | 46 | 46 | 46 | 46 | 46 | 46 |
| HEME-BIOSYNTHESIS-II | heme b biosynthesis I (aerobic) | contribution | 0 | 0 | 0 | 0 | 0 | 0 | 0.002605448 | 0.001923597 | 0 | 0 |
|  |  | rank | 99 | 99 | 99 | 99 | 99 | 99 | 51 | 60 | 99 | 99 |
| HEMESYN2-PWY | heme b biosynthesis II (oxygen-independent) | contribution | 0.009774039 | 0.006612078 | 0.000369086 | 0.00065152 | 0.000776716 | 0.000378775 | 0.001429559 | 0.001276372 | 0.001698136 | 0.00083883 |
|  |  | rank | 25 | 43 | 195 | 172 | 152 | 202 | 115 | 135 | 106 | 160 |
| HISDEG-PWY | L-histidine degradation I | contribution | 0 | 0 | 0 | 0 | 0 | 0 | 0 | 0 | 0 | 0 |
|  |  | rank | 38 | 38 | 38 | 38 | 38 | 38 | 38 | 38 | 38 | 38 |
| P162-PWY | L-glutamate degradation V (via hydroxyglutarate) | contribution | 0.010139224 | 0.014430157 | 0.001164444 | 0.000916904 | 0 | 0 | 0.00079615 | 0.000411171 | 0.000817696 | 0.000413597 |
|  |  | rank | 23 | 14 | 95 | 90 | 134 | 134 | 106 | 116 | 104 | 115 |
| P163-PWY | L-lysine fermentation to acetate and butanoate | contribution | 0.009671871 | 0.014069866 | 0.00050001 | 0.000834545 | 0.000381634 | 0.000205524 | 0.00090087 | 0.000690339 | 0.000715392 | 0.000555323 |
|  |  | rank | 27 | 16 | 191 | 161 | 201 | 230 | 152 | 173 | 169 | 184 |
| P164-PWY | purine nucleobases degradation I (anaerobic) | contribution | 0.01077699 | 0.005222552 | 0.000217909 | 0.000282454 | 0.000369945 | 0.000234667 | 0.000878458 | 0.00080921 | 0.00073822 | 0.000494662 |
|  |  | rank | 22 | 47 | 215 | 201 | 186 | 210 | 137 | 149 | 151 | 181 |
| PWY0-1319 | CDP-diacylglycerol biosynthesis II | contribution | 0.015015734 | 0.009005968 | 0.000363432 | 0.000453029 | 0.000629907 | 0.000422014 | 0.00108802 | 0.001331912 | 0.001168521 | 0.000887171 |
|  |  | rank | 17 | 31 | 200 | 199 | 170 | 202 | 140 | 135 | 135 | 163 |
| PWY0-1415 | superpathway of heme b biosynthesis from uroporphyrinogen-III | contribution | 0.01051992 | 0.004513698 | 0.000383588 | 0.000588617 | 0.000947798 | 0.000555483 | 0.001407277 | 0.001206785 | 0.001686787 | 0.000758595 |
|  |  | rank | 24 | 58 | 191 | 182 | 146 | 185 | 121 | 135 | 102 | 164 |
| PWY0-1533 | methylphosphonate degradation I | contribution | 0 | 0 | 0 | 0 | 0 | 0 | 0 | 0 | 0 | 0 |
|  |  | rank | 40 | 40 | 40 | 40 | 40 | 40 | 40 | 40 | 40 | 40 |
| PWY490-3 | nitrate reduction VI (assimilatory) | contribution | 0.008697133 | 0.008279771 | 0.000408321 | 0.000327231 | 0.000589943 | 0.000414946 | 0.001100702 | 0.001366493 | 0.001146784 | 0.00082849 |
|  |  | rank | 31 | 34 | 182 | 204 | 164 | 195 | 122 | 129 | 119 | 157 |
| PWY-5022 | 4-aminobutanoate degradation V | contribution | 0 | 0 | 0.001421753 | 0.001655528 | 0 | 0 | 0 | 0 | 0 | 0 |
|  |  | rank | 89 | 89 | 68 | 62 | 89 | 89 | 89 | 89 | 89 | 89 |
| PWY-5097 | L-lysine biosynthesis VI | contribution | 0.009455405 | 0.005501438 | 0.000354588 | 0.000437143 | 0.000614806 | 0.000403408 | 0.001347444 | 0.001593298 | 0.001160739 | 0.000852054 |
|  |  | rank | 29 | 43 | 202 | 195 | 173 | 198 | 116 | 121 | 130 | 168 |
| PWY-5667 | CDP-diacylglycerol biosynthesis I | contribution | 0.015015734 | 0.009005968 | 0.000363432 | 0.000453029 | 0.000629907 | 0.000422014 | 0.00108802 | 0.001331912 | 0.001168521 | 0.000887171 |
|  |  | rank | 17 | 31 | 200 | 199 | 170 | 202 | 140 | 135 | 135 | 163 |
| PWY-5677 | succinate fermentation to butanoate | contribution | 0 | 0 | 0.001491179 | 0.002414371 | 0 | 0 | 0 | 0 | 0 | 0 |
|  |  | rank | 88 | 88 | 69 | 56 | 88 | 88 | 88 | 88 | 88 | 88 |
| PWY-5705 | allantoin degradation to glyoxylate III | contribution | 0 | 0 | 0 | 0 | 0 | 0 | 0 | 0 | 0 | 0 |
|  |  | rank | 17 | 17 | 17 | 17 | 17 | 17 | 17 | 17 | 17 | 17 |
| PWY-5845 | superpathway of menaquinol-9 biosynthesis | contribution | 0 | 0 | 0 | 0 | 0 | 0 | 0 | 0 | 0 | 0 |
|  |  | rank | 69 | 69 | 69 | 69 | 69 | 69 | 69 | 69 | 69 | 69 |
| PWY-5850 | superpathway of menaquinol-6 biosynthesis I | contribution | 0 | 0 | 0 | 0 | 0 | 0 | 0 | 0 | 0 | 0 |
|  |  | rank | 69 | 69 | 69 | 69 | 69 | 69 | 69 | 69 | 69 | 69 |
| PWY-5860 | superpathway of demethylmenaquinol-6 biosynthesis I | contribution | 0 | 0 | 0 | 0 | 0 | 0 | 0 | 0 | 0 | 0 |
|  |  | rank | 53 | 53 | 53 | 53 | 53 | 53 | 53 | 53 | 53 | 53 |
| PWY-5862 | superpathway of demethylmenaquinol-9 biosynthesis | contribution | 0 | 0 | 0 | 0 | 0 | 0 | 0 | 0 | 0 | 0 |
|  |  | rank | 53 | 53 | 53 | 53 | 53 | 53 | 53 | 53 | 53 | 53 |
| **PWY-5896** | **superpathway of menaquinol-10 biosynthesis** | contribution | 0 | 0 | 0 | 0 | 0 | 0 | 0 | 0 | 0 | 0 |
|  |  | rank | 69 | 69 | 69 | 69 | 69 | 69 | 69 | 69 | 69 | 69 |
| PWY-5913 | partial TCA cycle (obligate autotrophs) | contribution | 0.010881797 | 0.004982946 | 0.000273458 | 0.000351027 | 0.000528602 | 0.000297668 | 0.000684684 | 0.000744866 | 0.001033616 | 0.00067988 |
|  |  | rank | 27 | 48 | 204 | 200 | 169 | 205 | 162 | 161 | 127 | 165 |
| PWY-5918 | superpathway of heme b biosynthesis from glutamate | contribution | 0 | 0 | 0.000110098 | 0.000172049 | 0.001577222 | 0.000813001 | 0.002358782 | 0.001962257 | 0.003316756 | 0.001592479 |
|  |  | rank | 247 | 247 | 229 | 224 | 106 | 151 | 84 | 99 | 59 | 112 |
| PWY-5971 | palmitate biosynthesis II (bacteria and plant cytoplasm) | contribution | 0.011366614 | 0.007297557 | 0.00026261 | 0.000270112 | 0.000737522 | 0.00054485 | 0.001418772 | 0.001515789 | 0.001469044 | 0.000755997 |
|  |  | rank | 24 | 40 | 207 | 208 | 150 | 180 | 113 | 119 | 111 | 161 |
| PWY-6151 | S-adenosyl-L-methionine cycle I | contribution | 0.021343896 | 0.012163508 | 0.0003704 | 0.000410588 | 0.000677304 | 0.000463712 | 0.00097775 | 0.001233469 | 0.001270544 | 0.001008723 |
|  |  | rank | 12 | 20 | 185 | 191 | 154 | 183 | 128 | 134 | 116 | 145 |
| PWY-6471 | peptidoglycan biosynthesis IV (Enterococcus faecium) | contribution | 0.017576751 | 0.010103461 | 0.000430963 | 0.000437999 | 0.000765643 | 0.00052919 | 0.001231263 | 0.001634593 | 0.001579042 | 0.001156235 |
|  |  | rank | 15 | 27 | 185 | 202 | 158 | 188 | 124 | 125 | 111 | 145 |
| PWY-6478 | GDP-D-glycero-alpha-D-manno-heptose biosynthesis | contribution | 0 | 0 | 0.001453715 | 0.001061533 | 0 | 0 | 0 | 0 | 0.004873372 | 0.003702599 |
|  |  | rank | 58 | 58 | 51 | 55 | 58 | 58 | 58 | 58 | 38 | 45 |
| PWY-6545 | pyrimidine deoxyribonucleotides de novo biosynthesis III | contribution | 0.040309746 | 0.018467189 | 0.000275907 | 0.00038298 | 0.00032707 | 0.000209916 | 0.001195449 | 0.001356894 | 0.000303375 | 0.00023238 |
|  |  | rank | 3 | 14 | 198 | 190 | 192 | 214 | 122 | 117 | 194 | 211 |
| PWY-6588 | pyruvate fermentation to acetone | contribution | 0.006669261 | 0.004684035 | 0.000421682 | 0.000496892 | 0.000661008 | 0.000386702 | 0.000948919 | 0.001009014 | 0.001157269 | 0.000806955 |
|  |  | rank | 37 | 52 | 187 | 182 | 169 | 197 | 140 | 140 | 124 | 155 |
| PWY-6590 | superpathway of Clostridium acetobutylicum acidogenic fermentation | contribution | 0.009692445 | 0.003820217 | 0.000460932 | 0.000678859 | 0.000921959 | 0.000440247 | 0.001194762 | 0.000984738 | 0.001758992 | 0.000870756 |
|  |  | rank | 31 | 59 | 184 | 173 | 147 | 194 | 127 | 145 | 101 | 151 |
| PWY-6749 | CMP-legionaminate biosynthesis I | contribution | 0.011843591 | 0.007153448 | 0.000344406 | 0.000322046 | 0.000762831 | 0.000524644 | 0.000982988 | 0.001273529 | 0.001304292 | 0.00100818 |
|  |  | rank | 19 | 34 | 193 | 207 | 149 | 183 | 132 | 126 | 119 | 145 |
| PWY-6891 | thiazole biosynthesis II (aerobic bacteria) | contribution | 0.014944466 | 0.010431513 | 0.000363612 | 0.000569152 | 0.00068956 | 0.000381752 | 0.001132893 | 0.001000422 | 0.001238448 | 0.000818047 |
|  |  | rank | 19 | 28 | 191 | 179 | 160 | 198 | 132 | 145 | 126 | 156 |
| PWY-6895 | superpathway of thiamine diphosphate biosynthesis II | contribution | 0.006103996 | 0.003576497 | 0.00038992 | 0.000562101 | 0.000703457 | 0.000434136 | 0.001220257 | 0.001152569 | 0.001557496 | 0.001013707 |
|  |  | rank | 44 | 68 | 187 | 180 | 165 | 185 | 126 | 140 | 110 | 146 |
| PWY-7003 | glycerol degradation to butanol | contribution | 0.02276839 | 0.010417008 | 0.00042301 | 0.000306087 | 0.000512544 | 0.000274517 | 0.000924573 | 0.000717027 | 0.001077397 | 0.000757945 |
|  |  | rank | 10 | 29 | 187 | 205 | 178 | 215 | 144 | 162 | 131 | 160 |
| PWY-7013 | (S)-propane-1,2-diol degradation | contribution | 0.034338362 | 0.015491117 | 0.00037251 | 0.000439773 | 0.001164519 | 0.000365306 | 0.001083033 | 0.001433122 | 0.000751237 | 0.00040179 |
|  |  | rank | 4 | 13 | 175 | 171 | 123 | 178 | 130 | 114 | 145 | 175 |
| PWY-7377 | cob(II)yrinate a,c-diamide biosynthesis I (early cobalt insertion) | contribution | 0 | 0 | 8.68E-05 | 0.000102118 | 0.000923212 | 0.00057225 | 0.000960444 | 0.001241486 | 0.001488584 | 0.0008573 |
|  |  | rank | 235 | 235 | 216 | 214 | 137 | 165 | 134 | 125 | 112 | 148 |
| REDCITCYC | TCA cycle VI (Helicobacter) | contribution | 0.00962084 | 0.004606268 | 0.000312227 | 0.000421133 | 0.000467313 | 0.000260659 | 0.00099882 | 0.000710465 | 0.00167071 | 0.000784534 |
|  |  | rank | 30 | 49 | 189 | 184 | 171 | 203 | 133 | 158 | 100 | 150 |
| TEICHOICACID-PWY | poly(glycerol phosphate) wall teichoic acid biosynthesis | contribution | 0.011144176 | 0.006309731 | 0.000204284 | 0.000173608 | 0.000538821 | 0.000420497 | 0.001116924 | 0.001377671 | 0 | 0 |
|  |  | rank | 18 | 33 | 169 | 174 | 136 | 149 | 105 | 99 | 190 | 190 |

**(Supplementary Table 13 continued)**

| **ASV ID** |  |  | **e9f900358bb9297e10b9fbb1321b1e6e** | | **f50508546ae13143015f8c4cb976d0e4** | |
| --- | --- | --- | --- | --- | --- | --- |
| **Function** | **Annotation** |  | **Adenoma** | **Cancer** | **Adenoma** | **Cancer** |
| ALL-CHORISMATE-PWY | superpathway of chorismate metabolism | contribution^†^ | 0.000464042 | 0.000167846 | **Adenoma** | **Cancer** |
|  |  | rank^#^ | 175 | 222 | 0.004587823 | 0.000998642 |
| CENTFERM-PWY | pyruvate fermentation to butanoate | contribution | 0.000157198 | 4.41E-05 | 53 | 143 |
|  |  | rank | 224 | 242 | 0.010058555 | 0.003075714 |
| FAO-PWY | fatty acid beta-oxidation I (generic) | contribution | 0 | 0 | 31 | 70 |
|  |  | rank | 165 | 165 | 0.01698784 | 0.005726924 |
| GLUCARDEG-PWY | D-glucarate degradation I | contribution | 0.002843525 | 0.000483209 | 15 | 31 |
|  |  | rank | 31 | 43 | 0 | 0 |
| HEME-BIOSYNTHESIS-II | heme b biosynthesis I (aerobic) | contribution | 0 | 0 | 46 | 46 |
|  |  | rank | 99 | 99 | 0.045709644 | 0.01568788 |
| HEMESYN2-PWY | heme b biosynthesis II (oxygen-independent) | contribution | 0.000253698 | 5.95E-05 | 6 | 18 |
|  |  | rank | 210 | 247 | 0.013109599 | 0.006674826 |
| HISDEG-PWY | L-histidine degradation I | contribution | 0 | 0 | 22 | 42 |
|  |  | rank | 38 | 38 | 0 | 0 |
| P162-PWY | L-glutamate degradation V (via hydroxyglutarate) | contribution | 0 | 0 | 38 | 38 |
|  |  | rank | 134 | 134 | 0.004398685 | 0.001188673 |
| P163-PWY | L-lysine fermentation to acetate and butanoate | contribution | 0.000891049 | 0.000221473 | 48 | 80 |
|  |  | rank | 154 | 223 | 0.005685261 | 0.002884907 |
| P164-PWY | purine nucleobases degradation I (anaerobic) | contribution | 0.00070811 | 0.000250318 | 48 | 75 |
|  |  | rank | 153 | 206 | 0.00386935 | 0.00212304 |
| PWY0-1319 | CDP-diacylglycerol biosynthesis II | contribution | 0.000733168 | 0.000288161 | 59 | 82 |
|  |  | rank | 165 | 223 | 0.005897194 | 0.0031064 |
| PWY0-1415 | superpathway of heme b biosynthesis from uroporphyrinogen-III | contribution | 0.000193471 | 5.78E-05 | 44 | 73 |
|  |  | rank | 229 | 248 | 0.018068488 | 0.006625025 |
| PWY0-1533 | methylphosphonate degradation I | contribution | 0 | 0 | 16 | 40 |
|  |  | rank | 40 | 40 | 0 | 0 |
| PWY490-3 | nitrate reduction VI (assimilatory) | contribution | 0.000377358 | 0.000184427 | 40 | 40 |
|  |  | rank | 189 | 226 | 0.005864663 | 0.002326717 |
| PWY-5022 | 4-aminobutanoate degradation V | contribution | 0 | 0 | 38 | 86 |
|  |  | rank | 89 | 89 | 0 | 0 |
| PWY-5097 | L-lysine biosynthesis VI | contribution | 0.000807334 | 0.000310047 | 89 | 89 |
|  |  | rank | 157 | 215 | 0.006012275 | 0.003141047 |
| PWY-5667 | CDP-diacylglycerol biosynthesis I | contribution | 0.000733168 | 0.000288161 | 42 | 70 |
|  |  | rank | 165 | 223 | 0.005897194 | 0.0031064 |
| PWY-5677 | succinate fermentation to butanoate | contribution | 0 | 0 | 44 | 73 |
|  |  | rank | 88 | 88 | 0 | 0 |
| PWY-5705 | allantoin degradation to glyoxylate III | contribution | 0 | 0 | 88 | 88 |
|  |  | rank | 17 | 17 | 0 | 0 |
| PWY-5845 | superpathway of menaquinol-9 biosynthesis | contribution | 0 | 0 | 17 | 17 |
|  |  | rank | 69 | 69 | 0 | 0 |
| PWY-5850 | superpathway of menaquinol-6 biosynthesis I | contribution | 0 | 0 | 69 | 69 |
|  |  | rank | 69 | 69 | 0 | 0 |
| PWY-5860 | superpathway of demethylmenaquinol-6 biosynthesis I | contribution | 0 | 0 | 69 | 69 |
|  |  | rank | 53 | 53 | 0 | 0 |
| PWY-5862 | superpathway of demethylmenaquinol-9 biosynthesis | contribution | 0 | 0 | 53 | 53 |
|  |  | rank | 53 | 53 | 0 | 0 |
| **PWY-5896** | **superpathway of menaquinol-10 biosynthesis** | contribution | 0 | 0 | 53 | 53 |
|  |  | rank | 69 | 69 | 0 | 0 |
| PWY-5913 | partial TCA cycle (obligate autotrophs) | contribution | 0.000460172 | 0.000128474 | 69 | 69 |
|  |  | rank | 175 | 236 | 0.003750609 | 0.002122307 |
| PWY-5918 | superpathway of heme b biosynthesis from glutamate | contribution | 0.000116379 | 3.10E-05 | 60 | 84 |
|  |  | rank | 228 | 243 | 0.011979653 | 0.004883075 |
| PWY-5971 | palmitate biosynthesis II (bacteria and plant cytoplasm) | contribution | 0.000337712 | 0.000117472 | 21 | 49 |
|  |  | rank | 196 | 235 | 0.007639982 | 0.002875348 |
| PWY-6151 | S-adenosyl-L-methionine cycle I | contribution | 0.001282843 | 0.000521877 | 34 | 70 |
|  |  | rank | 115 | 179 | 0.005297886 | 0.002744527 |
| PWY-6471 | peptidoglycan biosynthesis IV (Enterococcus faecium) | contribution | 0.000753178 | 0.000338762 | 44 | 78 |
|  |  | rank | 159 | 216 | 0.005436428 | 0.002735733 |
| PWY-6478 | GDP-D-glycero-alpha-D-manno-heptose biosynthesis | contribution | 0 | 0 | 44 | 84 |
|  |  | rank | 58 | 58 | 0 | 0 |
| PWY-6545 | pyrimidine deoxyribonucleotides de novo biosynthesis III | contribution | 0.001058766 | 0.000339772 | 58 | 58 |
|  |  | rank | 127 | 199 | 0.001315264 | 0.000662102 |
| PWY-6588 | pyruvate fermentation to acetone | contribution | 0.0003613 | 0.000139436 | 114 | 161 |
|  |  | rank | 197 | 235 | 0.008047113 | 0.004385893 |
| PWY-6590 | superpathway of Clostridium acetobutylicum acidogenic fermentation | contribution | 0.000343679 | 9.74E-05 | 29 | 54 |
|  |  | rank | 206 | 241 | 0.010695831 | 0.003551367 |
| PWY-6749 | CMP-legionaminate biosynthesis I | contribution | 0.000578212 | 0.000230709 | 25 | 63 |
|  |  | rank | 163 | 223 | 0.005781938 | 0.002403972 |
| PWY-6891 | thiazole biosynthesis II (aerobic bacteria) | contribution | 0.000558585 | 0.000166547 | 42 | 87 |
|  |  | rank | 171 | 235 | 0.003750202 | 0.001902919 |
| PWY-6895 | superpathway of thiamine diphosphate biosynthesis II | contribution | 0.000519331 | 0.000157548 | 62 | 103 |
|  |  | rank | 176 | 234 | 0.004725693 | 0.002411189 |
| PWY-7003 | glycerol degradation to butanol | contribution | 0.000758824 | 0.000387125 | 56 | 91 |
|  |  | rank | 157 | 191 | 0.004146886 | 0.001925673 |
| PWY-7013 | (S)-propane-1,2-diol degradation | contribution | 0.001164616 | 0.00029239 | 58 | 99 |
|  |  | rank | 122 | 185 | 0 | 0 |
| PWY-7377 | cob(II)yrinate a,c-diamide biosynthesis I (early cobalt insertion) | contribution | 8.02E-05 | 2.70E-05 | 221 | 221 |
|  |  | rank | 217 | 230 | 0.0051139 | 0.003601112 |
| REDCITCYC | TCA cycle VI (Helicobacter) | contribution | 0.000204618 | 4.26E-05 | 47 | 66 |
|  |  | rank | 212 | 246 | 0.005105802 | 0.002294746 |
| TEICHOICACID-PWY | poly(glycerol phosphate) wall teichoic acid biosynthesis | contribution | 0.002383495 | 0.001099614 | 45 | 78 |
|  |  | rank | 67 | 107 | 0.001964665 | 0.00113073 |

**§ The ASV ID asigned as Bacteroides.**

**† Defined as the proportion of a per ASV function abundance compared to the total function abundance of all ASVs corresponding to a given pathway.**

**# The rank of the contribution in control and adenoma.**

**Supplementary Table 14 | Characteristics of human samples for qRT-PCR**

|  | **Control (n=7)** | **Adenoma (n=6)** | **CRC(n=30)** |
| --- | --- | --- | --- |
| **Sex (F/M)** | 4/3 | 2/4 | 12/8 |
| **Age (years)** | 48-71 | 50-70 | 41-77 |
| **BMI(average±s.d.)^#^** | 24.07±0.65 | 21.95±2.2.0 | 22.74±1.95 |

**# standard deviation**

**Supplementary Table 15 | Primers for candidate genes**

| **ID** | **Primer name** | **Primer sequence (5'to3'）** |
| --- | --- | --- |
| 1 | fNL303-forward GmhA | TCTCCCGCTATGTTGAAGCG |
| 2 | rNL304-reverse GmhA | TCAATATCCGCCGTACCAGC |
| 3 | fNL305-forward hIdE | TCGTCGTATGGCGGTATTGG |
| 4 | rNL306-reverse hIdE | GCAGCAATCACTTCAGCACC |
| 5 | fNL307-forward gmhB | ACATCCGGGGATGCTTTTGT |
| 6 | rNL308-reverse gmhB | CCAGCACTTTTGTTCCCACG |
| 7 | fNL311-forward rfaD | AGCGTCGCTTTCCATCTCAA |
| 8 | rNL312-reverse rfaD | CCGAGATTGAAGATGCCGGA |
| 9 | fNL313-forward menC | GGCGGTGATCAGTTCTTCCA |
| 10 | rNL314-reverse menC | CATCAGATCCAGCGTGTCCA |
| 11 | fNL317-forward menH | GTTGATCTCCCAGGTCACGG |
| 12 | rNL318-reverse menH | TGTTCAGCATTTTGCAGCCC |
| 13 | fNL319-forward menF | ATCTTCGCCGCTGTATCTGG |
| 14 | rNL320-reverse menF | AATTTTTGCTGAGCGCAGGG |
